# H+ ions and ATP reshape the conformational landscape of an RNA recognition motif and regulate its fibrillation

**DOI:** 10.1101/2025.09.02.673685

**Authors:** Osama Aazmi, Akshit Rajendra Aswale, Jeetender Chugh

## Abstract

Proteins exist as dynamic ensembles, with their native states comprising interconverting conformational substates critical to their physiological functions and participation in disease states. Fused in Sarcoma (FUS), an RNA-binding protein implicated in neurodegenerative diseases such as Amyotrophic Lateral Sclerosis (ALS) and Frontotemporal Dementia (FTD), contains an RNA recognition motif (RRM) known to form fibrillar aggregates. Here, we investigate the conformational plasticity of FUS-RRM in its native state using advanced NMR techniques, particularly ^15^N chemical exchange saturation transfer and heteronuclear adiabatic relaxation dispersion experiments, to capture slow and fast microsecond (μs) timescale dynamics. We further examine the influence of environmental factors such as pH and ATP on the conformational plasticity and the aggregation behaviour of FUS-RRM. Our findings show that both ATP and pH perturb the fast and slow μs-timescale dynamics of FUS-RRM, and the aggregation behaviour. Specifically, a contrasting effect of ATP on slow and fast µs-ms dynamics at pH 6.4 and 4.6, along with the corresponding changes in aggregation behavior, suggest a complex relationship between ATP, pH, and protein aggregation kinetics. The study suggests that these environmental perturbations behave as kinetic regulators of FUS-RRM’s propensity for aggregation.

## Introduction

The energy landscape theory describes how proteins fold and function.^1^ It visualizes different possible conformational states of proteins as points on a multidimensional energy surface, or the “folding funnel”, whose populations are dependent on the different environmental factors. The conformational plasticity is the greatest near the top of the protein folding funnel, where the protein explores a wide range of structural possibilities. As it progresses toward its native state, this flexibility gradually diminishes, giving way to a more defined and energetically stable conformation. The native state of a protein, long regarded as a singular, well-defined structure, is now known to comprise a dynamic ensemble of interconverting substates, each subtly different in structure and potentially in function.^2,3^ However, the nature, extent, and biological implications of this conformational heterogeneity in the native state remain poorly understood. Unravelling this complexity is essential for gaining deeper insights into protein function, regulation, allostery, and the molecular basis of diseases associated with protein misfolding and dysfunction. While structure determination methods such as X-ray crystallography, NMR spectroscopy, cryo-electron microscopy, and predictive tools like AlphaFold primarily focus on the protein’s so-called native state,^4–8^ the multitude of other conformations across the energy landscape are often overlooked due to technical challenges. The sequence-structure-function paradigm,^2,9^ a fundamental concept in molecular biology and biochemistry, describes the relationship between a protein’s amino acid sequence, its three-dimensional structure, and its biological function. However, this paradigm primarily emphasizes the native fold, overlooking the broader conformational landscape that proteins may explore. Advanced experimental techniques like time-resolved Förster resonance energy transfer (FRET), single-molecule force spectroscopy, and molecular dynamics (MD) simulations have been useful in providing valuable insights into stable intermediate states.^10,11^ However, the limited understanding of these conformational substates or excited states stems primarily from their low population, rapid interconversion, and often isostructural nature— factors that make them elusive to traditional structural biology techniques. In this direction, NMR spectroscopy offers a multiprong approach via advanced NMR experiments such as Carr-Purcell-Meiboom-Gill (CPMG) relaxation dispersion experiments,^12,13^ rotating-frame relaxation dispersion experiments (R_1ρ_ and R_2ρ_),^14,15^ chemical exchange via saturation transfer experiments (CEST),^16,17^ dark exchange saturation transfer (DEST),^18,19^ and paramagnetic relaxation enhancement (PRE)^20^ to probe the conformational plasticity of biomolecules across a wide range of timescales (slow ms to fast μs). This unique ability makes NMR spectroscopy indispensable for a deeper understanding of processes such as ligand binding, allosteric regulation, folding, etc., via the lens of conformational plasticity.

It has been shown that conformational dynamics are closely related to protein aggregation and demand a full characterization of processes on a wide range of timescales. For example, conformational dynamics play an important role in the dissociation of the tetramers formed by transthyretin variants, which exposes the β-strands (F and H) for rapid aggregation.^21^ Moreover, simulations have established the role of conformational dynamics in regulating protein stability, highlighting their crucial roles in the process of protein aggregation.^22,23^ However, despite the growing interest in excited states’ functional implications, these have not been explored for many RNA binding proteins. Fused in sarcoma (FUS) is an RNA-binding protein that plays crucial roles in various cellular processes, including gene expression, RNA splicing, and the formation of ribonucleoprotein (RNP) complexes.^24^ Beyond its roles in RNA metabolism, FUS has been implicated in various neurodegenerative diseases, including ALS and FTD.^25^ FUS comprises several functional domains, including low-complexity (LC)-rich regions, RNA recognition motif (RRM), and arginine-glycine-glycine (RGG) motifs. The FUS-RRM exhibits a characteristic fold consisting of two α-helices and four β-strands.^26^ It has been shown that, similar to N- and C-terminal LC-rich regions, FUS-RRM can also form fibrillar aggregates.^27^ Earlier work on FUS-RRM has emphasized the role of picosecond-nanosecond (ps-ns) dynamics in aggregation, which are thought to contribute to high conformational flexibility, implying a low energy barrier between folded and unfolded states.^28^ In a recent study, we showed that the FUS-RRM monomer exhibits conformational heterogeneity at fast μs timescales, which gets perturbed by pH.^29^ We showed that the observed μs-timescale conformational plasticity is coupled to the aggregation kinetics. This prompted a deeper exploration into the extent and nature of structural variability within what is considered a functional “native” state, and how this intrinsic plasticity of FUS-RRM may influence the fibrillation process.

In this study, we have used ^15^N CEST and ^15^N rotating-frame relaxation dispersion experiments (R_1ρ_ and R_2ρ_) to capture slow and fast μs timescale conformational plasticity in the native state of FUS-RRM, respectively, and explored the effect of pH- and Adenosine 5′-triphosphate (ATP)-induced perturbations on the same. We report that in addition to pH, RRM binders including ATP can also modulate the conformational plasticity of FUS-RRM, which in turn influences its aggregation behaviour. The study provides an important insight into the dynamic regulation of FUS-RRM aggregation and its implications for neurodegenerative diseases such as ALS and FTD.

## Results

### FUS-RRM populates multiple excited state conformations

Taking hints from the presence of fast μs-ms timescale dynamics in FUS-RRM,^29^ we utilised the ^15^N-CEST NMR experiment^16^ to identify slower μs-ms dynamics and understand its role in aggregation, if any. The ^15^N-CEST data acquired at B_1_ field of 30 Hz shows two well-separated dips for many isolated residues of FUS-RRM at pH 6.4 (Figure 1A). Increasing or decreasing the B_1_ field leads to broadening of the minor dips, thereby allowing us to only get information from the 30 Hz B_1_ field (Figure S1). The presence of a distinct excited state can be discerned from the minor dip in the ^15^N-CEST profiles of a number of residues (Figures 1A and S2). A total of 16 residues were globally modelled to a two-state exchange process using the Bloch-McConnell equations (Figure S2). The fitting of the CEST data reveals that the population of the excited state (p_ES_) is 0.14±0.01% exchanging with a rate (k_ex_) of 257±36 s^-1^. The excited state conformation is thus calculated to be 3.9 kcal/mol higher in free energy than the ground state (Figure 1B).

**Figure 1.**
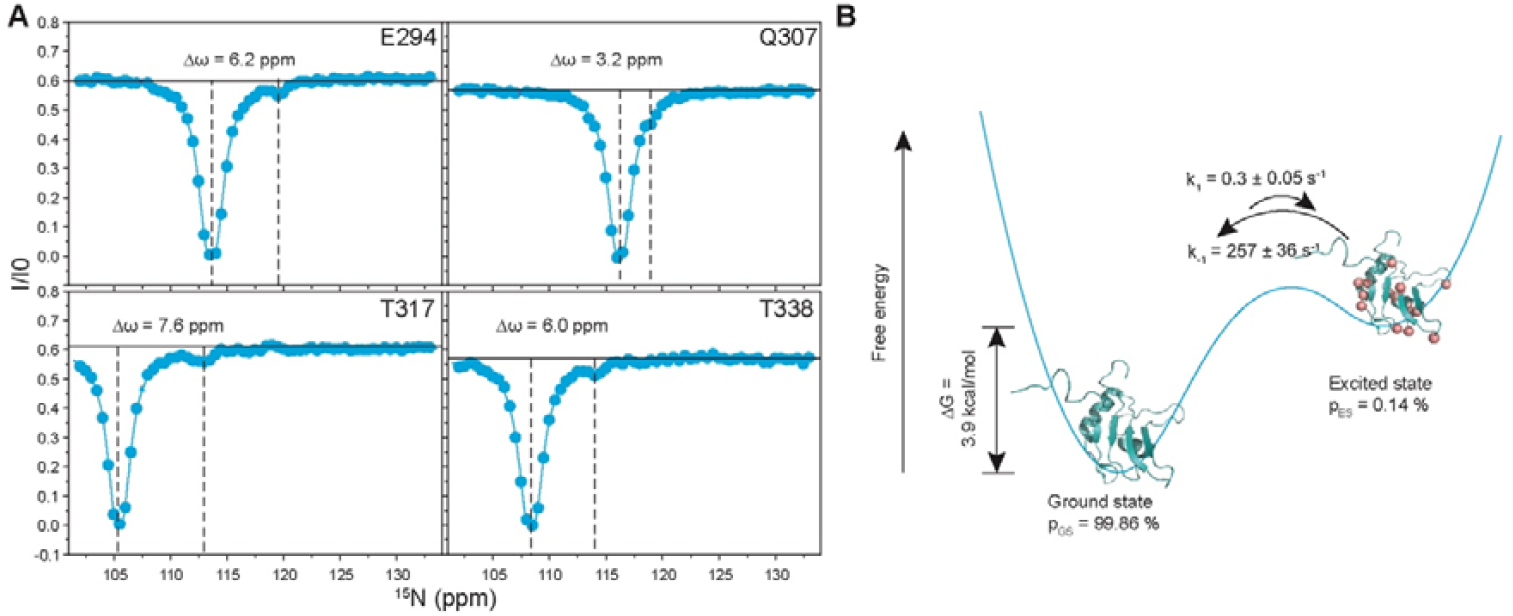
(A) ^15^N-CEST profiles of representative FUS-RRM residues measured at pH 6.4. The vertical dashed lines identify the different states of FUS-RRM and show the chemical shift difference between the ground and excited state. The obtained χ^2^_red_ is 2.3 using fit to a two-state model. (B) Representation of the energy landscape of FUS-RRM showing the two states (ground and excited states) and their interconversion rates with fractional populations.

Since we had earlier observed pH-induced quenching of a fast μs timescale excited state (ES^HARD^) in FUS-RRM (using ^15^N heteronuclear adiabatic relaxation dispersion (HARD) measurements),^29^ we probed the pH-induced perturbation on the excited state detected by the ^15^N-CEST above (referred to as ES^CEST^). Upon perturbing the pH, minor dips were detected for 10 residues in the ^15^N-CEST profiles of FUS-RRM at pH 4.6 (Figure 2A, Orange, and Figure S3). After global fitting in ChemEx, a k_ex_ of 142±36 s^-1^ and p_ES_ of 0.5±0.06% was obtained. Figure 2B shows a cartoon representation of the free energy of interconversion to the excited state. Compared to pH 6.4, ΔG reduces from 3.9 kcal/mol to 3.13 kcal/mol, indicating an ease of accessing the ES^CEST^ at lower pH.

**Figure 2.**
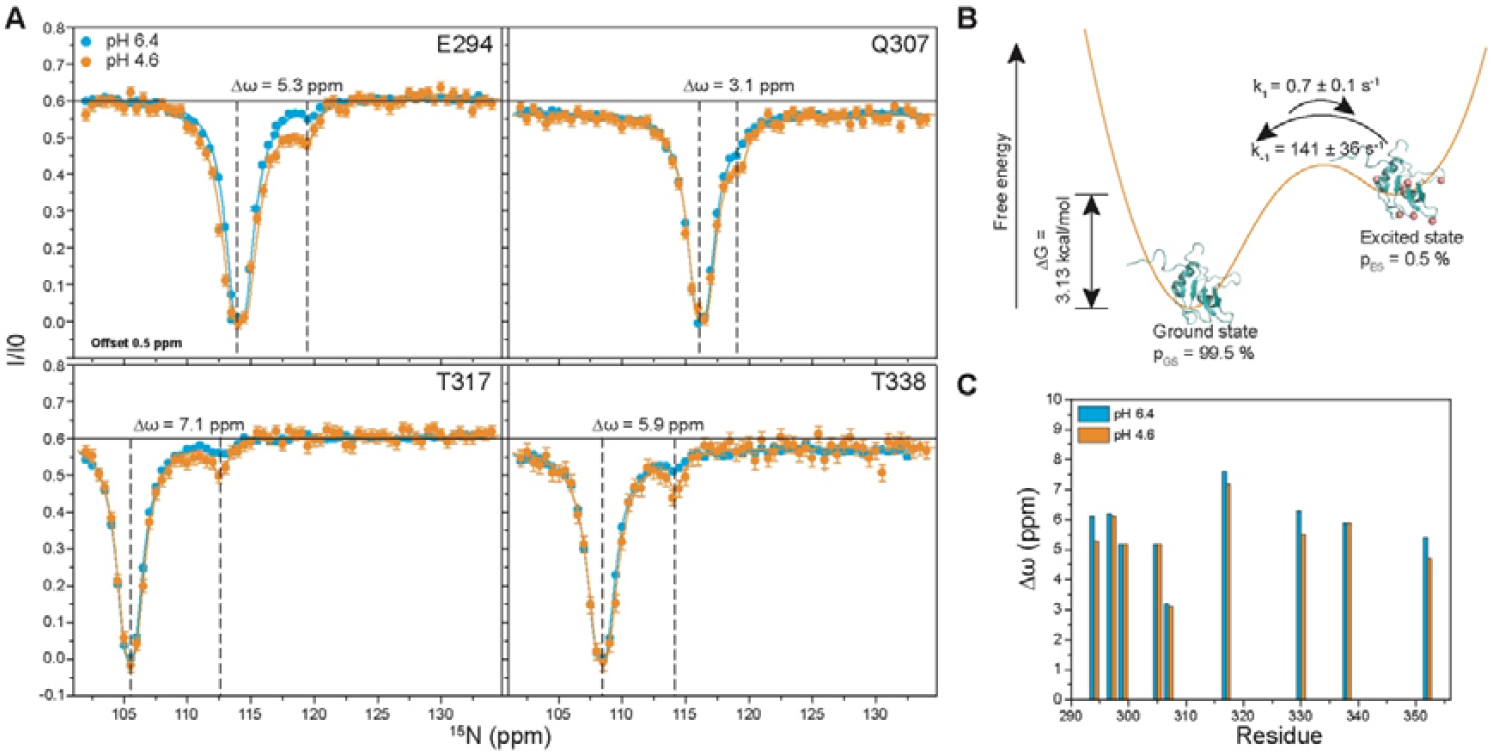
(A) An overlay of ^15^N-CEST profiles of representative FUS-RRM residues measured at the pH 6.4 (cyan) and pH 4.6 (orange). The vertical dashed lines show the chemical shift difference between the ground and the excited state. The obtained χ^2^_red_ is 1.3 using fit to a two-state model at pH 4.6. The vertical dashed lines show large chemical shift changes between the ground and excited state. (B) A cartoon representation of the energy landscape of FUS-RRM showing the two states and their interconversion rates with fractional populations at pH 4.6. (C) Comparison of delta omega (Δω) values obtained from ^15^N-CEST measured at pH 6.4 (cyan) and 4.6 (orange).

Despite a difference in the pH, the same residues show exchange at both pH 6.4 and pH 4.6 with similar Δω values (Figure 2C), thereby indicating that lowering the pH did not quench the conformational dynamics in this timescale regime. On the other hand, reducing the pH further stabilizes the ES^CEST^, suggesting that ES^CEST^ could be an unfolded/partially unfolded state. The residues undergoing exchange have been shown to belong to five clusters (residues N284-I287, T313-T317, L324-G331, E336-V339, and A369-N376), which are located within or near the known interaction interface of FUS-RRM with nucleic acids and/or ATP (Figure 3). This suggests that although the dynamic residues are spatially dispersed, they may collectively contribute to functionally relevant interactions. Notably, residues T317 (loop 2), T330 (loop 3), and T338 (β-3 strand) have been shown to contribute significant roles in nucleic acid binding.^30,31^ The results from the present study when combined with those reported from the earlier studies^28,29^ show that the FUS-RRM samples multiple conformational states, including a native state, ES^HARD^, and ES^CEST^.

**Figure 3:**
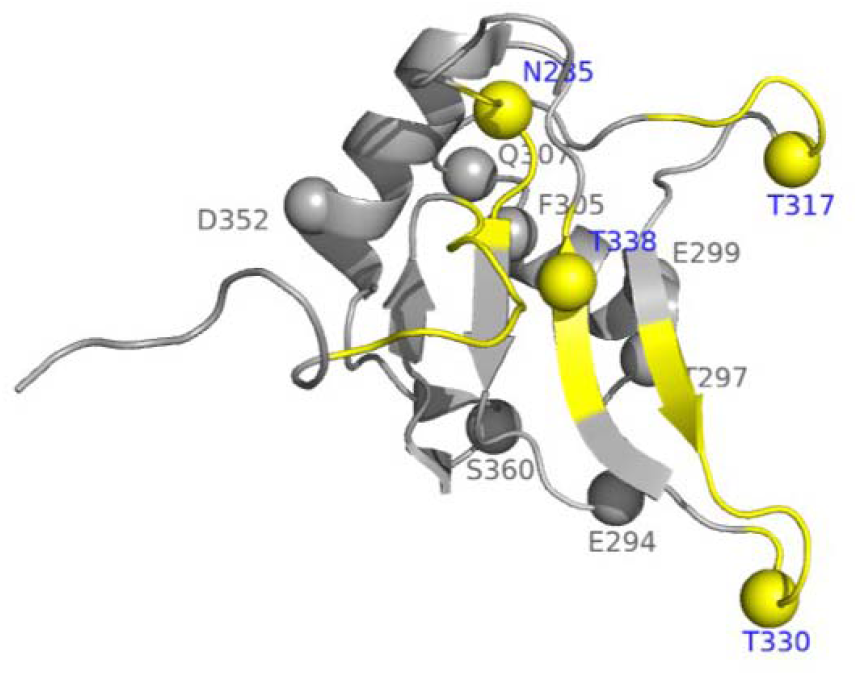
FUS-RRM tertiary structure (PDB ID: 2LCW) highlighting the nucleic-acid binding surface of the protein (marked in yellow) and the residues undergoing conformational exchange leading to formation of ES^CEST^. Residues shown in yellow spheres (R285, T317, T330, and T338) are present in the loop regions and at the terminal of a β-strand of the nucleic-acid binding surface.

### The ES^CEST^ of FUS-RRM is a partially unfolded state

Building on insights from the ES^CEST^ state stabilized at lower pH, we compared the random coil backbone ^15^N chemical shifts of FUS-RRM predicted by POTENCI^32^ with those measured using ^15^N-CEST (Figure 4A). The strong correlation (R^2^ = 0.74) between the two datasets indicates that ES^CEST^ indeed represents an unfolded conformation. To further investigate the nature of ES^CEST^, we acquired ^15^N-CEST data on FUS-RRM in the presence of 1 M urea, a chaotropic agent that is known to disrupt the protein structure and induce unfolding in proteins. The analysis showed that urea further stabilized the ES^CEST^ population, as evidenced by the global fit parameters (k_ex_ = 208±18.6 s^-1^, p_ES_ = 0.53±0.01) from a two-state model fit of the data (Figure 4B, 4C, and S4). Thus, the data suggest that the ES^CEST^ is an unfolded/partially unfolded state of the protein. A similar approach has been used recently by Deshmukh and co-workers to demonstrate that the dsRBD excited state is an unfolded state.^33^ We then compared the ΔG_GS-ES_ as obtained using ^15^N-CEST experiment with the ΔG_unfolding_ calculated using urea-induced unfolding of the FUS-RRM as measured by far-UV CD spectroscopy (Figures 4D and S5). The ΔG_ES-GS_ measured at pH 6.4, at pH 4.6, and at pH 6.4 in presence of 1 M urea was found to be 3.9, 3.1, and 3.1 kcal/mol (Table 1), respectively, and the ΔG_unfolding_ was found to be 4.8 kcal/mol (Table 2), confirming that the ES^CEST^ is a partially unfolded state, which is stabilized by lower pH (pH 4.6) and by 1 M urea.

**Table 1.** p_ES_ and k_ex_ obtained by fitting HARD-R_1ρ_ /_2ρ_ and ^15^N-CEST for RRM in various conditions and the corresponding free energy.

| HARD $R_{1\rho}$ and $R_{2\rho}$ | | | |
| --- | --- | --- | --- |
| RRM | $p_{ES}$ (%) | $k_{ex}$ ($ks^{-1}$) | $\Delta G$ (kcal/mol) |
| pH 6.4 | $6.2 \pm 5.0$ | $40.9 \pm 2.3$ | $1.6 \pm 0.5$ |
| pH 4.6 | $2.1 \pm 0.6$ | $0.3 \pm 0.2$ | $2.3 \pm 0.2$ |
| RRM + ATP | $p_{ES}$ (%) | $k_{ex}$ ($s^{-1}$) | $\Delta G$ (kcal/mol) |
| pH 6.4 | $2.7 \pm 1.4$ | $3.7 \pm 0.5$ | $2.1 \pm 0.3$ |
| pH 4.6 | $9.0 \pm 6.6$ | $23.9 \pm 1.8$ | $1.4 \pm 0.4$ |
| $^{15}N$-CEST | | | |

| **RRM** | **p<sub>ES</sub> (%)** | **k<sub>ex</sub> (ks<sup>-1</sup>)** | **ΔG (kcal/mol)** |
| --- | --- | --- | --- |
| pH 6.4 | 0.1 ± 0.0 | 0.3 ± 0.0 | 3.9±0.0 |
| pH 4.6 | 0.5 ± 0.1 | 0.1 ± 0.0 | 3.1±0.1 |
| **RRM + ATP** | **p<sub>ES</sub> (%)** | **k<sub>ex</sub> (ks<sup>-1</sup>)** | **ΔG (kcal/mol)** |
| pH 6.4 | 0.5 (fixed) | 0.1 (fixed) | 3.1±0.0 |
| pH 4.6 | 0.5 ± 0.0 | 0.9 (fixed) | 3.2±0.0 |
| **RRM + 1 M urea** | **p<sub>ES</sub> (%)** | **k<sub>ex</sub> (ks<sup>-1</sup>)** | **ΔG (kcal/mol)** |
| pH 6.4 | 0.5 ± 0.0 | 0.2 ± 0.0 | 3.1±0.0 |

**Table 2:** Fit parameters obtained by two-state (folded and unfolded) fitting of the fraction unfolded (as measured by far-UV CD) vs urea concentration data.

|  |  |
| --- | --- |
| ΔG<sub>unfolded</sub> (kcal/mol) | 4.8 ± 0.8 |
| m | 1.2 ± 0.2 |
| R<sup>2</sup> | 0.98 |

**Figure 4:**
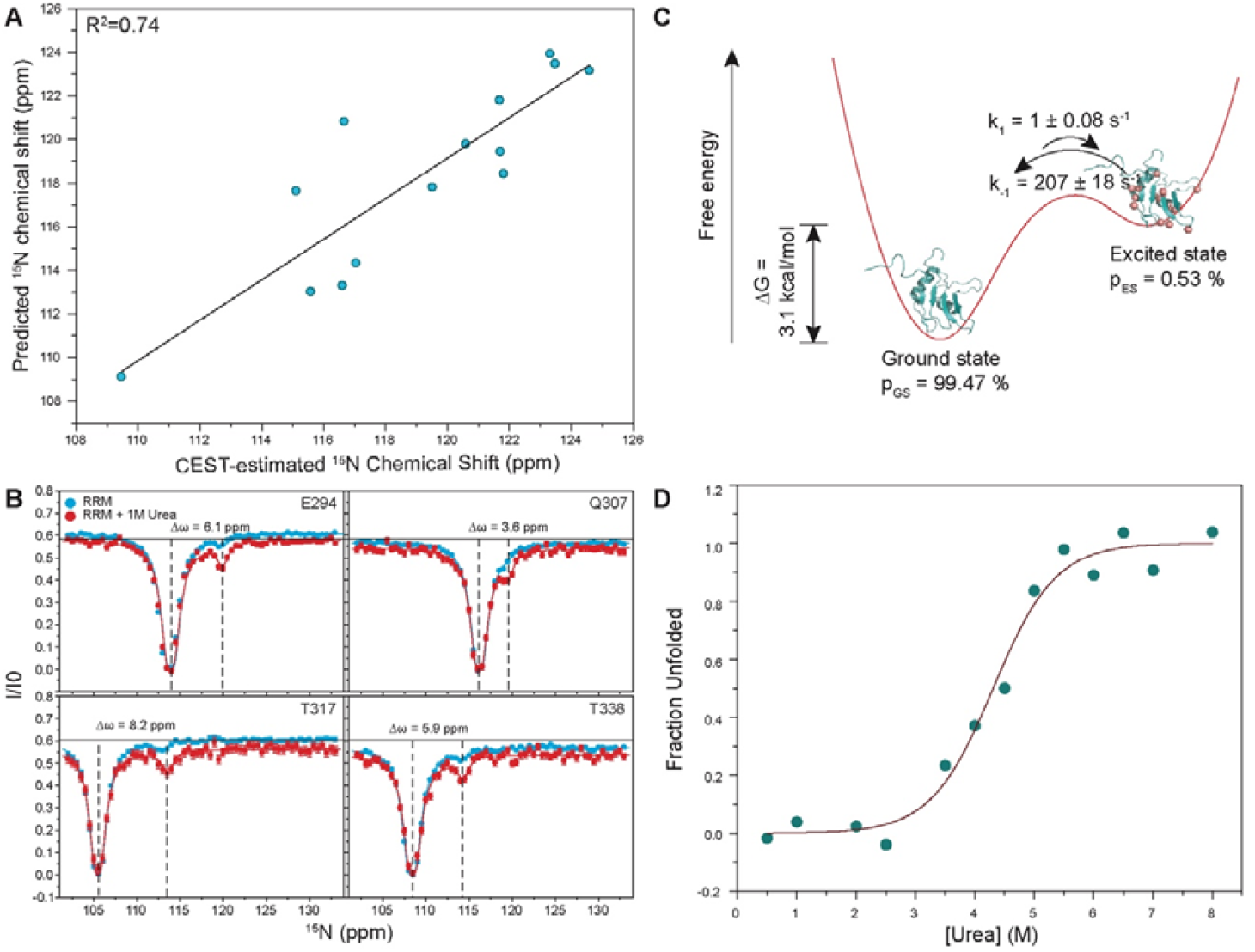
The excited state detected by ^15^N-CEST corresponds to an unfolded state. (A) Correlation between the CEST-estimated and predicted ^15^N chemical shifts for the residues showing an exchange. (B) An overlay of ^15^N-CEST profiles of FUS-RRM residues in absence (cyan) and presence (red) of 1M urea. The obtained χ^2^_red_ is 1.5 using fit to a two-state model. The vertical dashed lines show large chemical shift changes between the ground and excited state. (C) Representation of the energy landscape of FUS-RRM showing the two states, ground (folded) and excited (partially unfolded or unfolded), and their interconversion rates with fractional populations. (D) Two-state (folded and unfolded) fit to the plot of fraction unfolded (measured using far-UV CD) vs urea concentration to obtain ΔG_unfolding_.

To visually assess the conformational landscape of the FUS-RRM, we employed a generative deep-learning protein dynamics emulator, BioEMU, and sampled 300 lowest energy conformations.^34^ Clustering of the 300 sampled conformations highlighted 74 ensembles with different free energies (Figure 5). All the conformations were classified into folded and unfolded states based on their fraction of native contacts. The free-energy distribution of the conformations spanned from -4.6 to –2.1 kcal/mol, with an average ΔG of - 3.7 kcal/mol (Figure 5A). To assess the robustness of the program, we repeated the BioEMU sampling with 3000 conformations. The average free-energy obtained with increased sample size was similar (–3.5 kcal/mol) (data not shown), suggesting that an increase in the sampling size did not change the conformational landscape accessed by the program. While the lowest-energy structures converged to compact folded-like states, several alternative conformations were identified at slightly higher energies. These unfolded states retained β-sheet core but exhibited differences in loop flexibility and particularly α2 helix, which gets unfolded in some of the conformations (Figure 5B and S6). BioEMU also predicted a shorter stretch of β2 and longer stretch of β4 strand, in contrast to NMR solved structure (PDB ID 2LCW). The presence of such partially unfolded states is consistent with the ^15^N-CEST measurements, further confirming the nature of the ES^CEST^ state.

**Figure 5:**
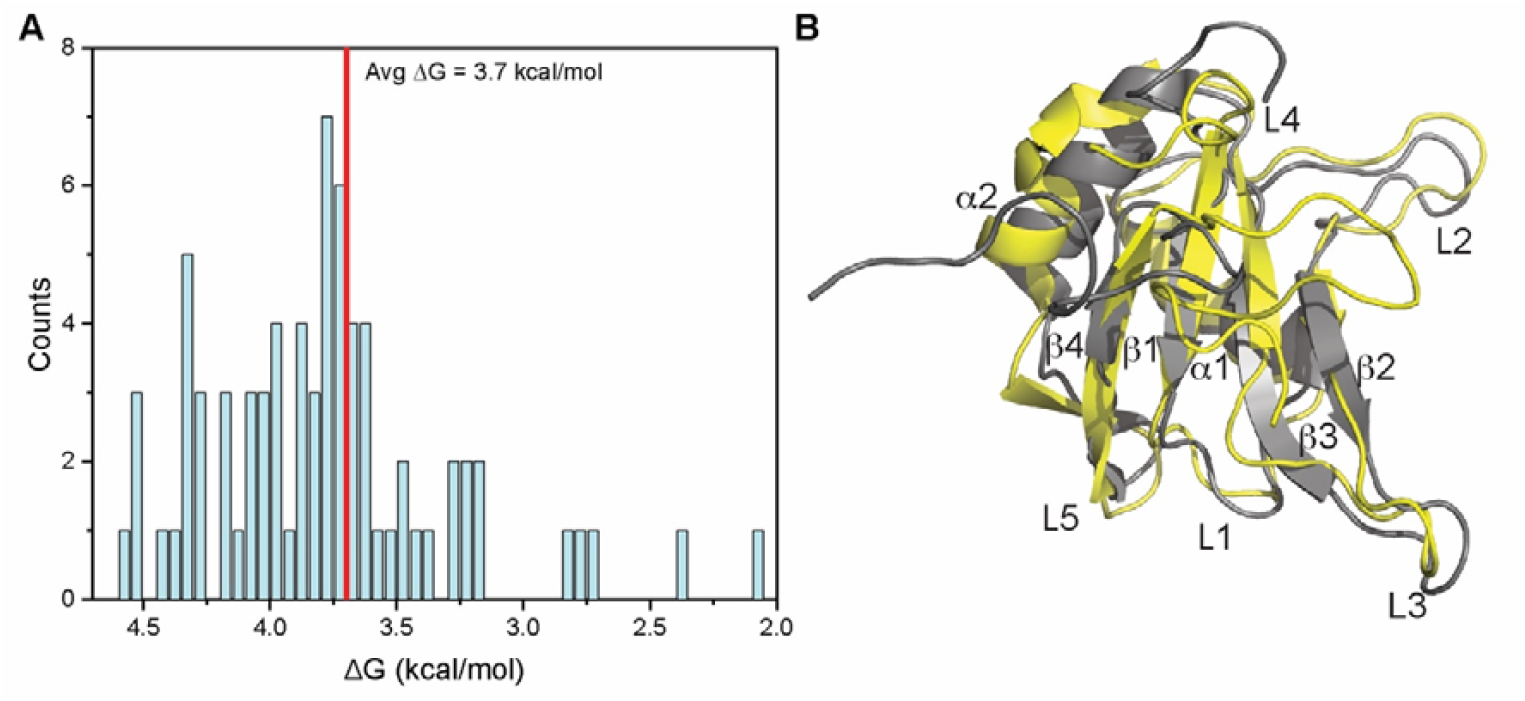
(A) Counts of conformations in different clusters obtained from BioEMU sampling of FUS-RRM conformations plotted against corresponding ΔG values (free energy difference in comparison to the lowest energy folded conformer). Average ΔG marked (3.7 kcal/mol) by vertical red line is very similar to the ΔG obtained by ^15^N-CEST measurements (3.9 kcal/mol) at pH 6.4. The histograms were constructed using a bin size of 0.06. (B) An overlay of one of the representative partially unfolded conformation (yellow) from the 300 sampled conformations with that of the NMR structure of FUS-RRM (PDB ID: 2LCW). Secondary structure segments have been marked on the structure. While the position of loops L2 and L3 is preserved (part of RNA-binding surface), α2 helix shows unfolding and the β2 and β4 strands show change in relative lengths, suggesting partial unfolding of the protein.

### ATP modulates the ES^CEST^ of FUS-RRM

ATP acts as an energy currency for most biochemical reactions carried out in cells. In addition to that, ATP seems to alter the solubility of proteins in the mM range. For example, previous studies have reported that ATP addition modulates the fibrillations of amyloidogenic proteins, such as amyloid-β, tau, and FUS.^35–37^ High ATP in these cases increases the aggregation rate by mediating protein-protein interactions and liquid-liquid phase separation, leading to amorphous aggregates. In contrast, Patel et al. have proposed a new role for ATP in increasing the solubility of amyloidogenic proteins.^38^ The IDP (intrinsically disordered protein) investigated by Patel et. al. is a Gly-rich low complexity domains (LCD), which suggest that ATP may affect amyloid formation differently, and depends on the specific structural and chemical properties of the protein, as has also been suggested using tau protein.^35^ To test ATP binding to FUS-RRM, we acquired two-dimensional 2D ^15^N-^1^H heteronuclear multiple quantum coherence (SOFAST-HMQC) spectra titrated with different ATP concentrations at pH 6.4 (Figure S7). For each titration point, non-overlapping residues showing a chemical shift perturbation (CSP) were fit globally to a two-state binding process using TITration ANalysis software (TITAN).^39^ The fitted binding constant (K_d_) of 6.4±0.1 mM is consistent with ATP binding in the mM range.^31^

To examine how ATP influences the ES^CEST^ state, we performed ^15^N-CEST experiment in presence of 10-fold excess of ATP. A 10-fold excess of ATP was used to study the dynamics of the ATP-bound state, as a higher ATP ratio led to increased aggregation kinetics at pH 6.4 (Figure S8) making the NMR data measurement difficult. Upon analysis, the same 13 residues that showed the characteristic minor dip in the ^15^N-CEST profiles at pH 6.4 were identified (Figure 6A, blue and Figure S9). After fitting in ChemEx, a decrease in k_ex_ to 58.9 s^-1^ and an increase in p_ES_ to 0.49% was observed, thereby indicating an increase in ES^CEST^ population (Figures 6A and 6B) when compared to the data measured in the absence of ATP (k_ex_ is 257 s^-1^ and p_ES_ is 0.14%). Interestingly, at pH 4.6, while ATP supplementation did not perturb the ES^CEST^ population (p_ES_ is 0.48%) when compared with that in the absence of ATP (p_ES_ is 0.5%), an increase in the k_ex_ from 142 to 895 s^-1^ was observed (Figures 6C, 6D, and S10).

**Figure 6.**
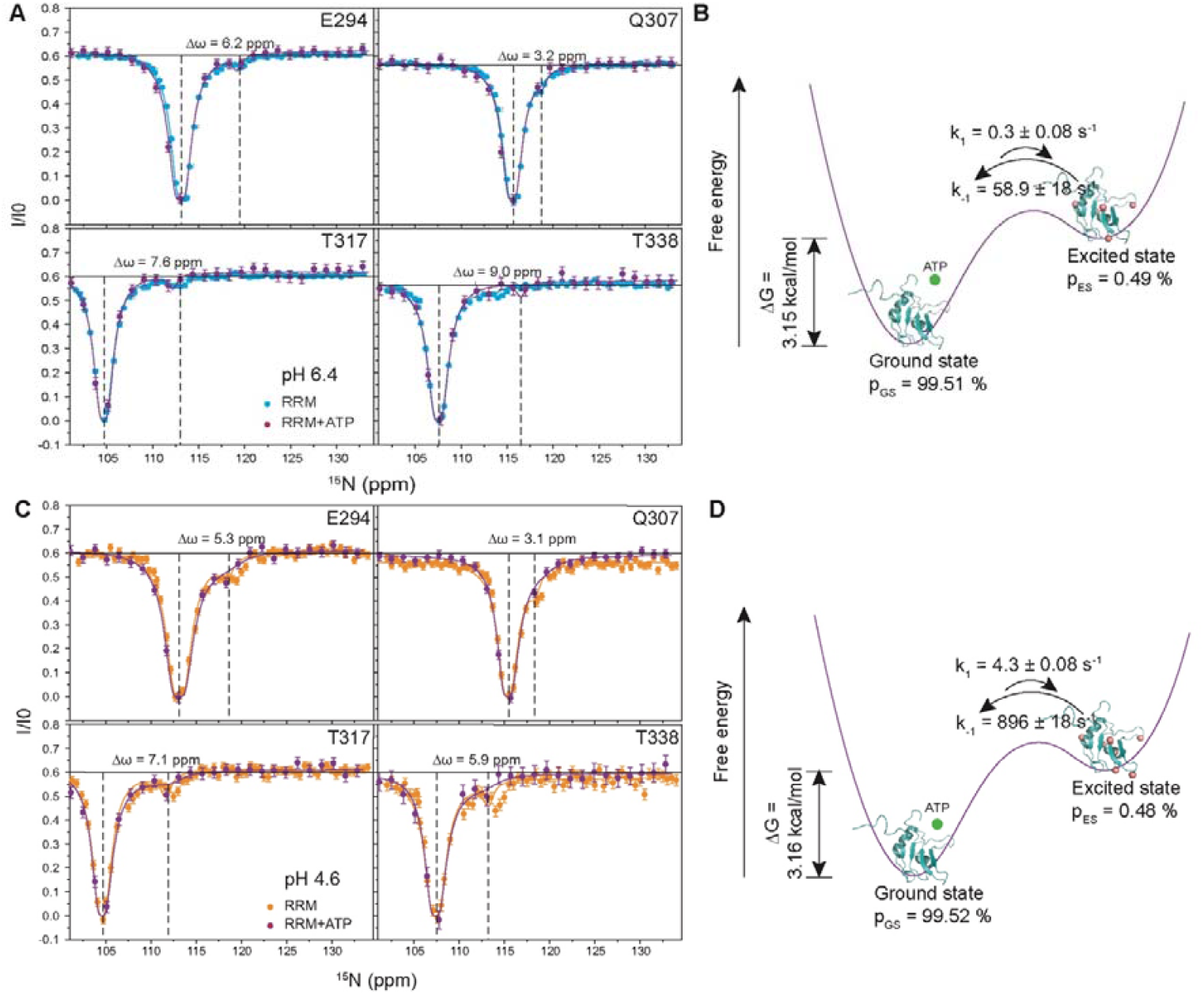
(A) An overlay of ^15^N CEST profiles of representative FUS-RRM residues at pH 6.4 with and without ATP and with corrected offsets for ground state of each residue. The obtained χ^2^_red_ is of 1.2 using fit to a two-state model. (B) Energy landscape of FUS-RRM with ATP showing the two states and their interconversion rates with fractional populations at pH 6.4. (C) ^15^N CEST profiles of representative FUS-RRM residues at pH 4.6 with ATP (red) when compared with the same in the absence of ATP. The obtained χ^2^_red_ is 0.6 using fit to a two-state model. (D) Energy landscape of FUS-RRM with ATP showing the two states and their interconversion rates with fractional populations at pH 4.6. The green circle has been used to represent ATP. The vertical dashed lines show large chemical shift changes between the ground and excited state.

### ATP modulates the ES^HARD^ of FUS-RRM

In order to understand if ATP affects fast μs-ms dynamics, we used similar adiabatic R_1ρ_ and R_2ρ_ relaxation dispersion, as reported earlier (Figure S11).^29^ In the presence of ATP, fast µs-ms dynamics is significantly quenched as reflected by both the average k_ex_ values and the p_ES_. The average k_ex_ reduces to 3.7 kHz from 40.9 kHz as measured earlier in the absence of ATP.^29^ The measured k_ex_ rates have been mapped onto the structure of the FUS-RRM in Figure 7A. The population of the ground state is shown in a scatter plot (Figure 7B). The average ground state population increases to 97.3% (in presence of ATP) from 93.8% in absence of ATP, thereby indicating a destabilization of ES^HARD^ under ATP conditions.

**Figure 7.**
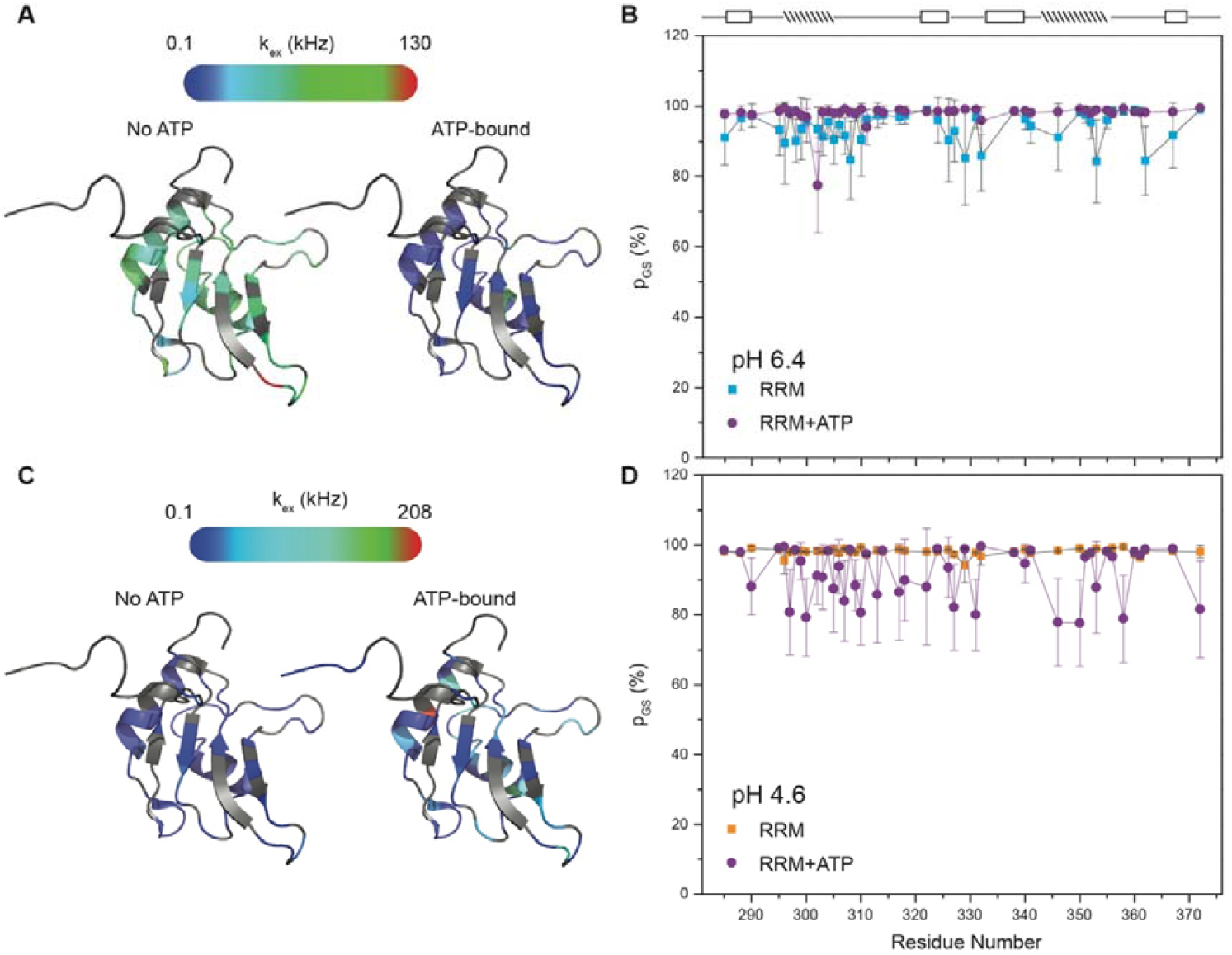
ATP-induced perturbation on fast µs-ms dynamics of FUS-RRM, measured at pH 6.4 and pH 4.6. (A) k_ex_ rates mapped to the FUS-RRM tertiary structure (PDB ID: 2LCW) and the population of the ground state in (B) at pH 6.4, respectively. (C) k_ex_ rates mapped to the FUS-RRM tertiary structure and the population of the ground state in (D) at pH 4.6, respectively. The residues highlighted in grey do not fit any model and, hence, have not been included in the discussion. The secondary structure of FUS-RRM is indicated on the top of panels B and D, with boxed and dashed rectangles denoting strands and helices, respectively.

At pH 4.6, supplementation of ATP led to an increase in both k_ex_ and p_ES_ (Figure 7C). The average k_ex_ increased from 0.2 kHz to 24 kHz in the presence of ATP, and the ground state population decrease significantly (average p_GS_ = 91%) upon ATP-binding at the lower pH (Figure 7D). This indicates that ATP binding significantly increases the fast μs-ms conformational dynamics at pH 4.6. Thus, it can be concluded that at lower pH, ATP stabilizes ES^HARD^ state.

### Aggregation kinetics and morphology of aggregates

Previously, we had observed that the rate of FUS-RRM aggregation increases significantly upon reducing the pH from 6.4 to 4.6.^29^ To further investigate the effect of ATP supplementation on FUS-RRM aggregation, we performed thioflavin T (ThT) kinetics at both the pH values 6.4 and 4.6, in the presence of ATP. At pH 6.4, ATP addition led to a significant increase in the aggregation kinetics (lag time reduced from 51.9 hours to 26.8 hours in the presence of ATP), while at pH 4.6, it was associated with a reduction in the rate of aggregation (lag time increases from 6 hours to 11.1 hours in the presence of ATP) (Figures 8A and 8B). The morphology of the aggregates, as observed by transmission electron microscope (TEM) imaging at the end of the kinetics measurement, showed the formation of amorphous aggregates (Figures 8C and 8D). These aggregates appeared highly heterogeneous in both shape and size, thus making quantitative analysis challenging. This implies that ATP-binding could modulate the final aggregate state of FUS-RRM, as has also been observed earlier in case of Aβ.^37^ The opposite effect of perturbed kinetics by ATP seen at the two pH values correlates well with the opposite perturbation of the observed slow and fast µs-ms dynamics, which suggests a direct role of conformational dynamics in modulating the aggregation.

**Figure 8.**
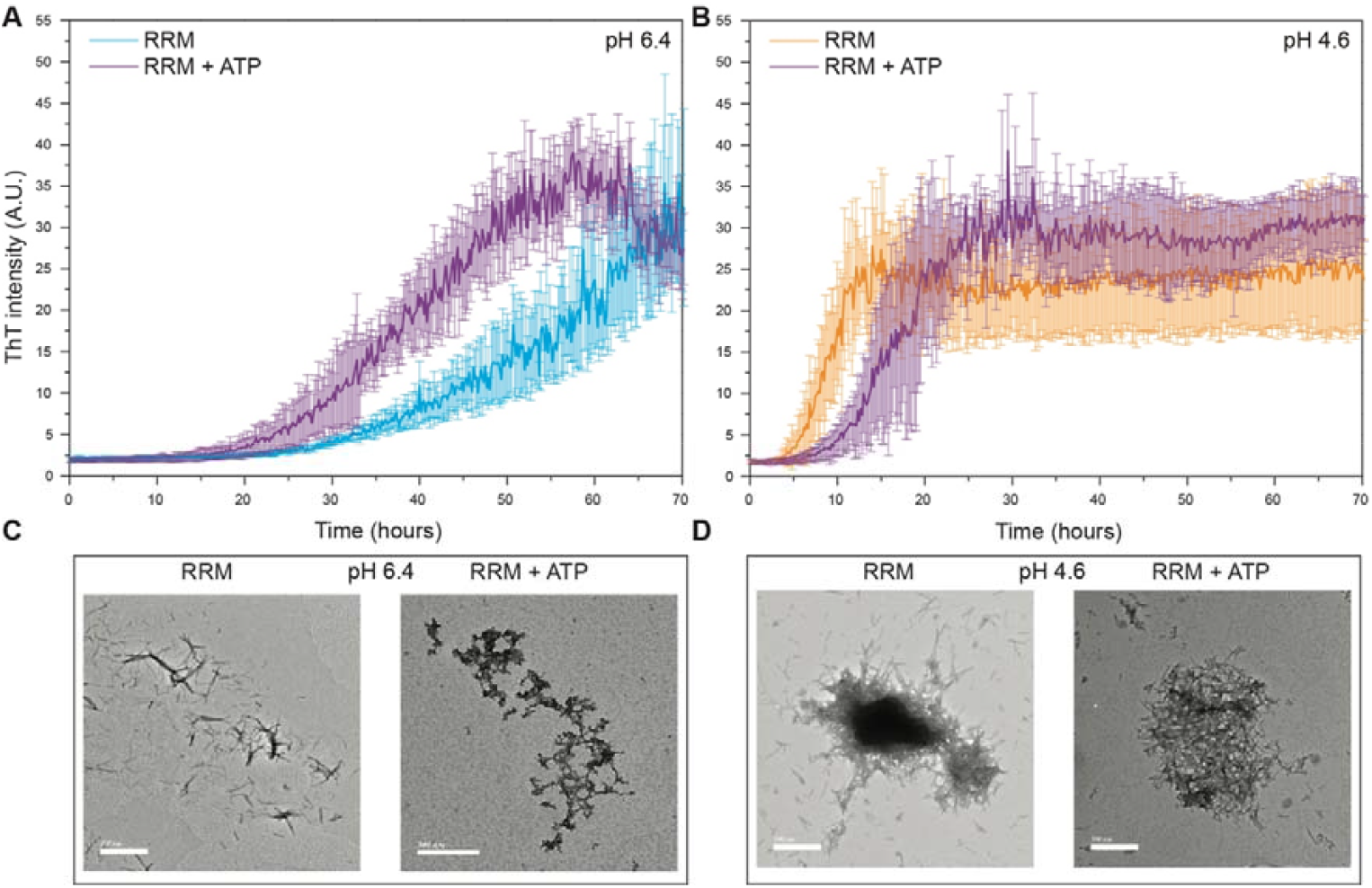
Aggregation kinetics for FUS-RRM as measured by monitoring ThT intensity with agitation in presence of ATP (10-fold) at (A) pH 6.4, and (B) pH 4.6. Aggregate formation for FUS-RRM, as imaged by TEM, are shown below the kinetics data acquired at the end of measurement. Data are represented as mean ± SD of three independent experiments.

## Discussion

The results from this study provide important insights into the conformational dynamics of FUS-RRM, a key domain involved in RNA binding and cellular regulation. By combining ^15^N CEST, adiabatic R_1ρ_, and R_2ρ_ relaxation dispersion data, this study uncovers the existence of two different excited states at different timescales. The exchange between these conformational states, and their relative stabilities are strongly dependent on environmental factors (e.g., pH and ATP that are probed in this study) and appear to be crucial for FUS-RRM’s functional roles, particularly its ability to aggregate under laboratory conditions and may be extrapolated to cellular conditions.

The current study reports that the FUS-RRM samples different excited states, including a low-populated excited state (ES^CEST^), identified using ^15^N-CEST NMR measurements, in addition to previously observed excited state, identified using HARD-R_1ρ_ /_2ρ_ NMR experiments (ES^HARD^), with a population of 0.14% exchanging at 257 s^-1^. The relaxation dispersion observed across the sequence, the stabilization (w.r.t. GS) of the ES^CEST^ from a ΔG = 3.9 kcal/mol to 3.13 kcal/mol at a lower pH (pH = 4.6), and a good correlation between the predicted ^15^N chemical shifts of the unfolded protein and CEST-estimated chemical shifts, suggest that the ES^CEST^ represents a partially unfolded state. The stabilization of this excited state in the presence of 1 M urea supported this hypothesis further.

Our study also shows that ATP binds weakly to FUS-RRM (K_d_ = 6.4 mM), but induces a significant perturbation in both slow and fast µs-ms dynamics. At pH 6.4, ATP-binding stabilizes ES^CEST^, while destabilizing ES^HARD^, and increases the rate of formation of aggregates; however, amorphous aggregates are formed under these conditions. On the other hand, at pH 4.6, ES^CEST^ remains more or less unperturbed in terms of population, but k_ex_ increases significantly from 142 to 895 s^-1^. Also, ATP-binding leads to stabilization of ES^HARD^ from 2% to 9%, and decreases the rate of aggregate formation. The contrasting effects of ATP on slow and fast µs-ms dynamics at pH 6.4 and 4.6, along with the corresponding changes in aggregation behavior, suggest a complex relationship between ATP, pH, and protein aggregation kinetics (Figure 9). This suggests that while population of ES^CEST^ is directly correlated with the rate of formation of aggregates, there is an inverse correlation between population of ES^HARD^ and the rate of formation of aggregates.

**Figure 9.**
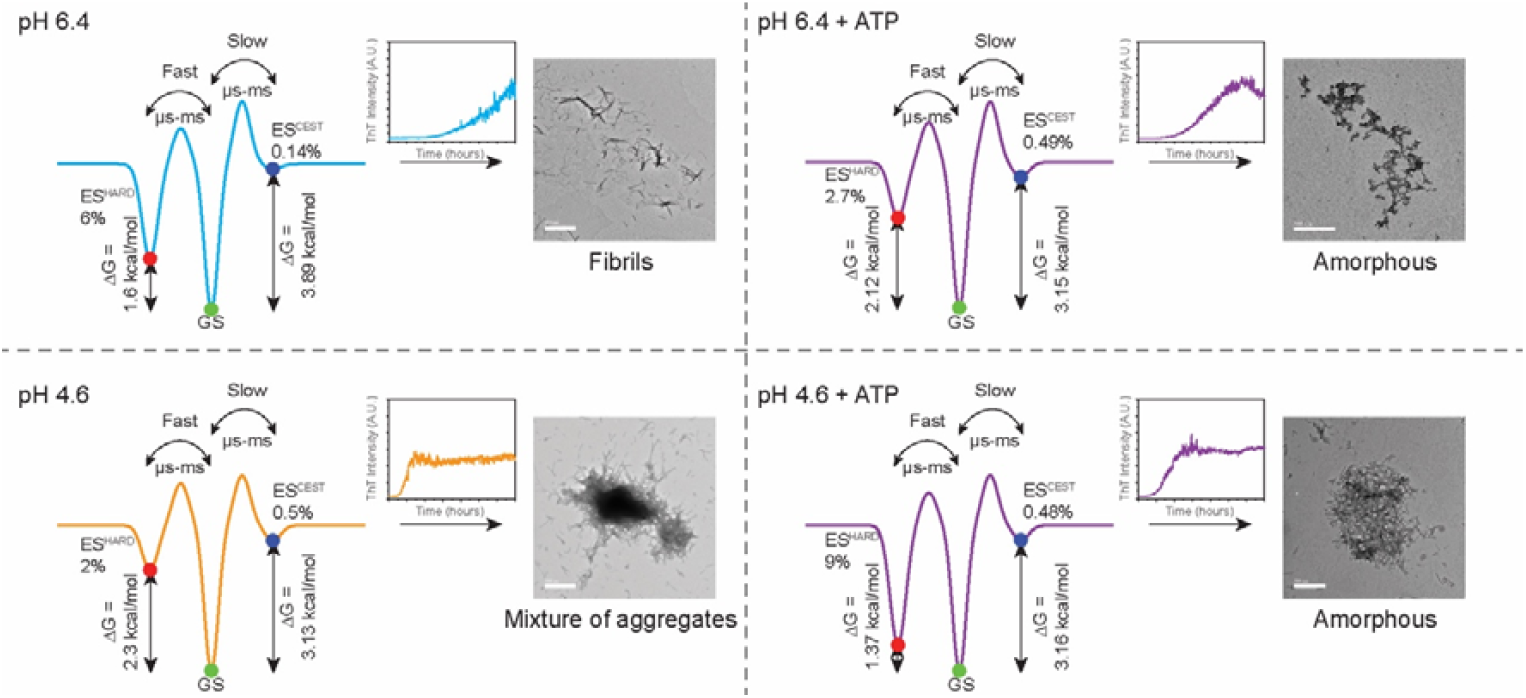
A cartoon representation of the free energy landscape of FUS-RRM showing conformational heterogeneity (a ground state, GS, and two excited states, ES^HARD^ and ES^CEST^, measured by HARD-R_1ρ_/2_ρ_ and ^15^N-CEST, respectively) and the associated aggregate formation monitored by TEM. On the arrow heads, the corresponding kinetics data as measured by ThT fluorescence are included to highlight a perturbation in the kinetics of aggregation. The experimentally determined free energy difference and population of each excited state have been mentioned.

Thus, the results of pH- and ATP-dependent aggregation kinetics suggest that there might be a certain optimum concentration of cellular pH and ATP levels relative to the protein concentration that can inhibit or delay the formation of amyloid aggregates. As cells undergo stress or aging,^40^ certain pH and ATP levels could stabilize a distinct conformation of the protein and thus prevent aggregation. These findings point to ATP being a modulator of protein solubility and aggregation—consistent with emerging literature that suggests that ATP has chaperone-like roles in cellular protein homeostasis.^38,41^

## Conclusion

It conclusion, it was observed that the FUS-RRM populates not only GS and ES^HARD^ but also ES^CEST^ exchanging with GS at slow μs-ms timescales. We showed using urea-perturbed ^15^N-CEST measurements and protein dynamics emulator that ES^CEST^ represents a partially unfolded state. A dynamic exchange was observed between these conformational substates in various environmental perturbations, including pH and ATP supplementation, used in this study. Further, a correlation between the populations of excited states (ES^HARD^ and ES^CEST^) and their exchange rates was observed to exist with the aggregation kinetics and aggregated states (fibrils and amorphous). Overall, from pH and ATP perturbations, we find that the environmental factors can re-shape the energy landscape, shifting populations and altering aggregation propensity. This work further underscores the critical role of conformational dynamics measurements to capture elusive states and integrating that data with structures to obtain the complete mechanistic picture of a given biological processes.

## Methods

### Protein overexpression and purification

The cDNA fragment encoding FUS-RRM (aa 282-376) cloned in pET28a vector was provided as a kind gift by Prof. Neel Sarovar Bhavesh, International Centre for Genetic Engineering and Biotechnology (ICGEB), New Delhi, India. Overexpression and purification of the recombinant protein was carried out in *Escherichia coli*, as described previously.^29^

### NMR spectroscopy

All the NMR experiments were recorded at 298 K on Ascend^TM^ Bruker AVANCE III HD 14.1 Tesla (600 MHz) NMR spectrometer equipped with a quad-channel (1H/13C/15N/19F) Cryoprobe (in-house). Spectral widths of 12 ppm (^1^H) and 32 ppm (^15^N) were used, for which an acquisition time of 141 ms was obtained in the direct dimension (F2). All the NMR spectra were processed in NMRPipe^42^ and analyzed in POKY.^43^

### ^*15*^*N CEST* measurements

^15^N CEST experiment utilizes irradiation of ^15^N z-magnetization using a weak radiofrequency (RF) pulse of amplitude B_1_ for a duration of T_ex_. At the end of T_ex_, intensities are recorded at each offset position across the ^15^N spectral width. Resonance intensities (I) at each offset are then quantified and normalized to corresponding intensities with T_ex_ of 0 (I_0_). The typical CEST profile consists of a ratio of I/I_0_ as a function of the offset value. For residue showing no exchange, the CEST profile shows a single dip in intensity. In contrast, residue undergoing exchange shows a major dip corresponding to the ground state and one or more minor dips for the excited state(s).^16^

^15^N CEST data was acquired using a B_1_ field of 30 Hz (other B_1_ fields like 10 Hz and 60 Hz were used for optimization of the minor dip intensity) with an exchange time (T_ex_) of 400 ms. A total of 64 planes were recorded between -912 and 972.8 Hz with a spacing of 30 Hz. For ATP-bound samples, the same sweep width was used with a spacing of 30 Hz, but the total frequency points recorded was 26 (to speed up the data collection due to faster aggregation in ATP-bound states) using the equation as described here.^44^ Similar strategy was also applied to ATP-free samples to test for the presence of dips at corresponding positions. The B_1_ field was calibrated using the method described by Guenneugues et. al. and used in the analyses.^45^

Global lineshape fitting was used to extract peak intensities from different planes of the CEST, using the FuDA package (https://www.ucl.ac.uk/hansen-lab/fuda/). The output data was then fit to the two-state exchange models using ChemEx (https://github.com/gbouvignies/ChemEx) to extract Δω, excited state population (p_ES_), and k_ex_ values. For the apo and urea samples, the global fit was performed by constraining R_2A_ = R_2B_ while for the ATP samples, the global fit was performed either by fixing p_ES_ and k_ex_.

### Heteronuclear Adiabatic Relaxation Dispersion (HARD)

HARD experiments were recorded and analysed under the different experimental conditions used for the study on the 600 MHz NMR spectrometer as described previously.^46–48^

The relaxation delays used for R_1ρ_ and R_2ρ_ were 0, 16, 32, 64, 128 and 176 ms. R_1_ experiments were acquired similarly to R_1_ρ and R_2ρ_ experiments but without using the adiabatic pulse during evolution. The delays used for the R_1_ experiment were 0, 48, 96, 192, 320, 480, and 640 ms.

### Aggregation kinetics

pH-dependent aggregation kinetics was carried out for 200 µL reactions with 50 µM protein and different molar ratios of ATP.MgCl_2_ (2-fold of MgCl_2_ used as otherwise mentioned) using 30 µM of thioflavin T (ThT) in aggregation buffer (10 mM sodium phosphate and/or sodium acetate, 100 mM NaCl) at 25°C, with agitation at 60 rpm in a 96-well plate (Fluoroskan, Thermo Scientific). The fibril formation was monitored by continuously measuring the ThT intensity at 475 nm upon excitation at 440 nm. ATP alone data was recorded at 10 mM ATP concentration.

### TEM imaging

After kinetics data were recorded, samples were centrifuged, and pellet was resuspended in Milli-Q water. 5 µL of the sample was transferred to a 300-mesh Formvar carbon-coated copper grid and incubated for 3 minutes. After incubation the sample was removed and the grid was negatively stained with uranyl acetate solution (1.5%). After 1 min, the solution was removed and grid was allowed to air-dry before imaging with transmission electron microscopy (JEM-2200FS) (TEM) at an accelerating voltage of 200 kV.

### Circular dichroism (CD) spectroscopy

Far-UV CD spectroscopy was recorded on a Jasco-815 spectrometer equipped with a thermal controller for 20 µM protein, incubated overnight in different urea concentrations (0 M to 8 M at an interval of 0.5 M). The scans were acquired from 210-260 nm in a 1 mm path length quartz cuvette with a bandwidth of 1 nm and a scanning speed of 50 nm/min. Data have been represented as Mean ± SD of two independent experiments after blank subtraction. The fraction of unfolded protein was calculated using CD ellipticity at 222 nm (equation 1), and was used to fit to a two-state model using (equation 2),

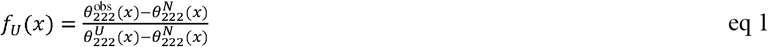

where, 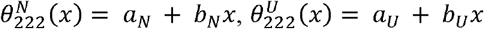, where, *x* is denaturant concentration (M), *a* and *b* are baseline intercepts and slopes, respectively.

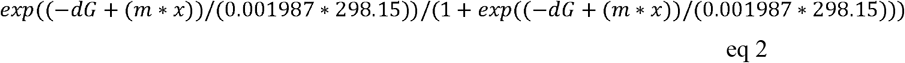

where, *dG* is the free energy of unfolding, and *m* is the slope.

### Conformational sampling using BioEMU

Conformational ensembles of FUS-RRM were generated using BioEMU.^34^ A total of 300 conformations were sampled. Sampled conformations were further clustered using Foldseek and clustering was performed using the following inputs:

n_write samples: -1, tm-score: 0.6, coverage: 0.7, and sequence identity: 0.95.

For each sampled conformation, the fraction of native contacts was computed using the BioEMU framework, and the corresponding free energy contribution was derived. Analyses were carried out directly on the backbone conformations generated during clustering. All calculations were performed in Google Colab (https://colab.research.google.com/). BioEMU was installed using !pip install bioemu. Visualizations of the structural ensembles were generated using PyMOL (https://www.pymol.org/).

## Acknowledgements

The authors acknowledge Prof. Neel Sarovar Bhavesh (ICGEB, New Delhi, India) for the FUS-RRM plasmids. OA acknowledges Prof. Jayant B. Udgaonkar (IISER Pune, India) for providing the facility to measure aggregation kinetics. OA acknowledges JC lab members for their inputs. The authors acknowledge the High-Field NMR facility at IISER-Pune (co-funded by DST-FIST and IISER Pune). J.C. acknowledges extramural funding from the Science and Engineering Research Board, Govt. of India (EMR/2015/001966 and CRG/2023/002931-C), Department of Biotechnology, Govt. of India (BT/PR24185/BRB/10/1605/2017), Department of Health Research, ICMR (R.11015/14/2023-GIA/HR), and the generous funding from IISER Pune. OA is grateful to DBT-JRF, Govt. of India, for providing a fellowship.

## Author contributions

J.C. and O.A. conceived the study and designed the approach to dynamically characterize the FUS-RRM. O.A. and A.A. prepared and purified protein samples. O.A. measured and analyzed all the NMR data, and A.A. performed the kinetics experiment. O.A. and J.C. have contributed toward manuscript writing.

## Data availability statement

All the raw NMR data have been uploaded to the Mendeley server and will be available to download.

All the raw intensities and relaxation rates data are available in supplementary tables (Tables S1 to S12).

## Table of Content Figure

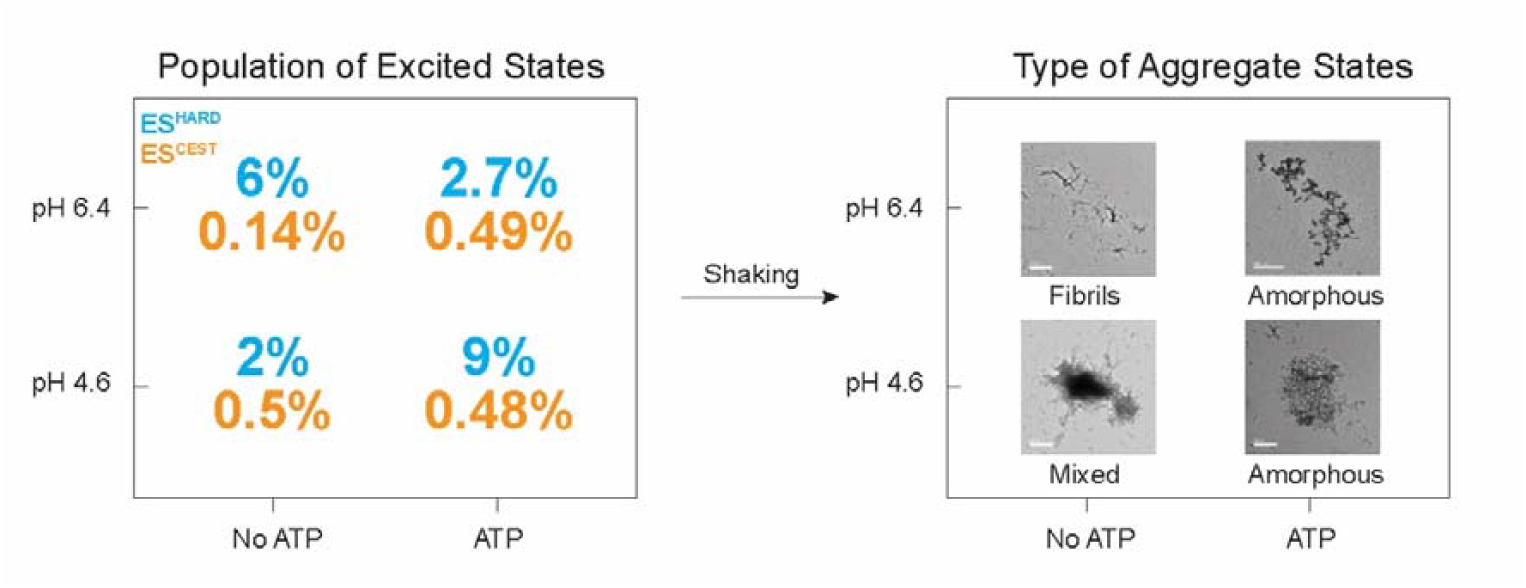

