## Supplementary for "H+ ions and ATP reshape the conformational landscape of an RNA recognition motif and regulate its fibrillation"

Dr. Jeetender Chugh

**Postal address**: C-115, Department of Chemistry, Main Building, Indian Institute of Science Education and Research (IISER), Dr. Homi Bhabha Road, Pashan, Pune 411008, India

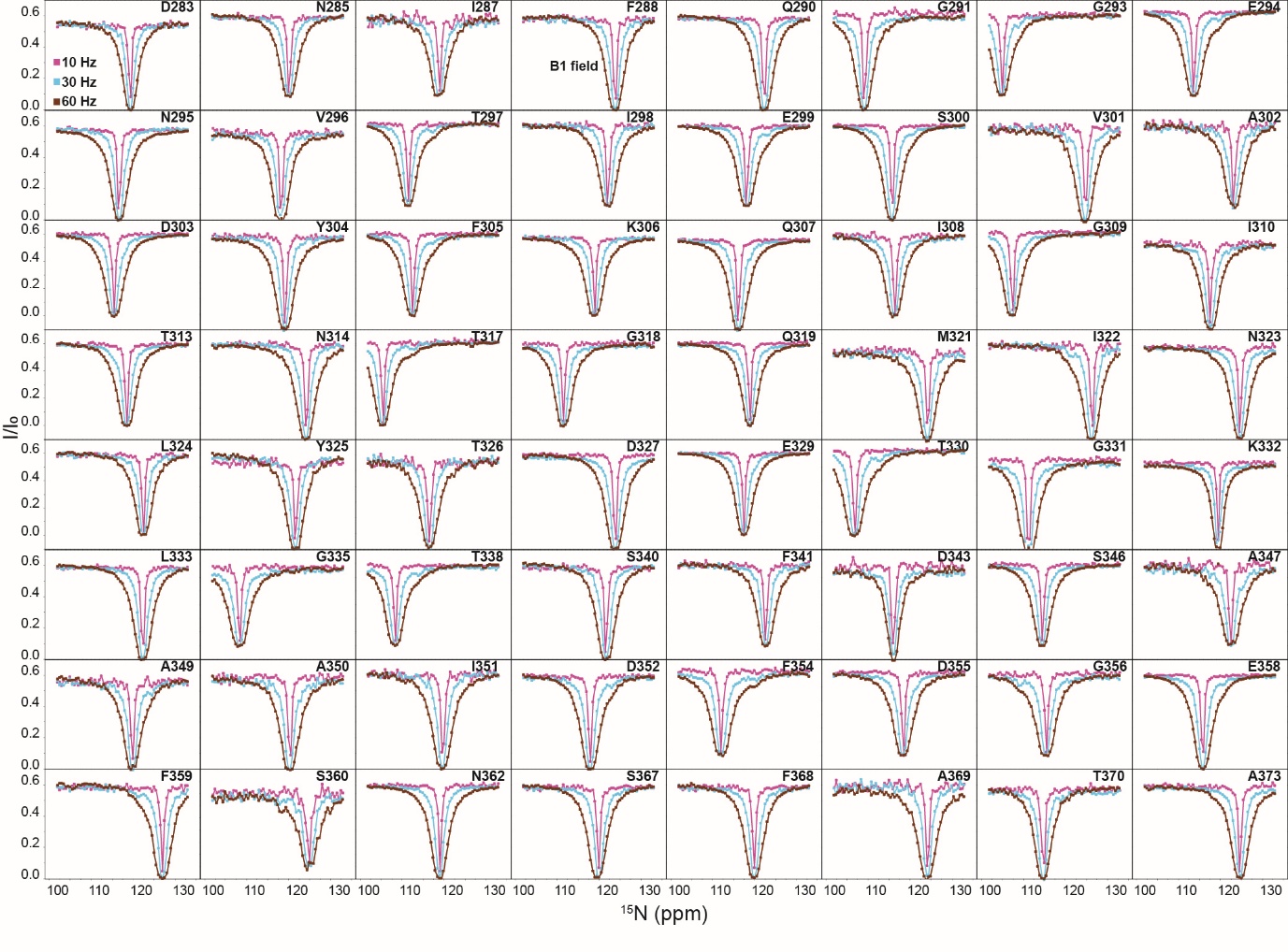
**Figure S1.** ^15^N CEST profiles with different B_1_ fields, pH 6.4.

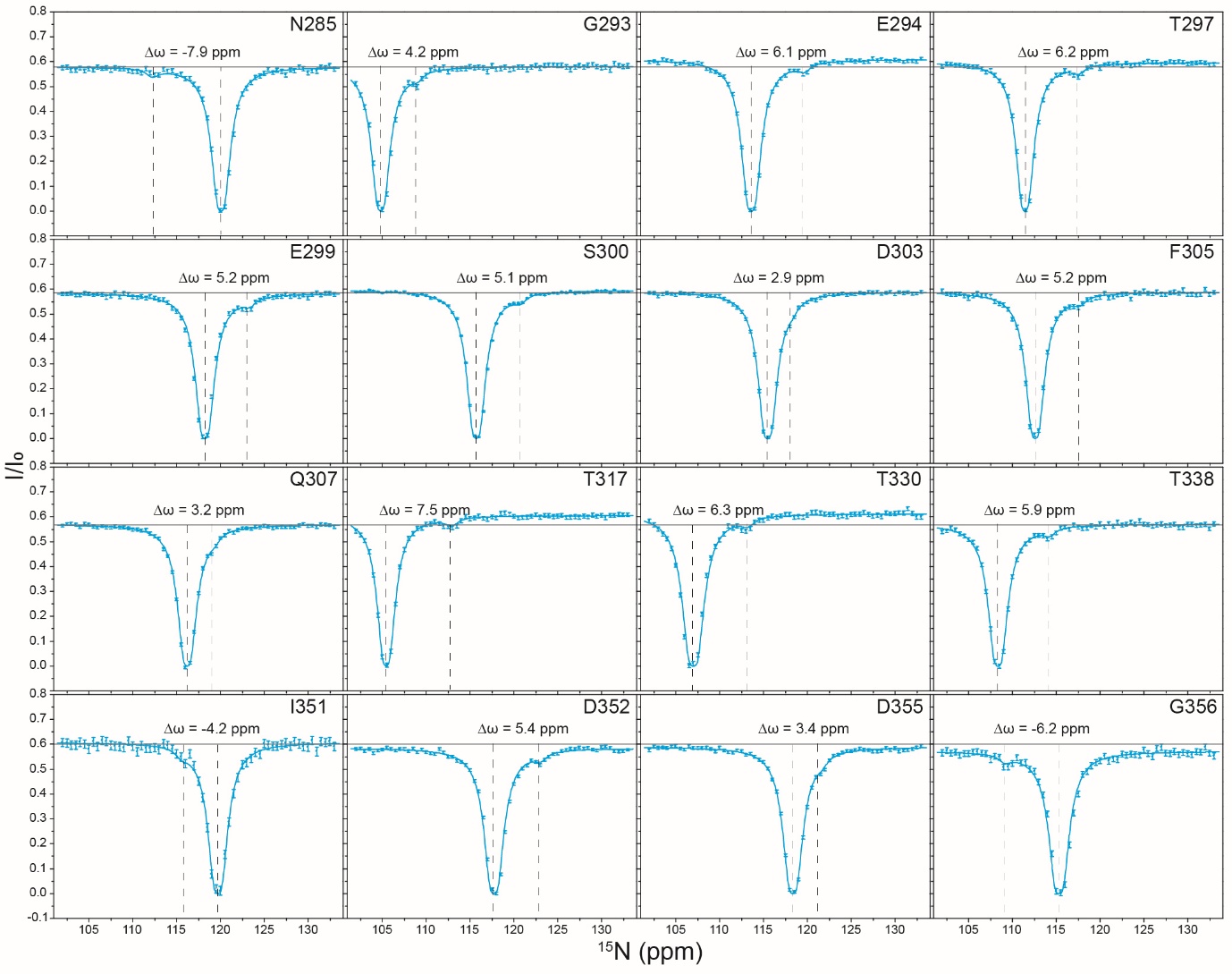
**Figure S2.** ^15^N CEST profiles of FUS-RRM residues showing minor dips measured at pH 6.4.

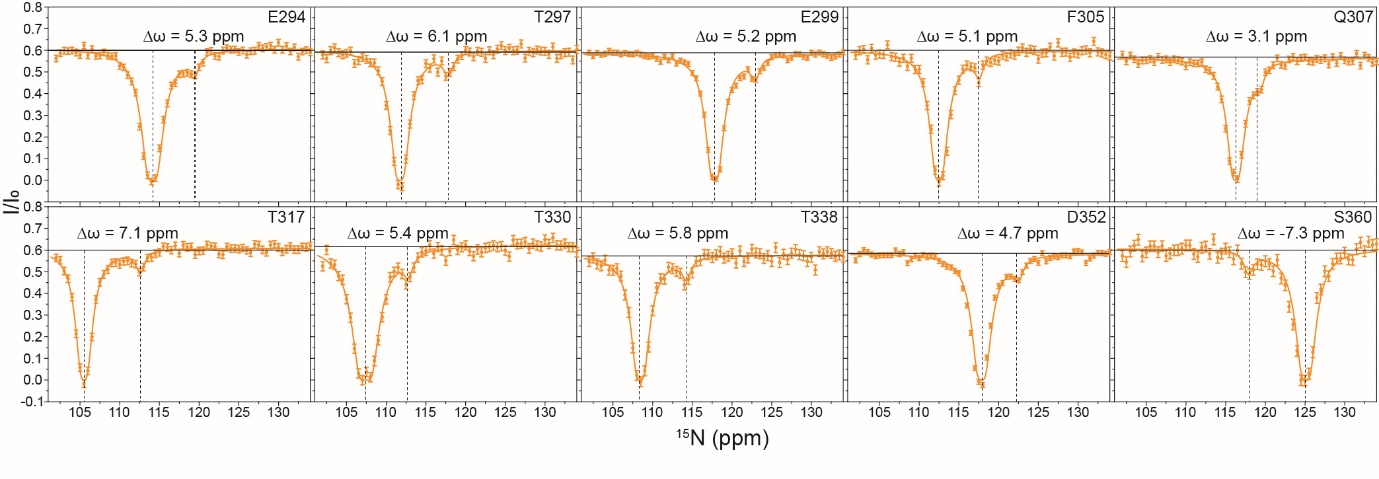
**Figure S3.** ^15^N CEST profiles of FUS-RRM residues showing minor dips measured at pH 4.6.

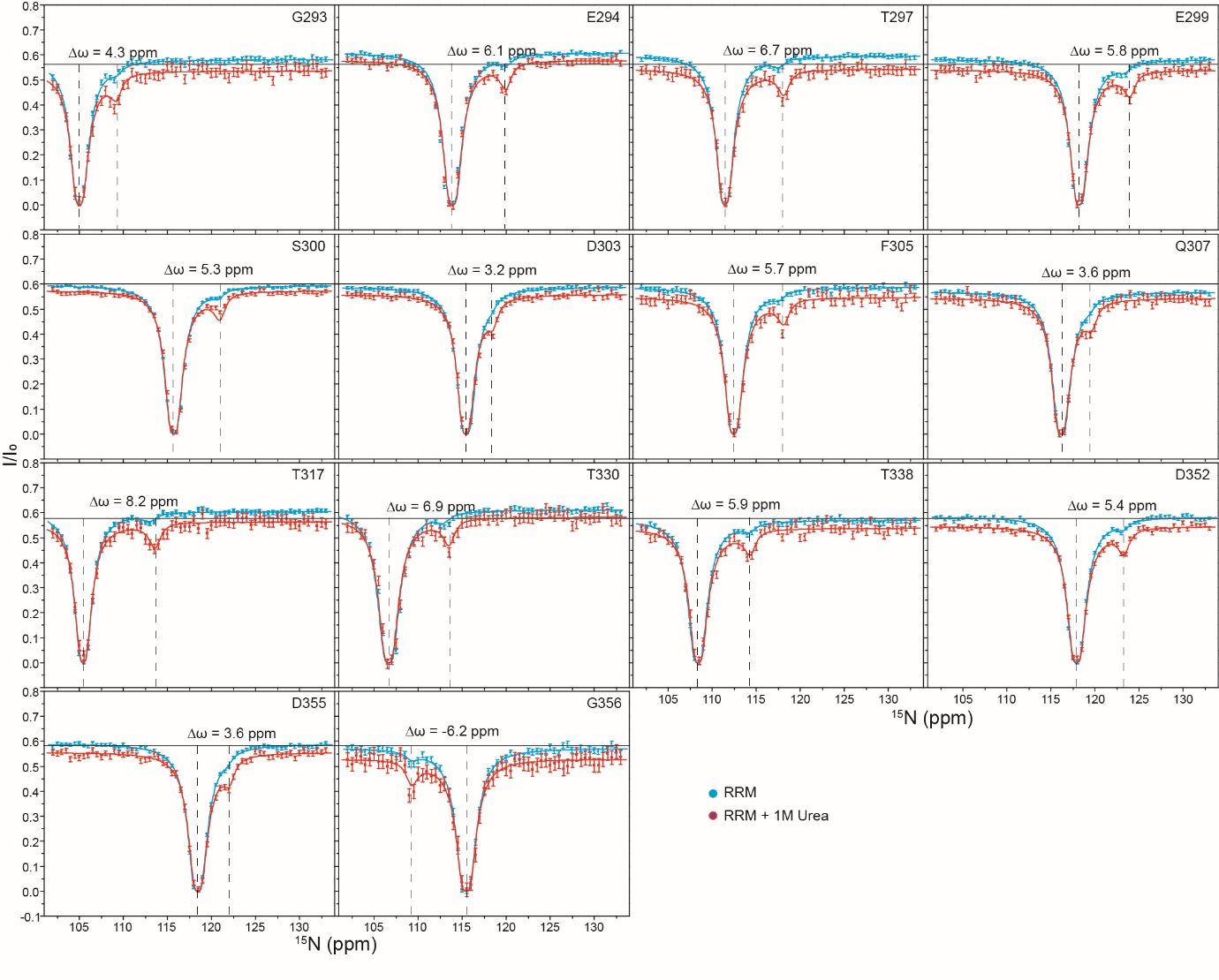
**Figure S4.** An overlay of ^15^N CEST profiles of FUS-RRM residues in presence of 1M urea.

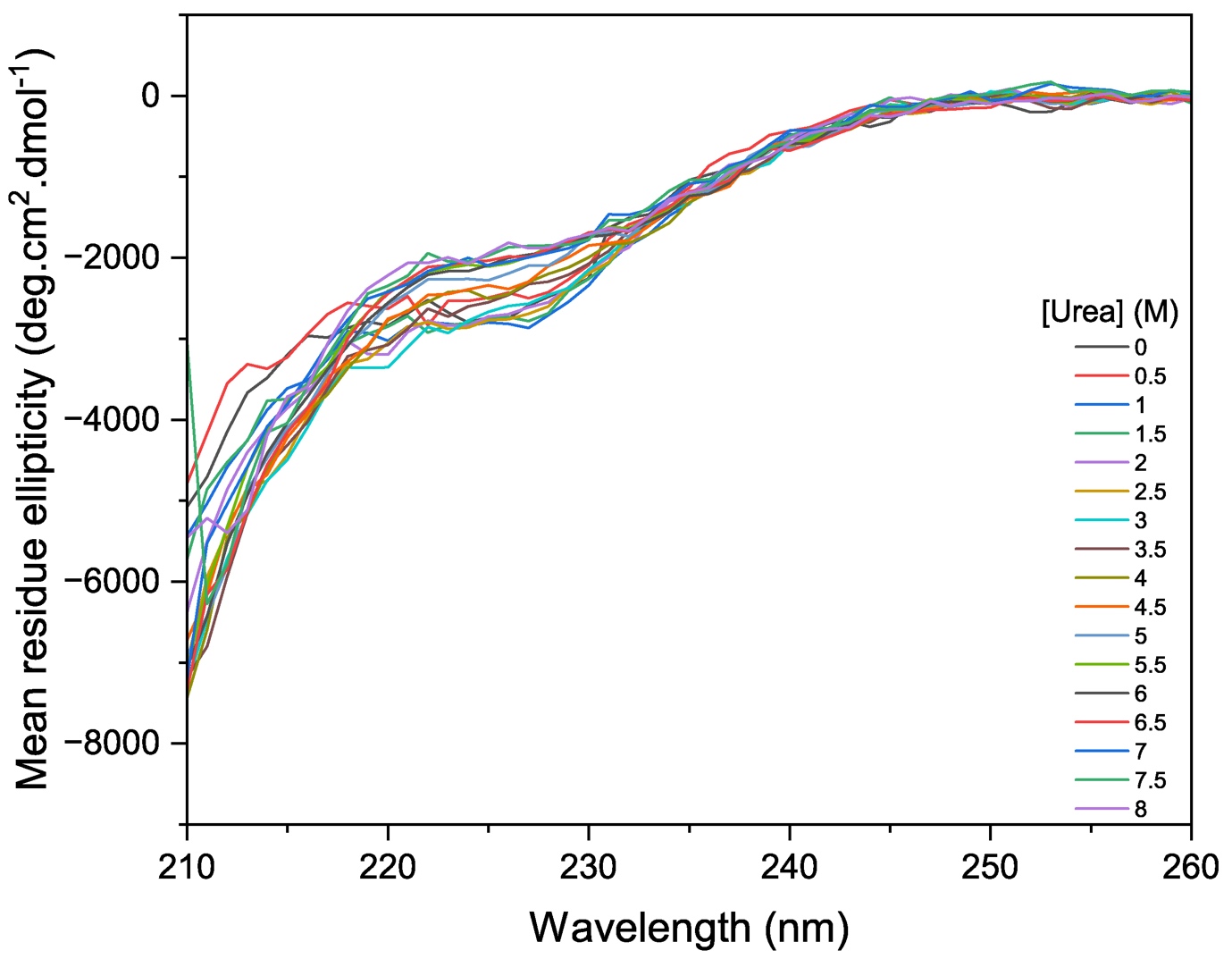

**Figure S5**: Far-UV circular dichroism spectrum of FUS-RRM measured as a function of urea concentration.

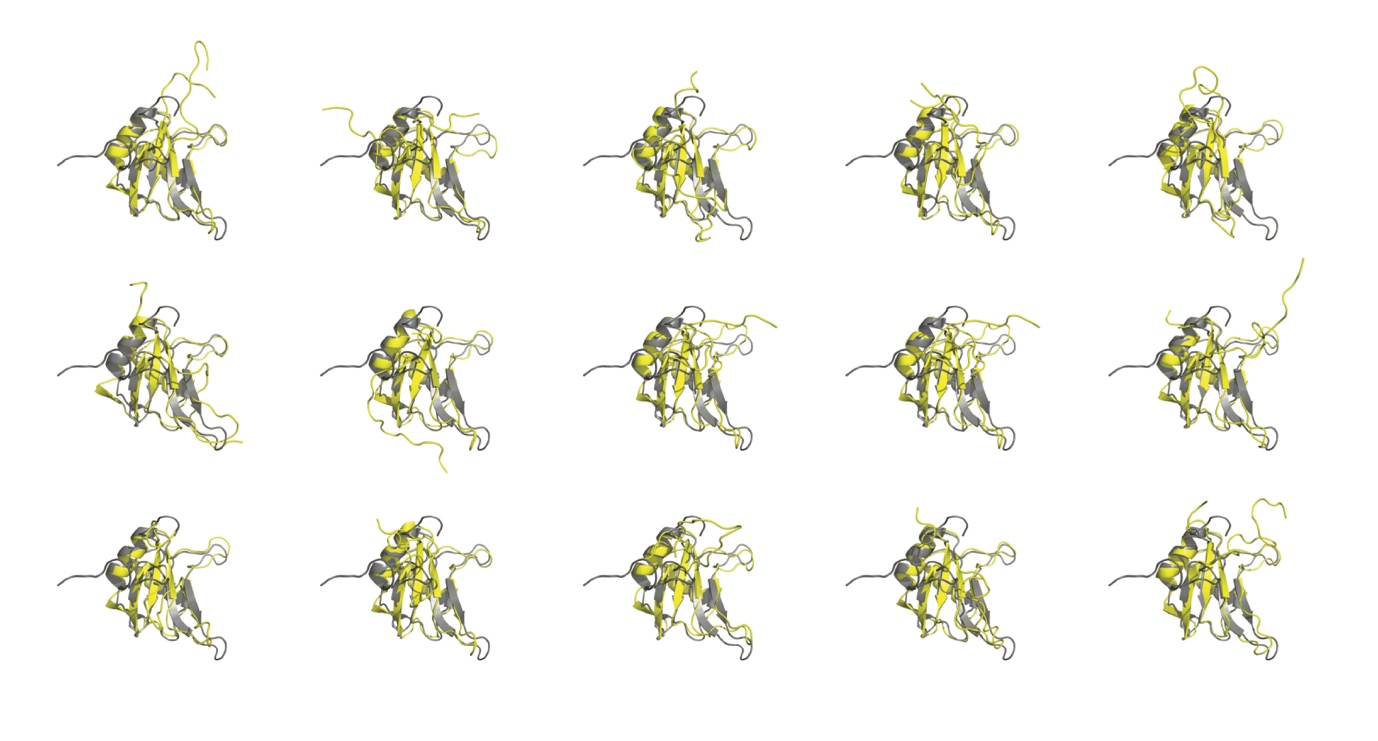

**Figure S6**: Overlay of the representative conformations (yellow) from the 300 sampled conformations with that of the NMR structure of FUS-RRM (PDB ID: 2LCW).

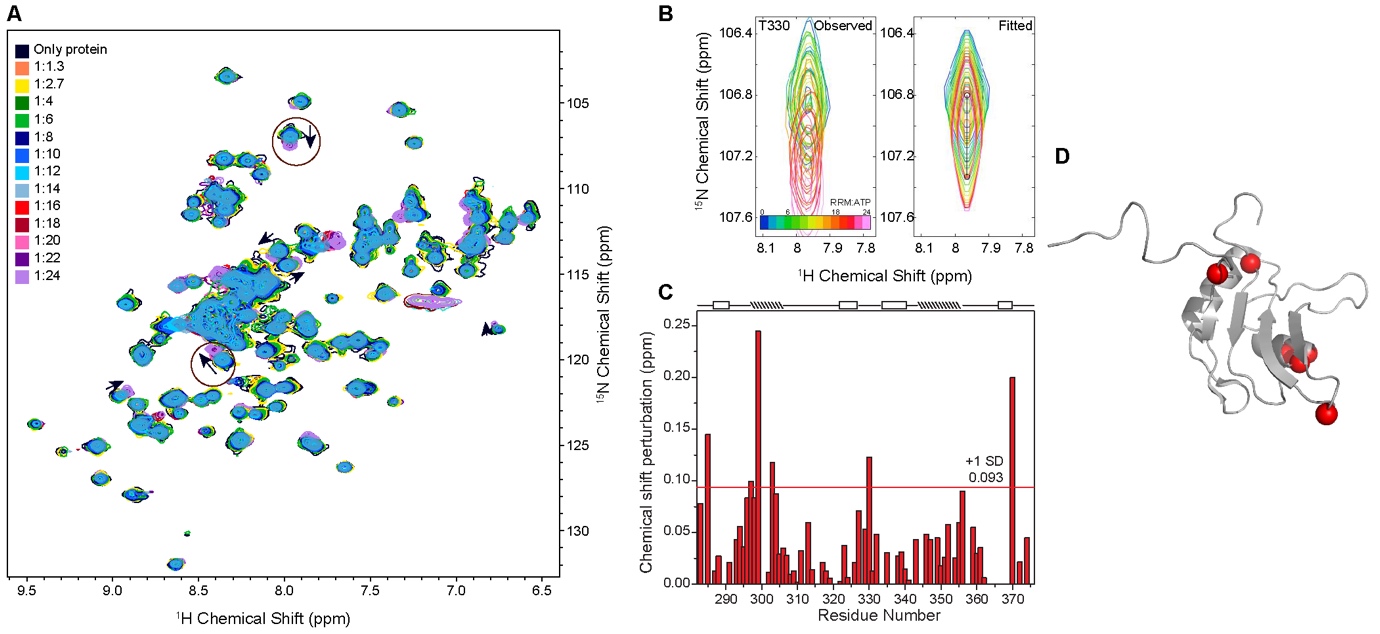
**Figure S7.** (A) An overlay of 2D ^15^N-^1^H HMQC spectra of FUS-RRM titrated with different molar ratios of ATP at pH 6.4. (B) Observed and fitted ^15^N-^1^H HMQC spectra of FUS-RRM upon titration with ATP. (C) Chemical shift perturbation (CSP) plot of residues titrated with the highest molar ratio of ATP used, and (D) shows the CSP of residues mapped onto the FUS-RRM PDB structure, which are above plus one standard deviation.

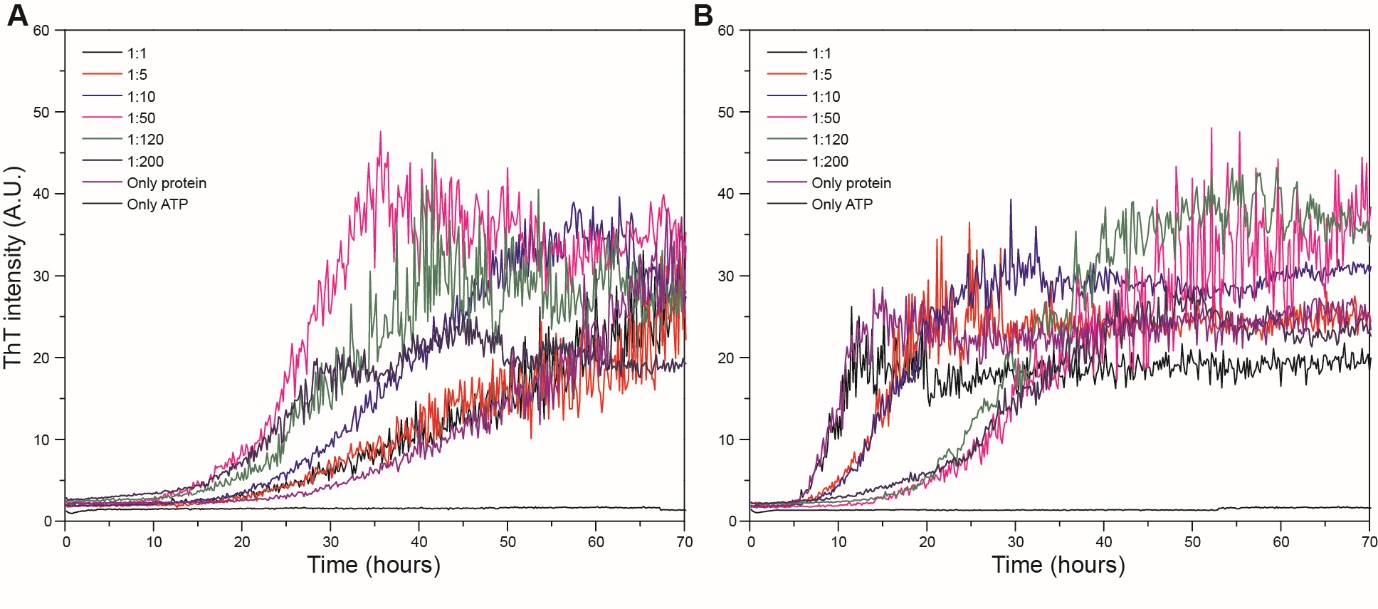
**Figure S8.** ATP-dependent aggregation kinetics of FUS-RRM measured at (A) pH 6.4 and (B) pH 4.6. Data are represented as mean ± SD of two independent experiments. Error bars are avoided for visual clarity.

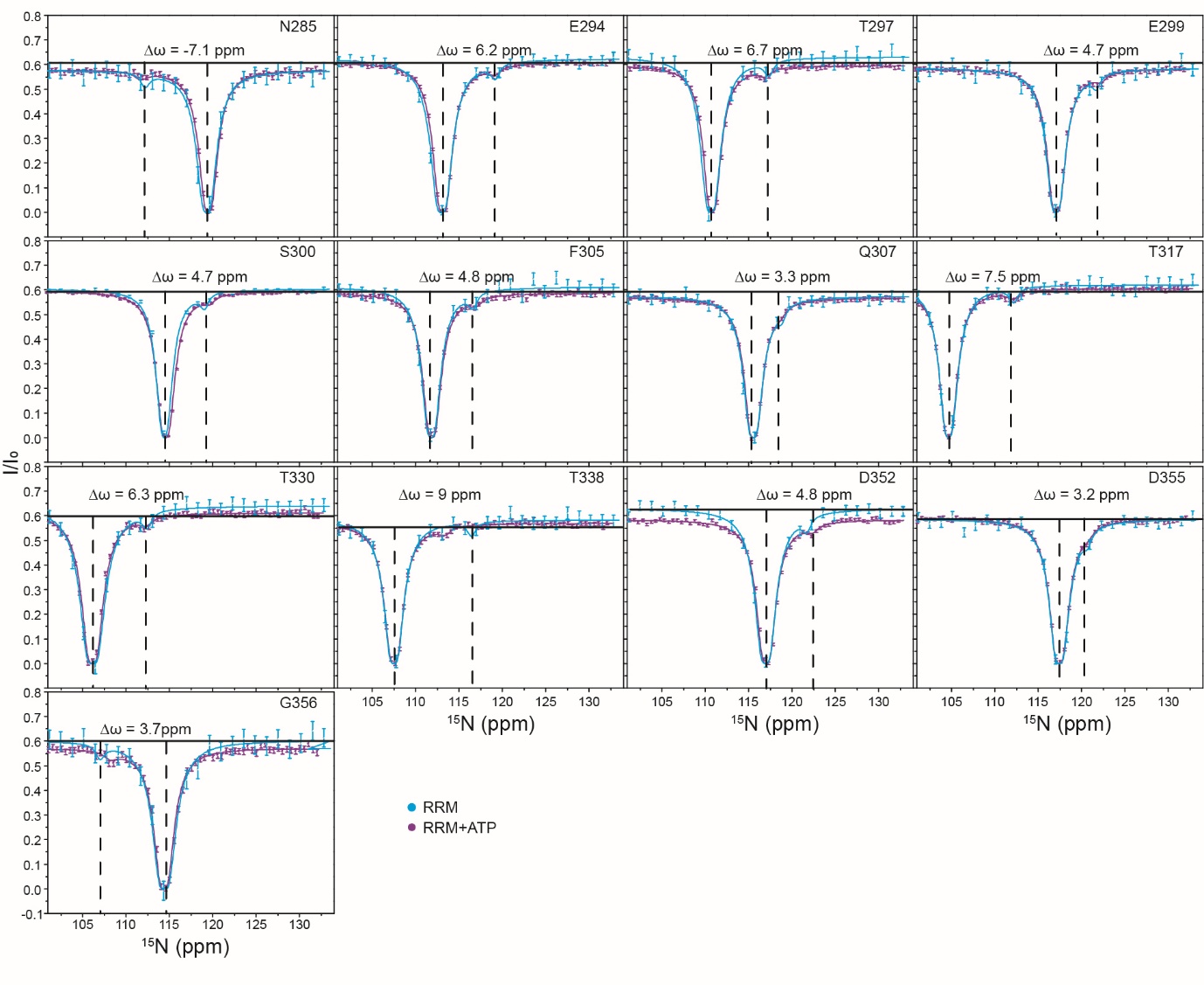
**Figure S9.** An overlay of ^15^N CEST profiles with and without ATP at pH 6.4.

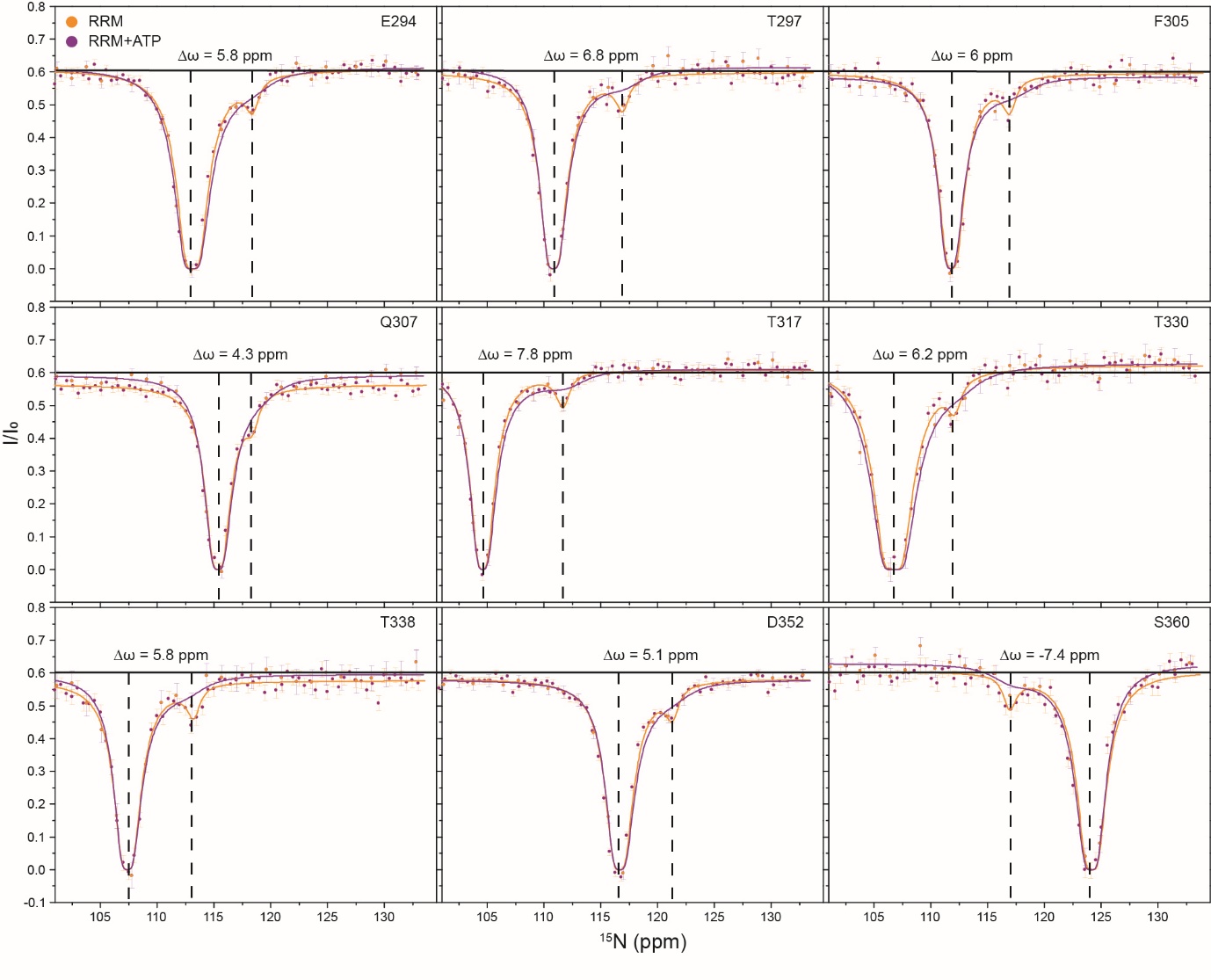
**Figure S10.** An overlay of ^15^N CEST profiles with and without ATP at pH 4.6.

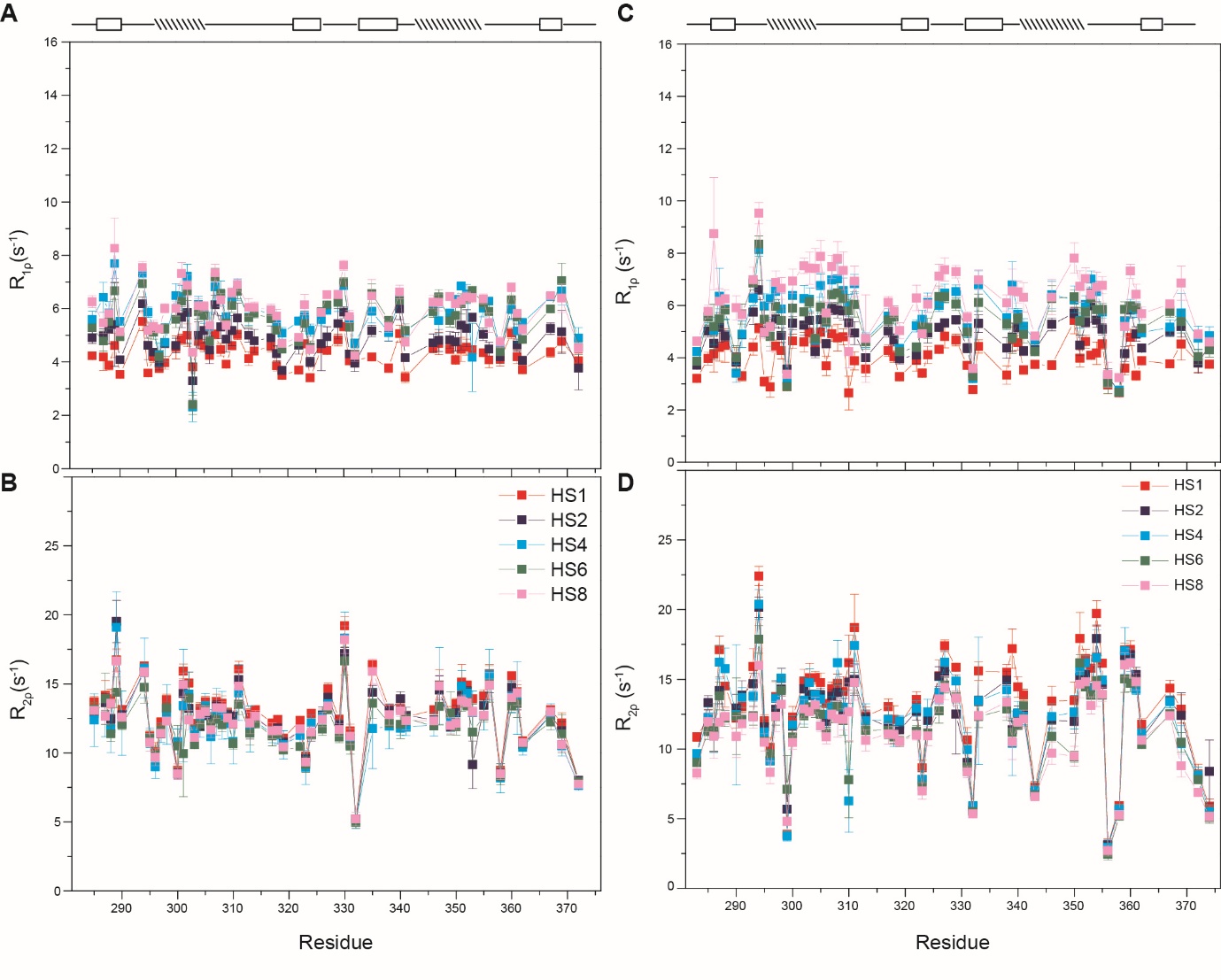
**Figure S11.** ATP-induced perturbation on fast µs-ms dynamics measured at pH 6.4 and 4.6. (A) R_1ρ_ and (B) R_2ρ_ relaxation rates at pH 6.4 and (C) and (D) at pH 4.6 at 25 °C. The trends in the relaxation rates for R_1ρ_ are HS1 (red) < HS2 (blue) < HS4 (sky blue) < HS6 (green) < HS8 (pink) and, (B) for R_2ρ_ trends are HS1 > HS2 > HS4 > HS6 > HS8.

**Table S1.** ^15^N CEST I/I_0_ data measured for FUS-RRM at pH 6.4 for different B_1_ field for various residues.

| Offset (ppm) | 10HzD283 | 30HzD283 | 60HzD283 | 10HzN285 | 30HzN285 | 60HzN285 | 10Hz I287 | 30Hz I287 | 60Hz I287 | 10HzF288 | 30HzF288 | 60HzF288 | 10HzQ290 | 30HzQ290 | 60HzQ290 |
| --- | --- | --- | --- | --- | --- | --- | --- | --- | --- | --- | --- | --- | --- | --- | --- |
| 102 | 0.53 | 0.55 | 0.55 | 0.58 | 0.58 | 0.6 | 0.5 | 0.55 | 0.57 | 0.56 | 0.58 | 0.59 | 0.58 | 0.59 | 0.59 |
| 102.5 | 0.54 | 0.55 | 0.56 | 0.57 | 0.58 | 0.59 | 0.57 | 0.55 | 0.56 | 0.58 | 0.59 | 0.58 | 0.58 | 0.59 | 0.6 |
| 103 | 0.55 | 0.54 | 0.56 | 0.57 | 0.57 | 0.59 | 0.5 | 0.59 | 0.57 | 0.57 | 0.59 | 0.59 | 0.59 | 0.58 | 0.59 |
| 103.5 | 0.54 | 0.55 | 0.54 | 0.56 | 0.56 | 0.58 | 0.57 | 0.53 | 0.55 | 0.59 | 0.57 | 0.59 | 0.57 | 0.58 | 0.6 |
| 104 | 0.55 | 0.54 | 0.56 | 0.56 | 0.58 | 0.58 | 0.56 | 0.52 | 0.58 | 0.58 | 0.59 | 0.59 | 0.58 | 0.59 | 0.6 |
| 104.5 | 0.55 | 0.54 | 0.55 | 0.58 | 0.56 | 0.58 | 0.55 | 0.56 | 0.57 | 0.57 | 0.58 | 0.58 | 0.57 | 0.58 | 0.6 |
| 105 | 0.54 | 0.57 | 0.54 | 0.57 | 0.57 | 0.59 | 0.56 | 0.51 | 0.59 | 0.58 | 0.58 | 0.59 | 0.58 | 0.58 | 0.6 |
| 105.5 | 0.54 | 0.53 | 0.55 | 0.58 | 0.57 | 0.58 | 0.53 | 0.55 | 0.59 | 0.59 | 0.59 | 0.58 | 0.58 | 0.58 | 0.6 |
| 106 | 0.55 | 0.54 | 0.56 | 0.56 | 0.58 | 0.59 | 0.58 | 0.57 | 0.56 | 0.58 | 0.59 | 0.6 | 0.57 | 0.58 | 0.59 |
| 106.5 | 0.54 | 0.54 | 0.55 | 0.58 | 0.57 | 0.58 | 0.55 | 0.55 | 0.55 | 0.58 | 0.59 | 0.59 | 0.58 | 0.58 | 0.59 |
| 107 | 0.54 | 0.54 | 0.55 | 0.56 | 0.58 | 0.59 | 0.54 | 0.52 | 0.55 | 0.58 | 0.58 | 0.6 | 0.59 | 0.58 | 0.59 |
| 107.5 | 0.53 | 0.53 | 0.55 | 0.58 | 0.56 | 0.58 | 0.57 | 0.59 | 0.56 | 0.57 | 0.57 | 0.58 | 0.58 | 0.58 | 0.58 |
| 108 | 0.54 | 0.53 | 0.55 | 0.6 | 0.58 | 0.58 | 0.57 | 0.53 | 0.58 | 0.59 | 0.56 | 0.6 | 0.58 | 0.58 | 0.59 |
| 108.5 | 0.55 | 0.53 | 0.55 | 0.57 | 0.57 | 0.57 | 0.57 | 0.56 | 0.53 | 0.56 | 0.57 | 0.6 | 0.59 | 0.58 | 0.59 |
| 109 | 0.52 | 0.53 | 0.54 | 0.58 | 0.57 | 0.59 | 0.55 | 0.55 | 0.54 | 0.59 | 0.58 | 0.59 | 0.58 | 0.59 | 0.59 |
| 109.5 | 0.54 | 0.55 | 0.55 | 0.58 | 0.56 | 0.58 | 0.55 | 0.54 | 0.54 | 0.59 | 0.56 | 0.59 | 0.58 | 0.57 | 0.59 |
| 110 | 0.54 | 0.54 | 0.53 | 0.56 | 0.56 | 0.57 | 0.53 | 0.56 | 0.54 | 0.6 | 0.58 | 0.6 | 0.58 | 0.57 | 0.59 |
| 110.5 | 0.54 | 0.54 | 0.54 | 0.57 | 0.56 | 0.55 | 0.61 | 0.54 | 0.56 | 0.59 | 0.6 | 0.59 | 0.58 | 0.58 | 0.59 |
| 111 | 0.54 | 0.53 | 0.52 | 0.58 | 0.55 | 0.56 | 0.58 | 0.53 | 0.54 | 0.59 | 0.59 | 0.6 | 0.59 | 0.58 | 0.58 |
| 111.5 | 0.53 | 0.52 | 0.53 | 0.58 | 0.57 | 0.56 | 0.58 | 0.52 | 0.51 | 0.6 | 0.56 | 0.59 | 0.6 | 0.57 | 0.58 |
| 112 | 0.54 | 0.54 | 0.52 | 0.55 | 0.56 | 0.55 | 0.54 | 0.56 | 0.52 | 0.58 | 0.56 | 0.59 | 0.58 | 0.58 | 0.57 |
| 112.5 | 0.53 | 0.54 | 0.53 | 0.57 | 0.55 | 0.55 | 0.53 | 0.54 | 0.5 | 0.6 | 0.57 | 0.59 | 0.58 | 0.57 | 0.57 |
| 113 | 0.56 | 0.55 | 0.51 | 0.57 | 0.55 | 0.54 | 0.59 | 0.53 | 0.47 | 0.58 | 0.57 | 0.57 | 0.58 | 0.57 | 0.57 |
| 113.5 | 0.53 | 0.53 | 0.49 | 0.57 | 0.57 | 0.54 | 0.62 | 0.52 | 0.49 | 0.57 | 0.58 | 0.58 | 0.58 | 0.58 | 0.56 |
| 114 | 0.54 | 0.53 | 0.48 | 0.58 | 0.56 | 0.52 | 0.58 | 0.52 | 0.48 | 0.59 | 0.59 | 0.57 | 0.58 | 0.58 | 0.56 |
| 114.5 | 0.53 | 0.52 | 0.49 | 0.58 | 0.56 | 0.51 | 0.52 | 0.53 | 0.46 | 0.57 | 0.57 | 0.57 | 0.58 | 0.57 | 0.56 |
| 115 | 0.55 | 0.52 | 0.46 | 0.57 | 0.54 | 0.49 | 0.57 | 0.51 | 0.4 | 0.56 | 0.57 | 0.56 | 0.58 | 0.58 | 0.55 |
| 115.5 | 0.54 | 0.5 | 0.44 | 0.56 | 0.55 | 0.48 | 0.57 | 0.49 | 0.38 | 0.57 | 0.58 | 0.56 | 0.57 | 0.57 | 0.55 |
| 116 | 0.52 | 0.5 | 0.41 | 0.58 | 0.53 | 0.45 | 0.53 | 0.45 | 0.32 | 0.58 | 0.57 | 0.56 | 0.58 | 0.56 | 0.53 |
| 116.5 | 0.54 | 0.47 | 0.35 | 0.57 | 0.53 | 0.41 | 0.53 | 0.45 | 0.27 | 0.56 | 0.57 | 0.54 | 0.57 | 0.57 | 0.51 |
| 117 | 0.53 | 0.48 | 0.29 | 0.57 | 0.51 | 0.37 | 0.54 | 0.4 | 0.18 | 0.59 | 0.58 | 0.53 | 0.59 | 0.56 | 0.5 |
| 117.5 | 0.52 | 0.42 | 0.22 | 0.57 | 0.49 | 0.3 | 0.54 | 0.3 | 0.06 | 0.58 | 0.56 | 0.54 | 0.57 | 0.55 | 0.48 |
| 118 | 0.52 | 0.32 | 0.12 | 0.57 | 0.45 | 0.22 | 0.5 | 0.12 | 0.01 | 0.58 | 0.56 | 0.52 | 0.58 | 0.55 | 0.45 |
| 118.5 | 0.41 | 0.16 | 0.03 | 0.56 | 0.37 | 0.12 | 0.25 | 0.01 | 0 | 0.59 | 0.56 | 0.5 | 0.58 | 0.54 | 0.42 |
| 119 | 0.22 | 0.02 | 0 | 0.51 | 0.25 | 0.05 | 0.04 | 0.01 | 0.01 | 0.57 | 0.55 | 0.49 | 0.58 | 0.52 | 0.38 |
| 119.5 | 0.08 | 0 | 0 | 0.41 | 0.08 | 0.01 | 0.36 | 0.04 | 0.02 | 0.56 | 0.53 | 0.47 | 0.57 | 0.5 | 0.32 |
| 120 | 0.46 | 0.14 | 0.02 | 0.03 | 0 | 0.01 | 0.53 | 0.2 | 0.04 | 0.56 | 0.53 | 0.43 | 0.58 | 0.47 | 0.25 |
| 120.5 | 0.52 | 0.31 | 0.09 | 0.25 | 0.02 | 0 | 0.54 | 0.33 | 0.14 | 0.57 | 0.53 | 0.38 | 0.56 | 0.41 | 0.16 |
| 121 | 0.52 | 0.4 | 0.19 | 0.48 | 0.15 | 0.02 | 0.54 | 0.4 | 0.22 | 0.56 | 0.5 | 0.32 | 0.54 | 0.3 | 0.08 |
| 121.5 | 0.53 | 0.45 | 0.28 | 0.54 | 0.31 | 0.08 | 0.5 | 0.45 | 0.28 | 0.58 | 0.47 | 0.25 | 0.49 | 0.14 | 0.01 |
| 122 | 0.52 | 0.47 | 0.34 | 0.54 | 0.42 | 0.17 | 0.49 | 0.47 | 0.33 | 0.55 | 0.4 | 0.16 | 0.18 | 0.01 | 0 |
| 122.5 | 0.54 | 0.51 | 0.4 | 0.57 | 0.47 | 0.25 | 0.55 | 0.48 | 0.39 | 0.55 | 0.3 | 0.07 | 0.1 | 0.01 | 0 |
| 123 | 0.54 | 0.51 | 0.42 | 0.55 | 0.49 | 0.33 | 0.59 | 0.51 | 0.41 | 0.48 | 0.13 | 0.02 | 0.45 | 0.09 | 0.01 |
| 123.5 | 0.53 | 0.53 | 0.45 | 0.58 | 0.52 | 0.38 | 0.54 | 0.53 | 0.46 | 0.18 | 0.01 | 0 | 0.53 | 0.27 | 0.06 |
| 124 | 0.54 | 0.52 | 0.47 | 0.56 | 0.54 | 0.41 | 0.55 | 0.51 | 0.47 | 0.07 | 0.01 | 0 | 0.55 | 0.37 | 0.14 |
| 124.5 | 0.53 | 0.53 | 0.49 | 0.57 | 0.54 | 0.46 | 0.6 | 0.53 | 0.47 | 0.44 | 0.09 | 0.01 | 0.56 | 0.44 | 0.23 |
| 125 | 0.54 | 0.52 | 0.5 | 0.57 | 0.55 | 0.48 | 0.54 | 0.51 | 0.53 | 0.55 | 0.28 | 0.07 | 0.58 | 0.49 | 0.31 |
| 125.5 | 0.52 | 0.53 | 0.5 | 0.57 | 0.57 | 0.5 | 0.53 | 0.54 | 0.52 | 0.55 | 0.39 | 0.15 | 0.56 | 0.51 | 0.37 |
| 126 | 0.54 | 0.52 | 0.52 | 0.58 | 0.56 | 0.51 | 0.58 | 0.53 | 0.55 | 0.54 | 0.47 | 0.24 | 0.58 | 0.53 | 0.41 |
| 126.5 | 0.54 | 0.55 | 0.52 | 0.57 | 0.56 | 0.53 | 0.56 | 0.5 | 0.53 | 0.58 | 0.49 | 0.31 | 0.57 | 0.54 | 0.45 |
| 127 | 0.54 | 0.55 | 0.53 | 0.57 | 0.56 | 0.54 | 0.54 | 0.54 | 0.52 | 0.59 | 0.52 | 0.38 | 0.58 | 0.55 | 0.48 |
| 127.5 | 0.53 | 0.53 | 0.55 | 0.58 | 0.55 | 0.54 | 0.55 | 0.52 | 0.56 | 0.57 | 0.54 | 0.43 | 0.59 | 0.56 | 0.5 |
| 128 | 0.55 | 0.54 | 0.55 | 0.57 | 0.57 | 0.55 | 0.53 | 0.52 | 0.55 | 0.57 | 0.53 | 0.44 | 0.59 | 0.57 | 0.52 |
| 128.5 | 0.54 | 0.53 | 0.53 | 0.58 | 0.56 | 0.55 | 0.57 | 0.54 | 0.57 | 0.57 | 0.56 | 0.48 | 0.58 | 0.56 | 0.53 |
| 129 | 0.57 | 0.54 | 0.53 | 0.57 | 0.58 | 0.55 | 0.55 | 0.54 | 0.58 | 0.59 | 0.56 | 0.5 | 0.59 | 0.58 | 0.53 |
| 129.5 | 0.55 | 0.52 | 0.55 | 0.57 | 0.57 | 0.57 | 0.53 | 0.55 | 0.55 | 0.58 | 0.56 | 0.51 | 0.59 | 0.56 | 0.54 |
| 130 | 0.54 | 0.54 | 0.54 | 0.57 | 0.56 | 0.58 | 0.6 | 0.5 | 0.57 | 0.56 | 0.54 | 0.52 | 0.57 | 0.57 | 0.55 |
| 130.5 | 0.54 | 0.55 | 0.55 | 0.57 | 0.56 | 0.57 | 0.59 | 0.54 | 0.55 | 0.59 | 0.58 | 0.55 | 0.58 | 0.58 | 0.56 |
| 131 | 0.53 | 0.53 | 0.56 | 0.57 | 0.56 | 0.56 | 0.6 | 0.53 | 0.55 | 0.59 | 0.57 | 0.56 | 0.58 | 0.56 | 0.57 |
| 131.5 | 0.55 | 0.54 | 0.54 | 0.57 | 0.58 | 0.57 | 0.54 | 0.55 | 0.56 | 0.57 | 0.58 | 0.56 | 0.58 | 0.58 | 0.56 |
| 132 | 0.54 | 0.52 | 0.54 | 0.58 | 0.58 | 0.57 | 0.55 | 0.53 | 0.56 | 0.58 | 0.57 | 0.56 | 0.58 | 0.58 | 0.58 |
| 132.5 | 0.55 | 0.53 | 0.54 | 0.57 | 0.58 | 0.58 | 0.58 | 0.55 | 0.57 | 0.57 | 0.57 | 0.56 | 0.58 | 0.58 | 0.58 |
| 133 | 0.55 | 0.54 | 0.55 | 0.57 | 0.57 | 0.59 | 0.53 | 0.54 | 0.55 | 0.59 | 0.56 | 0.56 | 0.59 | 0.57 | 0.57 |

| 10HzG291 | 30HzG291 | 60HzG291 | 10HzG293 | 30HzG293 | 60HzG293 | 10HzE294 | 30HzE294 | 60HzE294 | 10HzN295 | 30HzN295 | 60HzN295 | 10HzV296 | 30HzV296 | 60HzV296 |
| --- | --- | --- | --- | --- | --- | --- | --- | --- | --- | --- | --- | --- | --- | --- |
| 0.61 | 0.58 | 0.56 | 0.59 | 0.51 | 0.34 | 0.62 | 0.6 | 0.6 | 0.57 | 0.59 | 0.56 | 0.68 | 0.61 | 0.64 |
| 0.63 | 0.59 | 0.54 | 0.57 | 0.47 | 0.28 | 0.61 | 0.6 | 0.6 | 0.57 | 0.58 | 0.57 | 0.67 | 0.63 | 0.64 |
| 0.6 | 0.56 | 0.53 | 0.57 | 0.42 | 0.19 | 0.6 | 0.59 | 0.59 | 0.58 | 0.57 | 0.56 | 0.68 | 0.66 | 0.64 |
| 0.6 | 0.56 | 0.52 | 0.55 | 0.35 | 0.1 | 0.6 | 0.61 | 0.59 | 0.57 | 0.58 | 0.56 | 0.64 | 0.65 | 0.65 |
| 0.6 | 0.56 | 0.5 | 0.52 | 0.19 | 0.03 | 0.61 | 0.6 | 0.58 | 0.57 | 0.58 | 0.56 | 0.68 | 0.63 | 0.64 |
| 0.6 | 0.56 | 0.49 | 0.33 | 0.03 | 0.01 | 0.62 | 0.6 | 0.59 | 0.59 | 0.59 | 0.56 | 0.67 | 0.67 | 0.64 |
| 0.61 | 0.55 | 0.46 | 0.01 | 0.01 | 0.01 | 0.61 | 0.59 | 0.58 | 0.59 | 0.58 | 0.56 | 0.66 | 0.65 | 0.64 |
| 0.6 | 0.51 | 0.43 | 0.42 | 0.07 | 0.01 | 0.61 | 0.59 | 0.58 | 0.57 | 0.58 | 0.56 | 0.66 | 0.64 | 0.64 |
| 0.6 | 0.51 | 0.38 | 0.53 | 0.23 | 0.05 | 0.61 | 0.6 | 0.56 | 0.58 | 0.57 | 0.55 | 0.66 | 0.63 | 0.64 |
| 0.61 | 0.5 | 0.32 | 0.58 | 0.37 | 0.12 | 0.6 | 0.59 | 0.56 | 0.58 | 0.57 | 0.56 | 0.66 | 0.65 | 0.65 |
| 0.58 | 0.47 | 0.25 | 0.57 | 0.43 | 0.21 | 0.61 | 0.59 | 0.55 | 0.56 | 0.57 | 0.55 | 0.63 | 0.65 | 0.63 |
| 0.58 | 0.4 | 0.15 | 0.58 | 0.49 | 0.28 | 0.61 | 0.6 | 0.53 | 0.56 | 0.57 | 0.55 | 0.67 | 0.64 | 0.62 |
| 0.58 | 0.28 | 0.06 | 0.58 | 0.52 | 0.34 | 0.61 | 0.59 | 0.53 | 0.57 | 0.57 | 0.54 | 0.68 | 0.62 | 0.63 |
| 0.49 | 0.1 | 0.02 | 0.58 | 0.51 | 0.38 | 0.61 | 0.57 | 0.51 | 0.58 | 0.57 | 0.54 | 0.63 | 0.64 | 0.63 |
| 0.08 | 0.01 | 0 | 0.57 | 0.5 | 0.41 | 0.61 | 0.56 | 0.48 | 0.58 | 0.58 | 0.54 | 0.65 | 0.64 | 0.62 |
| 0.23 | 0.01 | 0.01 | 0.58 | 0.53 | 0.43 | 0.6 | 0.56 | 0.45 | 0.56 | 0.56 | 0.53 | 0.65 | 0.64 | 0.63 |
| 0.5 | 0.15 | 0.02 | 0.59 | 0.54 | 0.46 | 0.61 | 0.55 | 0.41 | 0.58 | 0.55 | 0.52 | 0.67 | 0.63 | 0.62 |
| 0.55 | 0.3 | 0.08 | 0.58 | 0.57 | 0.49 | 0.59 | 0.54 | 0.37 | 0.58 | 0.56 | 0.51 | 0.63 | 0.64 | 0.62 |
| 0.59 | 0.43 | 0.18 | 0.6 | 0.58 | 0.51 | 0.6 | 0.51 | 0.3 | 0.57 | 0.55 | 0.5 | 0.65 | 0.64 | 0.61 |
| 0.61 | 0.47 | 0.26 | 0.59 | 0.57 | 0.52 | 0.6 | 0.47 | 0.22 | 0.57 | 0.56 | 0.48 | 0.65 | 0.63 | 0.6 |
| 0.6 | 0.52 | 0.33 | 0.58 | 0.58 | 0.52 | 0.58 | 0.39 | 0.13 | 0.58 | 0.55 | 0.47 | 0.65 | 0.62 | 0.59 |
| 0.6 | 0.52 | 0.39 | 0.58 | 0.56 | 0.54 | 0.56 | 0.26 | 0.05 | 0.57 | 0.55 | 0.45 | 0.66 | 0.65 | 0.59 |
| 0.61 | 0.53 | 0.44 | 0.59 | 0.57 | 0.55 | 0.46 | 0.07 | 0.01 | 0.56 | 0.53 | 0.42 | 0.64 | 0.64 | 0.57 |
| 0.61 | 0.55 | 0.45 | 0.57 | 0.57 | 0.55 | 0.03 | 0 | 0 | 0.57 | 0.52 | 0.38 | 0.67 | 0.62 | 0.56 |
| 0.6 | 0.55 | 0.5 | 0.61 | 0.57 | 0.55 | 0.24 | 0.01 | 0 | 0.56 | 0.51 | 0.32 | 0.65 | 0.63 | 0.55 |
| 0.61 | 0.57 | 0.51 | 0.59 | 0.57 | 0.56 | 0.51 | 0.14 | 0.02 | 0.56 | 0.46 | 0.24 | 0.64 | 0.6 | 0.5 |
| 0.63 | 0.56 | 0.5 | 0.59 | 0.58 | 0.56 | 0.57 | 0.31 | 0.07 | 0.56 | 0.41 | 0.15 | 0.64 | 0.61 | 0.47 |
| 0.62 | 0.57 | 0.53 | 0.61 | 0.58 | 0.58 | 0.58 | 0.42 | 0.16 | 0.54 | 0.29 | 0.07 | 0.64 | 0.58 | 0.42 |
| 0.62 | 0.57 | 0.54 | 0.58 | 0.58 | 0.57 | 0.59 | 0.48 | 0.25 | 0.45 | 0.1 | 0.01 | 0.65 | 0.54 | 0.34 |
| 0.65 | 0.58 | 0.55 | 0.59 | 0.58 | 0.57 | 0.6 | 0.51 | 0.32 | 0.08 | 0 | 0 | 0.64 | 0.5 | 0.25 |
| 0.63 | 0.57 | 0.56 | 0.59 | 0.56 | 0.57 | 0.6 | 0.53 | 0.37 | 0.24 | 0.02 | 0 | 0.61 | 0.39 | 0.13 |
| 0.61 | 0.57 | 0.56 | 0.58 | 0.58 | 0.58 | 0.62 | 0.54 | 0.42 | 0.48 | 0.16 | 0.02 | 0.56 | 0.18 | 0.03 |
| 0.6 | 0.58 | 0.56 | 0.6 | 0.57 | 0.58 | 0.61 | 0.56 | 0.45 | 0.54 | 0.32 | 0.09 | 0.18 | 0.02 | -0.01 |
| 0.61 | 0.56 | 0.58 | 0.61 | 0.57 | 0.58 | 0.6 | 0.56 | 0.46 | 0.55 | 0.38 | 0.18 | 0.25 | 0.01 | 0 |
| 0.62 | 0.6 | 0.57 | 0.59 | 0.58 | 0.57 | 0.6 | 0.56 | 0.48 | 0.55 | 0.45 | 0.26 | 0.56 | 0.2 | 0.03 |
| 0.61 | 0.58 | 0.57 | 0.59 | 0.57 | 0.57 | 0.6 | 0.55 | 0.48 | 0.56 | 0.5 | 0.33 | 0.6 | 0.38 | 0.14 |
| 0.6 | 0.59 | 0.57 | 0.61 | 0.57 | 0.58 | 0.61 | 0.56 | 0.5 | 0.58 | 0.52 | 0.38 | 0.62 | 0.48 | 0.23 |
| 0.63 | 0.59 | 0.58 | 0.58 | 0.58 | 0.57 | 0.6 | 0.58 | 0.52 | 0.57 | 0.54 | 0.42 | 0.65 | 0.52 | 0.34 |
| 0.62 | 0.59 | 0.58 | 0.6 | 0.58 | 0.59 | 0.6 | 0.59 | 0.54 | 0.56 | 0.56 | 0.45 | 0.64 | 0.56 | 0.41 |
| 0.63 | 0.57 | 0.58 | 0.6 | 0.57 | 0.58 | 0.61 | 0.6 | 0.56 | 0.57 | 0.55 | 0.47 | 0.65 | 0.6 | 0.47 |
| 0.62 | 0.59 | 0.59 | 0.6 | 0.58 | 0.58 | 0.61 | 0.6 | 0.57 | 0.58 | 0.56 | 0.49 | 0.64 | 0.62 | 0.51 |
| 0.61 | 0.59 | 0.6 | 0.6 | 0.58 | 0.58 | 0.61 | 0.59 | 0.58 | 0.57 | 0.56 | 0.51 | 0.67 | 0.62 | 0.53 |
| 0.6 | 0.59 | 0.59 | 0.57 | 0.57 | 0.59 | 0.6 | 0.6 | 0.58 | 0.57 | 0.57 | 0.51 | 0.64 | 0.62 | 0.55 |
| 0.6 | 0.59 | 0.59 | 0.59 | 0.58 | 0.59 | 0.61 | 0.6 | 0.58 | 0.57 | 0.56 | 0.52 | 0.66 | 0.63 | 0.58 |
| 0.61 | 0.6 | 0.59 | 0.59 | 0.59 | 0.58 | 0.61 | 0.6 | 0.59 | 0.57 | 0.57 | 0.53 | 0.66 | 0.62 | 0.58 |
| 0.63 | 0.58 | 0.59 | 0.59 | 0.59 | 0.58 | 0.61 | 0.61 | 0.6 | 0.57 | 0.57 | 0.53 | 0.66 | 0.63 | 0.59 |
| 0.62 | 0.57 | 0.59 | 0.59 | 0.58 | 0.59 | 0.6 | 0.6 | 0.59 | 0.57 | 0.58 | 0.54 | 0.65 | 0.64 | 0.61 |
| 0.62 | 0.59 | 0.59 | 0.59 | 0.59 | 0.58 | 0.61 | 0.6 | 0.6 | 0.57 | 0.58 | 0.55 | 0.66 | 0.64 | 0.61 |
| 0.6 | 0.58 | 0.59 | 0.6 | 0.58 | 0.57 | 0.61 | 0.6 | 0.6 | 0.57 | 0.58 | 0.55 | 0.65 | 0.62 | 0.62 |
| 0.6 | 0.58 | 0.59 | 0.6 | 0.57 | 0.56 | 0.61 | 0.6 | 0.6 | 0.57 | 0.57 | 0.55 | 0.64 | 0.64 | 0.62 |
| 0.62 | 0.58 | 0.59 | 0.61 | 0.58 | 0.58 | 0.62 | 0.6 | 0.6 | 0.58 | 0.58 | 0.55 | 0.65 | 0.65 | 0.62 |
| 0.62 | 0.58 | 0.59 | 0.58 | 0.58 | 0.58 | 0.61 | 0.61 | 0.61 | 0.57 | 0.58 | 0.55 | 0.64 | 0.65 | 0.62 |
| 0.6 | 0.59 | 0.59 | 0.59 | 0.58 | 0.59 | 0.61 | 0.6 | 0.6 | 0.57 | 0.59 | 0.55 | 0.65 | 0.65 | 0.62 |
| 0.59 | 0.58 | 0.59 | 0.58 | 0.59 | 0.58 | 0.6 | 0.61 | 0.6 | 0.57 | 0.58 | 0.56 | 0.65 | 0.65 | 0.62 |
| 0.61 | 0.58 | 0.58 | 0.6 | 0.59 | 0.59 | 0.6 | 0.6 | 0.61 | 0.56 | 0.58 | 0.56 | 0.68 | 0.64 | 0.62 |
| 0.62 | 0.59 | 0.59 | 0.6 | 0.58 | 0.59 | 0.61 | 0.6 | 0.61 | 0.58 | 0.58 | 0.56 | 0.67 | 0.65 | 0.64 |
| 0.6 | 0.58 | 0.58 | 0.57 | 0.58 | 0.59 | 0.61 | 0.61 | 0.61 | 0.58 | 0.57 | 0.56 | 0.65 | 0.65 | 0.63 |
| 0.62 | 0.57 | 0.58 | 0.6 | 0.58 | 0.59 | 0.61 | 0.6 | 0.61 | 0.58 | 0.58 | 0.57 | 0.65 | 0.64 | 0.64 |
| 0.62 | 0.59 | 0.58 | 0.59 | 0.57 | 0.59 | 0.62 | 0.6 | 0.61 | 0.58 | 0.58 | 0.56 | 0.65 | 0.64 | 0.63 |
| 0.64 | 0.59 | 0.6 | 0.58 | 0.59 | 0.58 | 0.61 | 0.6 | 0.61 | 0.56 | 0.58 | 0.56 | 0.66 | 0.66 | 0.65 |
| 0.62 | 0.6 | 0.59 | 0.59 | 0.58 | 0.59 | 0.62 | 0.6 | 0.61 | 0.57 | 0.58 | 0.56 | 0.64 | 0.65 | 0.64 |
| 0.63 | 0.57 | 0.6 | 0.59 | 0.58 | 0.59 | 0.62 | 0.6 | 0.61 | 0.57 | 0.58 | 0.57 | 0.66 | 0.64 | 0.63 |
| 0.61 | 0.58 | 0.6 | 0.58 | 0.58 | 0.59 | 0.61 | 0.61 | 0.62 | 0.57 | 0.58 | 0.57 | 0.64 | 0.65 | 0.64 |

| 10HzT297 | 30HzT297 | 60HzT297 | 10Hz I298 | 30Hz I298 | 60Hz I298 | 10HzE299 | 30HzE299 | 60HzE299 | 10Hz S300 | 30Hz S300 | 60Hz S300 | 10HzV301 | 30HzV301 | 60HzV301 |
| --- | --- | --- | --- | --- | --- | --- | --- | --- | --- | --- | --- | --- | --- | --- |
| 0.6 | 0.58 | 0.58 | 0.6 | 0.58 | 0.59 | 0.58 | 0.58 | 0.58 | 0.6 | 0.59 | 0.6 | 0.57 | 0.59 | 0.57 |
| 0.61 | 0.59 | 0.58 | 0.59 | 0.59 | 0.6 | 0.58 | 0.58 | 0.58 | 0.6 | 0.59 | 0.59 | 0.57 | 0.54 | 0.59 |
| 0.6 | 0.59 | 0.57 | 0.58 | 0.59 | 0.6 | 0.59 | 0.58 | 0.58 | 0.61 | 0.59 | 0.59 | 0.61 | 0.58 | 0.56 |
| 0.61 | 0.59 | 0.57 | 0.59 | 0.57 | 0.59 | 0.58 | 0.58 | 0.58 | 0.6 | 0.6 | 0.59 | 0.59 | 0.55 | 0.55 |
| 0.6 | 0.58 | 0.57 | 0.57 | 0.59 | 0.57 | 0.59 | 0.58 | 0.57 | 0.6 | 0.59 | 0.59 | 0.55 | 0.58 | 0.57 |
| 0.59 | 0.58 | 0.55 | 0.59 | 0.58 | 0.59 | 0.59 | 0.58 | 0.58 | 0.6 | 0.59 | 0.58 | 0.61 | 0.6 | 0.58 |
| 0.6 | 0.58 | 0.55 | 0.59 | 0.58 | 0.57 | 0.58 | 0.58 | 0.58 | 0.6 | 0.58 | 0.59 | 0.59 | 0.58 | 0.58 |
| 0.6 | 0.58 | 0.54 | 0.57 | 0.59 | 0.58 | 0.57 | 0.58 | 0.57 | 0.6 | 0.59 | 0.58 | 0.59 | 0.57 | 0.58 |
| 0.59 | 0.58 | 0.52 | 0.57 | 0.59 | 0.57 | 0.59 | 0.58 | 0.57 | 0.6 | 0.59 | 0.58 | 0.56 | 0.58 | 0.58 |
| 0.6 | 0.57 | 0.5 | 0.59 | 0.57 | 0.59 | 0.59 | 0.58 | 0.58 | 0.6 | 0.59 | 0.58 | 0.58 | 0.58 | 0.57 |
| 0.6 | 0.56 | 0.49 | 0.58 | 0.58 | 0.58 | 0.58 | 0.57 | 0.57 | 0.6 | 0.59 | 0.58 | 0.59 | 0.6 | 0.56 |
| 0.6 | 0.56 | 0.46 | 0.57 | 0.59 | 0.58 | 0.6 | 0.57 | 0.57 | 0.6 | 0.59 | 0.57 | 0.56 | 0.58 | 0.57 |
| 0.59 | 0.55 | 0.42 | 0.59 | 0.59 | 0.58 | 0.59 | 0.59 | 0.56 | 0.6 | 0.58 | 0.56 | 0.61 | 0.59 | 0.57 |
| 0.6 | 0.52 | 0.38 | 0.58 | 0.57 | 0.59 | 0.58 | 0.57 | 0.57 | 0.6 | 0.58 | 0.55 | 0.61 | 0.59 | 0.55 |
| 0.58 | 0.5 | 0.31 | 0.59 | 0.57 | 0.58 | 0.59 | 0.58 | 0.56 | 0.6 | 0.57 | 0.55 | 0.6 | 0.56 | 0.56 |
| 0.6 | 0.46 | 0.23 | 0.59 | 0.58 | 0.59 | 0.58 | 0.58 | 0.55 | 0.61 | 0.58 | 0.54 | 0.59 | 0.58 | 0.56 |
| 0.59 | 0.38 | 0.13 | 0.61 | 0.6 | 0.58 | 0.58 | 0.57 | 0.55 | 0.6 | 0.57 | 0.52 | 0.58 | 0.58 | 0.58 |
| 0.54 | 0.24 | 0.04 | 0.61 | 0.59 | 0.57 | 0.58 | 0.57 | 0.54 | 0.6 | 0.56 | 0.51 | 0.56 | 0.56 | 0.57 |
| 0.41 | 0.06 | 0.01 | 0.59 | 0.58 | 0.58 | 0.58 | 0.57 | 0.53 | 0.6 | 0.56 | 0.49 | 0.59 | 0.59 | 0.55 |
| 0.04 | 0 | 0 | 0.57 | 0.59 | 0.57 | 0.58 | 0.57 | 0.53 | 0.6 | 0.55 | 0.47 | 0.59 | 0.59 | 0.56 |
| 0.38 | 0.04 | 0 | 0.58 | 0.6 | 0.58 | 0.6 | 0.57 | 0.52 | 0.6 | 0.55 | 0.43 | 0.61 | 0.61 | 0.57 |
| 0.55 | 0.22 | 0.04 | 0.57 | 0.58 | 0.55 | 0.58 | 0.56 | 0.51 | 0.6 | 0.53 | 0.39 | 0.56 | 0.59 | 0.57 |
| 0.57 | 0.36 | 0.11 | 0.61 | 0.59 | 0.57 | 0.58 | 0.56 | 0.48 | 0.59 | 0.51 | 0.34 | 0.59 | 0.56 | 0.55 |
| 0.58 | 0.45 | 0.21 | 0.57 | 0.59 | 0.56 | 0.59 | 0.56 | 0.47 | 0.59 | 0.48 | 0.26 | 0.58 | 0.57 | 0.56 |
| 0.59 | 0.5 | 0.3 | 0.57 | 0.59 | 0.55 | 0.59 | 0.55 | 0.43 | 0.59 | 0.41 | 0.17 | 0.59 | 0.58 | 0.56 |
| 0.6 | 0.52 | 0.36 | 0.6 | 0.58 | 0.55 | 0.58 | 0.52 | 0.4 | 0.56 | 0.3 | 0.08 | 0.6 | 0.58 | 0.55 |
| 0.59 | 0.55 | 0.4 | 0.59 | 0.59 | 0.55 | 0.57 | 0.51 | 0.36 | 0.49 | 0.13 | 0.02 | 0.57 | 0.56 | 0.56 |
| 0.6 | 0.55 | 0.43 | 0.6 | 0.59 | 0.53 | 0.57 | 0.48 | 0.29 | 0.19 | 0.01 | 0 | 0.58 | 0.59 | 0.56 |
| 0.61 | 0.56 | 0.46 | 0.57 | 0.54 | 0.52 | 0.57 | 0.44 | 0.21 | 0.11 | 0.01 | 0 | 0.56 | 0.57 | 0.54 |
| 0.58 | 0.55 | 0.47 | 0.57 | 0.58 | 0.5 | 0.56 | 0.38 | 0.12 | 0.48 | 0.11 | 0.02 | 0.61 | 0.56 | 0.54 |
| 0.59 | 0.55 | 0.49 | 0.62 | 0.57 | 0.49 | 0.53 | 0.24 | 0.05 | 0.56 | 0.28 | 0.07 | 0.59 | 0.54 | 0.55 |
| 0.59 | 0.54 | 0.5 | 0.59 | 0.57 | 0.46 | 0.42 | 0.07 | 0.01 | 0.58 | 0.39 | 0.15 | 0.58 | 0.58 | 0.53 |
| 0.59 | 0.55 | 0.5 | 0.59 | 0.55 | 0.45 | 0.02 | 0.01 | 0 | 0.59 | 0.46 | 0.24 | 0.6 | 0.6 | 0.53 |
| 0.59 | 0.57 | 0.52 | 0.59 | 0.55 | 0.4 | 0.29 | 0.01 | 0 | 0.6 | 0.5 | 0.31 | 0.59 | 0.56 | 0.53 |
| 0.61 | 0.58 | 0.54 | 0.59 | 0.52 | 0.36 | 0.5 | 0.17 | 0.02 | 0.59 | 0.53 | 0.37 | 0.54 | 0.56 | 0.51 |
| 0.6 | 0.59 | 0.55 | 0.58 | 0.5 | 0.3 | 0.56 | 0.32 | 0.09 | 0.59 | 0.54 | 0.42 | 0.57 | 0.55 | 0.5 |
| 0.6 | 0.57 | 0.57 | 0.56 | 0.45 | 0.22 | 0.56 | 0.42 | 0.17 | 0.6 | 0.54 | 0.44 | 0.61 | 0.55 | 0.45 |
| 0.6 | 0.59 | 0.56 | 0.56 | 0.37 | 0.14 | 0.57 | 0.46 | 0.25 | 0.59 | 0.54 | 0.46 | 0.58 | 0.5 | 0.45 |
| 0.59 | 0.59 | 0.58 | 0.55 | 0.25 | 0.04 | 0.57 | 0.5 | 0.31 | 0.59 | 0.55 | 0.48 | 0.61 | 0.52 | 0.41 |
| 0.6 | 0.59 | 0.58 | 0.44 | 0.09 | 0.01 | 0.57 | 0.51 | 0.36 | 0.6 | 0.56 | 0.51 | 0.6 | 0.51 | 0.34 |
| 0.6 | 0.59 | 0.58 | 0.02 | 0 | 0.01 | 0.58 | 0.53 | 0.39 | 0.6 | 0.58 | 0.52 | 0.58 | 0.5 | 0.3 |
| 0.6 | 0.58 | 0.59 | 0.27 | 0 | 0 | 0.57 | 0.52 | 0.42 | 0.6 | 0.58 | 0.54 | 0.56 | 0.46 | 0.21 |
| 0.59 | 0.59 | 0.59 | 0.5 | 0.16 | 0.03 | 0.58 | 0.52 | 0.44 | 0.6 | 0.58 | 0.55 | 0.57 | 0.35 | 0.14 |
| 0.59 | 0.58 | 0.6 | 0.53 | 0.33 | 0.09 | 0.57 | 0.52 | 0.46 | 0.6 | 0.58 | 0.56 | 0.56 | 0.27 | 0.06 |
| 0.58 | 0.59 | 0.59 | 0.59 | 0.43 | 0.18 | 0.57 | 0.55 | 0.49 | 0.6 | 0.58 | 0.56 | 0.46 | 0.1 | 0 |
| 0.61 | 0.59 | 0.59 | 0.59 | 0.47 | 0.26 | 0.59 | 0.56 | 0.51 | 0.6 | 0.58 | 0.57 | 0.14 | 0.01 | -0.01 |
| 0.6 | 0.6 | 0.6 | 0.59 | 0.5 | 0.33 | 0.59 | 0.56 | 0.52 | 0.59 | 0.58 | 0.57 | 0.13 | 0 | -0.01 |
| 0.59 | 0.6 | 0.59 | 0.6 | 0.53 | 0.38 | 0.59 | 0.57 | 0.53 | 0.61 | 0.59 | 0.58 | 0.44 | 0.09 | 0 |
| 0.61 | 0.6 | 0.6 | 0.58 | 0.56 | 0.43 | 0.58 | 0.57 | 0.54 | 0.6 | 0.59 | 0.58 | 0.54 | 0.28 | 0.06 |
| 0.61 | 0.6 | 0.61 | 0.57 | 0.57 | 0.45 | 0.59 | 0.57 | 0.54 | 0.6 | 0.59 | 0.59 | 0.59 | 0.39 | 0.14 |
| 0.59 | 0.59 | 0.6 | 0.59 | 0.56 | 0.48 | 0.59 | 0.57 | 0.55 | 0.6 | 0.58 | 0.59 | 0.59 | 0.46 | 0.24 |
| 0.6 | 0.59 | 0.6 | 0.59 | 0.57 | 0.5 | 0.59 | 0.58 | 0.56 | 0.61 | 0.59 | 0.59 | 0.59 | 0.51 | 0.29 |
| 0.6 | 0.6 | 0.6 | 0.58 | 0.56 | 0.51 | 0.58 | 0.57 | 0.56 | 0.6 | 0.58 | 0.59 | 0.58 | 0.51 | 0.35 |
| 0.6 | 0.6 | 0.6 | 0.57 | 0.57 | 0.53 | 0.58 | 0.58 | 0.57 | 0.61 | 0.59 | 0.6 | 0.58 | 0.53 | 0.4 |
| 0.61 | 0.6 | 0.6 | 0.58 | 0.57 | 0.53 | 0.58 | 0.57 | 0.56 | 0.61 | 0.59 | 0.59 | 0.58 | 0.53 | 0.44 |
| 0.61 | 0.6 | 0.61 | 0.59 | 0.58 | 0.54 | 0.59 | 0.58 | 0.58 | 0.6 | 0.59 | 0.6 | 0.6 | 0.54 | 0.47 |
| 0.6 | 0.59 | 0.6 | 0.57 | 0.57 | 0.54 | 0.59 | 0.56 | 0.57 | 0.6 | 0.59 | 0.59 | 0.6 | 0.56 | 0.48 |
| 0.59 | 0.59 | 0.61 | 0.57 | 0.57 | 0.56 | 0.58 | 0.58 | 0.57 | 0.6 | 0.59 | 0.6 | 0.6 | 0.58 | 0.49 |
| 0.59 | 0.6 | 0.6 | 0.6 | 0.57 | 0.57 | 0.58 | 0.58 | 0.57 | 0.6 | 0.59 | 0.6 | 0.57 | 0.58 | 0.53 |
| 0.59 | 0.6 | 0.6 | 0.6 | 0.59 | 0.56 | 0.59 | 0.58 | 0.58 | 0.6 | 0.6 | 0.6 | 0.56 | 0.56 | 0.52 |
| 0.6 | 0.6 | 0.6 | 0.58 | 0.58 | 0.58 | 0.59 | 0.58 | 0.57 | 0.6 | 0.59 | 0.6 | 0.58 | 0.57 | 0.53 |
| 0.6 | 0.59 | 0.59 | 0.59 | 0.6 | 0.57 | 0.58 | 0.58 | 0.57 | 0.6 | 0.59 | 0.6 | 0.59 | 0.57 | 0.54 |
| 0.6 | 0.59 | 0.58 | 0.58 | 0.59 | 0.57 | 0.59 | 0.58 | 0.58 | 0.6 | 0.59 | 0.6 | 0.56 | 0.59 | 0.54 |

| 10HzA302 | 30HzA302 | 60HzA302 | 10HzD303 | 30HzD303 | 60HzD303 | 10HzY304 | 30HzY304 | 60HzY304 | 10Hz F305 | 30Hz F305 | 60Hz F305 | 10HzK306 | 30HzK306 | 60HzK306 |
| --- | --- | --- | --- | --- | --- | --- | --- | --- | --- | --- | --- | --- | --- | --- |
| 0.59 | 0.58 | 0.56 | 0.6 | 0.58 | 0.59 | 0.6 | 0.58 | 0.58 | 0.58 | 0.59 | 0.58 | 0.57 | 0.56 | 0.56 |
| 0.56 | 0.58 | 0.61 | 0.6 | 0.58 | 0.58 | 0.61 | 0.57 | 0.58 | 0.59 | 0.58 | 0.57 | 0.57 | 0.58 | 0.56 |
| 0.55 | 0.57 | 0.6 | 0.6 | 0.57 | 0.58 | 0.58 | 0.6 | 0.57 | 0.59 | 0.58 | 0.57 | 0.56 | 0.56 | 0.56 |
| 0.57 | 0.56 | 0.57 | 0.6 | 0.58 | 0.58 | 0.59 | 0.58 | 0.57 | 0.59 | 0.58 | 0.57 | 0.57 | 0.57 | 0.56 |
| 0.63 | 0.58 | 0.59 | 0.59 | 0.58 | 0.58 | 0.59 | 0.58 | 0.58 | 0.61 | 0.58 | 0.56 | 0.57 | 0.57 | 0.56 |
| 0.57 | 0.56 | 0.6 | 0.6 | 0.58 | 0.57 | 0.59 | 0.58 | 0.57 | 0.59 | 0.57 | 0.55 | 0.58 | 0.57 | 0.57 |
| 0.57 | 0.59 | 0.56 | 0.58 | 0.58 | 0.58 | 0.61 | 0.58 | 0.58 | 0.6 | 0.56 | 0.55 | 0.56 | 0.57 | 0.57 |
| 0.56 | 0.57 | 0.57 | 0.6 | 0.58 | 0.57 | 0.59 | 0.59 | 0.57 | 0.6 | 0.57 | 0.55 | 0.56 | 0.57 | 0.55 |
| 0.58 | 0.54 | 0.58 | 0.6 | 0.58 | 0.57 | 0.61 | 0.58 | 0.58 | 0.61 | 0.58 | 0.54 | 0.57 | 0.58 | 0.56 |
| 0.63 | 0.58 | 0.58 | 0.61 | 0.58 | 0.57 | 0.6 | 0.59 | 0.57 | 0.59 | 0.57 | 0.53 | 0.56 | 0.58 | 0.56 |
| 0.59 | 0.57 | 0.59 | 0.6 | 0.58 | 0.57 | 0.59 | 0.58 | 0.57 | 0.61 | 0.57 | 0.52 | 0.56 | 0.57 | 0.55 |
| 0.58 | 0.56 | 0.59 | 0.59 | 0.57 | 0.56 | 0.58 | 0.58 | 0.57 | 0.6 | 0.57 | 0.49 | 0.58 | 0.56 | 0.56 |
| 0.62 | 0.56 | 0.59 | 0.61 | 0.58 | 0.55 | 0.59 | 0.59 | 0.57 | 0.59 | 0.55 | 0.47 | 0.58 | 0.56 | 0.55 |
| 0.61 | 0.59 | 0.6 | 0.59 | 0.57 | 0.55 | 0.6 | 0.59 | 0.56 | 0.59 | 0.54 | 0.44 | 0.56 | 0.56 | 0.55 |
| 0.59 | 0.58 | 0.58 | 0.59 | 0.57 | 0.53 | 0.58 | 0.59 | 0.56 | 0.59 | 0.54 | 0.42 | 0.56 | 0.57 | 0.55 |
| 0.56 | 0.56 | 0.58 | 0.59 | 0.57 | 0.52 | 0.61 | 0.57 | 0.56 | 0.6 | 0.52 | 0.36 | 0.57 | 0.56 | 0.54 |
| 0.58 | 0.57 | 0.59 | 0.59 | 0.57 | 0.51 | 0.6 | 0.59 | 0.56 | 0.58 | 0.48 | 0.3 | 0.57 | 0.57 | 0.54 |
| 0.63 | 0.58 | 0.57 | 0.59 | 0.56 | 0.49 | 0.6 | 0.57 | 0.56 | 0.57 | 0.45 | 0.21 | 0.57 | 0.56 | 0.54 |
| 0.61 | 0.56 | 0.59 | 0.6 | 0.55 | 0.46 | 0.6 | 0.58 | 0.55 | 0.57 | 0.37 | 0.13 | 0.56 | 0.56 | 0.53 |
| 0.57 | 0.6 | 0.57 | 0.59 | 0.54 | 0.43 | 0.59 | 0.59 | 0.54 | 0.55 | 0.22 | 0.05 | 0.56 | 0.55 | 0.52 |
| 0.54 | 0.6 | 0.57 | 0.6 | 0.53 | 0.4 | 0.6 | 0.58 | 0.53 | 0.42 | 0.05 | 0.01 | 0.57 | 0.57 | 0.51 |
| 0.58 | 0.54 | 0.57 | 0.59 | 0.51 | 0.35 | 0.58 | 0.57 | 0.53 | 0.02 | 0.01 | 0.01 | 0.55 | 0.56 | 0.5 |
| 0.61 | 0.55 | 0.56 | 0.6 | 0.48 | 0.28 | 0.61 | 0.57 | 0.52 | 0.35 | 0.03 | 0 | 0.56 | 0.56 | 0.49 |
| 0.57 | 0.58 | 0.55 | 0.58 | 0.43 | 0.19 | 0.59 | 0.56 | 0.5 | 0.51 | 0.2 | 0.04 | 0.57 | 0.56 | 0.48 |
| 0.56 | 0.57 | 0.54 | 0.57 | 0.34 | 0.1 | 0.59 | 0.57 | 0.49 | 0.56 | 0.35 | 0.11 | 0.57 | 0.54 | 0.46 |
| 0.57 | 0.56 | 0.58 | 0.51 | 0.19 | 0.03 | 0.59 | 0.56 | 0.46 | 0.58 | 0.44 | 0.19 | 0.57 | 0.53 | 0.45 |
| 0.57 | 0.58 | 0.55 | 0.34 | 0.03 | 0 | 0.61 | 0.54 | 0.44 | 0.59 | 0.48 | 0.27 | 0.57 | 0.53 | 0.42 |
| 0.59 | 0.59 | 0.54 | 0.03 | 0 | 0 | 0.58 | 0.55 | 0.41 | 0.58 | 0.5 | 0.34 | 0.57 | 0.51 | 0.38 |
| 0.53 | 0.55 | 0.53 | 0.4 | 0.05 | 0.01 | 0.59 | 0.52 | 0.36 | 0.59 | 0.52 | 0.39 | 0.57 | 0.5 | 0.33 |
| 0.59 | 0.55 | 0.52 | 0.53 | 0.22 | 0.03 | 0.57 | 0.5 | 0.3 | 0.59 | 0.53 | 0.41 | 0.55 | 0.46 | 0.27 |
| 0.57 | 0.57 | 0.52 | 0.56 | 0.35 | 0.11 | 0.58 | 0.46 | 0.22 | 0.6 | 0.53 | 0.44 | 0.54 | 0.43 | 0.18 |
| 0.58 | 0.55 | 0.49 | 0.58 | 0.42 | 0.19 | 0.56 | 0.39 | 0.13 | 0.59 | 0.53 | 0.45 | 0.53 | 0.35 | 0.09 |
| 0.59 | 0.57 | 0.48 | 0.57 | 0.45 | 0.27 | 0.55 | 0.26 | 0.05 | 0.58 | 0.54 | 0.47 | 0.5 | 0.18 | 0.02 |
| 0.57 | 0.54 | 0.47 | 0.58 | 0.49 | 0.34 | 0.44 | 0.09 | 0.01 | 0.59 | 0.56 | 0.5 | 0.32 | 0.03 | 0 |
| 0.55 | 0.51 | 0.44 | 0.59 | 0.53 | 0.39 | 0.05 | -0.01 | 0.01 | 0.6 | 0.56 | 0.52 | 0.01 | 0.01 | 0 |
| 0.59 | 0.53 | 0.4 | 0.59 | 0.54 | 0.43 | 0.23 | 0.02 | 0 | 0.6 | 0.57 | 0.53 | 0.4 | 0.05 | 0.01 |
| 0.58 | 0.51 | 0.38 | 0.59 | 0.55 | 0.47 | 0.49 | 0.14 | 0.02 | 0.6 | 0.57 | 0.55 | 0.51 | 0.23 | 0.05 |
| 0.61 | 0.47 | 0.29 | 0.6 | 0.56 | 0.49 | 0.53 | 0.3 | 0.07 | 0.6 | 0.58 | 0.55 | 0.54 | 0.35 | 0.12 |
| 0.54 | 0.41 | 0.21 | 0.6 | 0.55 | 0.51 | 0.56 | 0.4 | 0.16 | 0.59 | 0.56 | 0.56 | 0.55 | 0.42 | 0.21 |
| 0.57 | 0.36 | 0.11 | 0.59 | 0.57 | 0.52 | 0.58 | 0.46 | 0.25 | 0.59 | 0.57 | 0.57 | 0.55 | 0.47 | 0.29 |
| 0.56 | 0.29 | 0.07 | 0.6 | 0.57 | 0.53 | 0.6 | 0.52 | 0.32 | 0.6 | 0.58 | 0.57 | 0.56 | 0.49 | 0.36 |
| 0.43 | 0.08 | 0.02 | 0.59 | 0.58 | 0.54 | 0.59 | 0.52 | 0.38 | 0.6 | 0.58 | 0.56 | 0.56 | 0.52 | 0.4 |
| 0.01 | -0.01 | 0.01 | 0.59 | 0.58 | 0.55 | 0.58 | 0.54 | 0.42 | 0.6 | 0.58 | 0.58 | 0.56 | 0.53 | 0.43 |
| 0.23 | 0.02 | -0.01 | 0.59 | 0.58 | 0.55 | 0.59 | 0.54 | 0.44 | 0.6 | 0.56 | 0.59 | 0.55 | 0.54 | 0.45 |
| 0.47 | 0.16 | 0.01 | 0.59 | 0.58 | 0.56 | 0.58 | 0.55 | 0.47 | 0.61 | 0.59 | 0.58 | 0.56 | 0.54 | 0.47 |
| 0.53 | 0.28 | 0.07 | 0.61 | 0.58 | 0.56 | 0.59 | 0.58 | 0.5 | 0.58 | 0.58 | 0.59 | 0.56 | 0.56 | 0.49 |
| 0.58 | 0.39 | 0.16 | 0.6 | 0.58 | 0.57 | 0.58 | 0.58 | 0.5 | 0.6 | 0.59 | 0.58 | 0.56 | 0.56 | 0.5 |
| 0.58 | 0.44 | 0.24 | 0.59 | 0.58 | 0.57 | 0.6 | 0.58 | 0.52 | 0.6 | 0.58 | 0.59 | 0.56 | 0.57 | 0.51 |
| 0.56 | 0.5 | 0.32 | 0.6 | 0.59 | 0.58 | 0.57 | 0.55 | 0.52 | 0.59 | 0.58 | 0.59 | 0.56 | 0.57 | 0.52 |
| 0.59 | 0.52 | 0.37 | 0.59 | 0.58 | 0.58 | 0.61 | 0.57 | 0.53 | 0.6 | 0.59 | 0.59 | 0.56 | 0.57 | 0.52 |
| 0.61 | 0.51 | 0.41 | 0.6 | 0.59 | 0.58 | 0.59 | 0.58 | 0.53 | 0.6 | 0.58 | 0.59 | 0.55 | 0.56 | 0.53 |
| 0.58 | 0.56 | 0.46 | 0.6 | 0.59 | 0.58 | 0.59 | 0.59 | 0.55 | 0.59 | 0.58 | 0.59 | 0.57 | 0.57 | 0.54 |
| 0.57 | 0.52 | 0.47 | 0.59 | 0.59 | 0.58 | 0.6 | 0.59 | 0.55 | 0.6 | 0.59 | 0.59 | 0.57 | 0.56 | 0.54 |
| 0.6 | 0.56 | 0.48 | 0.59 | 0.58 | 0.59 | 0.59 | 0.59 | 0.55 | 0.61 | 0.59 | 0.58 | 0.55 | 0.57 | 0.54 |
| 0.57 | 0.58 | 0.51 | 0.59 | 0.58 | 0.58 | 0.59 | 0.58 | 0.56 | 0.6 | 0.58 | 0.59 | 0.54 | 0.57 | 0.55 |
| 0.59 | 0.55 | 0.53 | 0.59 | 0.59 | 0.59 | 0.58 | 0.57 | 0.56 | 0.6 | 0.6 | 0.6 | 0.57 | 0.57 | 0.55 |
| 0.58 | 0.56 | 0.54 | 0.6 | 0.58 | 0.59 | 0.58 | 0.59 | 0.57 | 0.59 | 0.58 | 0.59 | 0.56 | 0.56 | 0.55 |
| 0.57 | 0.57 | 0.54 | 0.6 | 0.59 | 0.59 | 0.58 | 0.59 | 0.57 | 0.61 | 0.58 | 0.59 | 0.56 | 0.57 | 0.56 |
| 0.61 | 0.56 | 0.55 | 0.6 | 0.58 | 0.59 | 0.59 | 0.58 | 0.58 | 0.6 | 0.58 | 0.59 | 0.58 | 0.57 | 0.56 |
| 0.57 | 0.58 | 0.55 | 0.6 | 0.59 | 0.59 | 0.59 | 0.59 | 0.56 | 0.6 | 0.58 | 0.6 | 0.56 | 0.57 | 0.55 |
| 0.58 | 0.6 | 0.58 | 0.6 | 0.58 | 0.59 | 0.57 | 0.58 | 0.57 | 0.59 | 0.6 | 0.59 | 0.57 | 0.57 | 0.56 |
| 0.57 | 0.59 | 0.58 | 0.59 | 0.58 | 0.59 | 0.58 | 0.6 | 0.57 | 0.61 | 0.58 | 0.59 | 0.56 | 0.56 | 0.56 |
| 0.61 | 0.56 | 0.56 | 0.6 | 0.59 | 0.59 | 0.59 | 0.58 | 0.58 | 0.62 | 0.59 | 0.59 | 0.57 | 0.57 | 0.56 |

| 10HzQ307 | 30HzQ307 | 60HzQ307 | 10Hz I308 | 30Hz I308 | 60Hz I308 | 10HzG309 | 30HzG309 | 60HzG309 | 10Hz I310 | 30Hz I310 | 60Hz I310 | 10HzT313 | 30HzT313 | 60HzT313 |
| --- | --- | --- | --- | --- | --- | --- | --- | --- | --- | --- | --- | --- | --- | --- |
| 0.57 | 0.56 | 0.56 | 0.59 | 0.56 | 0.59 | 0.6 | 0.58 | 0.51 | 0.63 | 0.6 | 0.61 | 0.58 | 0.59 | 0.6 |
| 0.58 | 0.57 | 0.56 | 0.6 | 0.57 | 0.58 | 0.61 | 0.56 | 0.49 | 0.64 | 0.6 | 0.62 | 0.6 | 0.59 | 0.6 |
| 0.58 | 0.57 | 0.56 | 0.6 | 0.56 | 0.57 | 0.6 | 0.56 | 0.48 | 0.62 | 0.61 | 0.62 | 0.6 | 0.59 | 0.6 |
| 0.58 | 0.57 | 0.57 | 0.6 | 0.58 | 0.58 | 0.62 | 0.56 | 0.44 | 0.64 | 0.61 | 0.62 | 0.6 | 0.59 | 0.6 |
| 0.56 | 0.56 | 0.56 | 0.59 | 0.58 | 0.59 | 0.61 | 0.54 | 0.4 | 0.62 | 0.62 | 0.62 | 0.59 | 0.58 | 0.6 |
| 0.57 | 0.57 | 0.56 | 0.57 | 0.56 | 0.56 | 0.61 | 0.52 | 0.34 | 0.62 | 0.61 | 0.62 | 0.59 | 0.59 | 0.59 |
| 0.56 | 0.57 | 0.56 | 0.59 | 0.57 | 0.57 | 0.6 | 0.49 | 0.27 | 0.62 | 0.6 | 0.61 | 0.59 | 0.58 | 0.6 |
| 0.57 | 0.56 | 0.55 | 0.58 | 0.56 | 0.56 | 0.58 | 0.43 | 0.18 | 0.64 | 0.61 | 0.6 | 0.59 | 0.58 | 0.59 |
| 0.57 | 0.56 | 0.55 | 0.58 | 0.57 | 0.57 | 0.57 | 0.33 | 0.08 | 0.65 | 0.6 | 0.61 | 0.57 | 0.58 | 0.58 |
| 0.57 | 0.56 | 0.55 | 0.59 | 0.58 | 0.56 | 0.51 | 0.16 | 0.02 | 0.64 | 0.61 | 0.62 | 0.59 | 0.59 | 0.59 |
| 0.56 | 0.56 | 0.54 | 0.58 | 0.55 | 0.56 | 0.27 | 0.01 | 0.01 | 0.64 | 0.62 | 0.61 | 0.6 | 0.58 | 0.58 |
| 0.57 | 0.56 | 0.54 | 0.6 | 0.56 | 0.56 | 0.04 | 0 | 0.01 | 0.64 | 0.61 | 0.61 | 0.58 | 0.59 | 0.58 |
| 0.57 | 0.56 | 0.54 | 0.59 | 0.57 | 0.56 | 0.47 | 0.09 | 0.01 | 0.64 | 0.62 | 0.61 | 0.58 | 0.58 | 0.58 |
| 0.57 | 0.56 | 0.52 | 0.58 | 0.58 | 0.55 | 0.55 | 0.27 | 0.06 | 0.61 | 0.61 | 0.61 | 0.59 | 0.58 | 0.58 |
| 0.56 | 0.56 | 0.52 | 0.58 | 0.55 | 0.53 | 0.58 | 0.4 | 0.14 | 0.6 | 0.59 | 0.58 | 0.59 | 0.58 | 0.57 |
| 0.57 | 0.55 | 0.52 | 0.57 | 0.55 | 0.52 | 0.59 | 0.45 | 0.23 | 0.63 | 0.61 | 0.58 | 0.6 | 0.58 | 0.56 |
| 0.57 | 0.55 | 0.51 | 0.58 | 0.54 | 0.51 | 0.59 | 0.48 | 0.3 | 0.62 | 0.61 | 0.57 | 0.6 | 0.58 | 0.57 |
| 0.56 | 0.55 | 0.5 | 0.57 | 0.55 | 0.5 | 0.59 | 0.49 | 0.35 | 0.62 | 0.61 | 0.57 | 0.61 | 0.58 | 0.56 |
| 0.56 | 0.55 | 0.48 | 0.61 | 0.55 | 0.49 | 0.6 | 0.52 | 0.41 | 0.62 | 0.59 | 0.57 | 0.59 | 0.57 | 0.55 |
| 0.56 | 0.55 | 0.46 | 0.61 | 0.54 | 0.48 | 0.61 | 0.54 | 0.44 | 0.61 | 0.59 | 0.56 | 0.58 | 0.57 | 0.55 |
| 0.57 | 0.53 | 0.43 | 0.57 | 0.54 | 0.46 | 0.6 | 0.56 | 0.48 | 0.62 | 0.58 | 0.56 | 0.6 | 0.58 | 0.54 |
| 0.57 | 0.52 | 0.4 | 0.56 | 0.52 | 0.43 | 0.6 | 0.57 | 0.49 | 0.62 | 0.58 | 0.55 | 0.59 | 0.57 | 0.52 |
| 0.56 | 0.51 | 0.36 | 0.57 | 0.5 | 0.39 | 0.61 | 0.58 | 0.52 | 0.64 | 0.58 | 0.52 | 0.6 | 0.56 | 0.5 |
| 0.56 | 0.47 | 0.3 | 0.57 | 0.48 | 0.32 | 0.61 | 0.57 | 0.53 | 0.61 | 0.57 | 0.48 | 0.58 | 0.55 | 0.47 |
| 0.56 | 0.44 | 0.23 | 0.57 | 0.44 | 0.25 | 0.61 | 0.57 | 0.54 | 0.62 | 0.57 | 0.45 | 0.59 | 0.54 | 0.46 |
| 0.54 | 0.38 | 0.14 | 0.55 | 0.39 | 0.17 | 0.61 | 0.58 | 0.55 | 0.6 | 0.57 | 0.4 | 0.58 | 0.52 | 0.42 |
| 0.53 | 0.27 | 0.06 | 0.56 | 0.33 | 0.07 | 0.6 | 0.58 | 0.55 | 0.63 | 0.51 | 0.33 | 0.58 | 0.5 | 0.38 |
| 0.44 | 0.09 | 0 | 0.51 | 0.15 | 0.02 | 0.6 | 0.58 | 0.56 | 0.6 | 0.48 | 0.25 | 0.58 | 0.48 | 0.32 |
| 0.06 | -0.01 | 0.01 | 0.26 | 0.02 | -0.01 | 0.61 | 0.58 | 0.57 | 0.59 | 0.38 | 0.14 | 0.58 | 0.46 | 0.26 |
| 0.21 | 0.01 | 0 | 0.02 | 0 | 0.01 | 0.6 | 0.59 | 0.57 | 0.57 | 0.26 | 0.06 | 0.56 | 0.42 | 0.18 |
| 0.47 | 0.14 | 0.02 | 0.4 | 0.07 | 0.01 | 0.62 | 0.59 | 0.58 | 0.44 | 0.07 | 0.02 | 0.56 | 0.33 | 0.09 |
| 0.53 | 0.29 | 0.08 | 0.52 | 0.22 | 0.05 | 0.61 | 0.59 | 0.58 | 0.05 | 0.02 | 0 | 0.52 | 0.18 | 0.02 |
| 0.54 | 0.39 | 0.16 | 0.56 | 0.34 | 0.12 | 0.61 | 0.58 | 0.59 | 0.38 | 0.03 | 0 | 0.32 | 0.03 | 0 |
| 0.54 | 0.44 | 0.24 | 0.57 | 0.43 | 0.21 | 0.61 | 0.59 | 0.59 | 0.52 | 0.21 | 0.03 | 0.01 | 0.01 | 0 |
| 0.55 | 0.45 | 0.3 | 0.57 | 0.46 | 0.3 | 0.62 | 0.58 | 0.59 | 0.58 | 0.36 | 0.13 | 0.38 | 0.06 | 0 |
| 0.56 | 0.48 | 0.35 | 0.57 | 0.51 | 0.36 | 0.61 | 0.59 | 0.59 | 0.57 | 0.44 | 0.2 | 0.53 | 0.23 | 0.03 |
| 0.56 | 0.51 | 0.4 | 0.56 | 0.52 | 0.39 | 0.61 | 0.59 | 0.59 | 0.6 | 0.49 | 0.31 | 0.55 | 0.37 | 0.11 |
| 0.57 | 0.53 | 0.44 | 0.59 | 0.53 | 0.44 | 0.62 | 0.6 | 0.58 | 0.62 | 0.54 | 0.38 | 0.58 | 0.45 | 0.21 |
| 0.57 | 0.54 | 0.46 | 0.57 | 0.53 | 0.47 | 0.61 | 0.59 | 0.6 | 0.62 | 0.56 | 0.42 | 0.58 | 0.49 | 0.28 |
| 0.57 | 0.54 | 0.48 | 0.59 | 0.54 | 0.48 | 0.6 | 0.59 | 0.58 | 0.63 | 0.57 | 0.46 | 0.58 | 0.52 | 0.35 |
| 0.56 | 0.55 | 0.49 | 0.59 | 0.55 | 0.5 | 0.6 | 0.6 | 0.59 | 0.62 | 0.56 | 0.49 | 0.59 | 0.53 | 0.4 |
| 0.57 | 0.55 | 0.5 | 0.57 | 0.55 | 0.52 | 0.61 | 0.59 | 0.6 | 0.61 | 0.59 | 0.52 | 0.59 | 0.55 | 0.44 |
| 0.57 | 0.55 | 0.52 | 0.6 | 0.55 | 0.52 | 0.61 | 0.59 | 0.59 | 0.61 | 0.6 | 0.54 | 0.58 | 0.55 | 0.47 |
| 0.56 | 0.55 | 0.52 | 0.6 | 0.56 | 0.55 | 0.61 | 0.59 | 0.6 | 0.59 | 0.59 | 0.55 | 0.58 | 0.56 | 0.5 |
| 0.57 | 0.56 | 0.53 | 0.58 | 0.58 | 0.54 | 0.61 | 0.58 | 0.6 | 0.61 | 0.59 | 0.55 | 0.59 | 0.56 | 0.52 |
| 0.57 | 0.56 | 0.54 | 0.6 | 0.55 | 0.55 | 0.61 | 0.58 | 0.6 | 0.64 | 0.6 | 0.58 | 0.6 | 0.57 | 0.52 |
| 0.57 | 0.56 | 0.54 | 0.58 | 0.56 | 0.56 | 0.61 | 0.6 | 0.6 | 0.63 | 0.6 | 0.57 | 0.59 | 0.58 | 0.53 |
| 0.57 | 0.56 | 0.54 | 0.56 | 0.55 | 0.57 | 0.62 | 0.59 | 0.6 | 0.65 | 0.61 | 0.59 | 0.58 | 0.57 | 0.54 |
| 0.58 | 0.55 | 0.55 | 0.55 | 0.54 | 0.56 | 0.62 | 0.59 | 0.6 | 0.63 | 0.61 | 0.59 | 0.59 | 0.58 | 0.55 |
| 0.57 | 0.56 | 0.55 | 0.58 | 0.55 | 0.56 | 0.6 | 0.59 | 0.6 | 0.64 | 0.6 | 0.6 | 0.59 | 0.58 | 0.56 |
| 0.58 | 0.56 | 0.55 | 0.57 | 0.57 | 0.58 | 0.6 | 0.59 | 0.6 | 0.61 | 0.6 | 0.6 | 0.6 | 0.57 | 0.56 |
| 0.58 | 0.56 | 0.55 | 0.6 | 0.55 | 0.57 | 0.61 | 0.59 | 0.6 | 0.63 | 0.62 | 0.61 | 0.59 | 0.58 | 0.57 |
| 0.57 | 0.56 | 0.56 | 0.58 | 0.56 | 0.57 | 0.6 | 0.59 | 0.6 | 0.62 | 0.61 | 0.61 | 0.6 | 0.59 | 0.58 |
| 0.57 | 0.56 | 0.56 | 0.58 | 0.57 | 0.57 | 0.61 | 0.6 | 0.58 | 0.64 | 0.63 | 0.6 | 0.59 | 0.58 | 0.57 |
| 0.56 | 0.57 | 0.57 | 0.58 | 0.56 | 0.59 | 0.61 | 0.59 | 0.59 | 0.64 | 0.61 | 0.6 | 0.6 | 0.58 | 0.57 |
| 0.56 | 0.57 | 0.57 | 0.57 | 0.56 | 0.57 | 0.61 | 0.59 | 0.6 | 0.65 | 0.61 | 0.62 | 0.58 | 0.58 | 0.58 |
| 0.58 | 0.56 | 0.56 | 0.57 | 0.58 | 0.58 | 0.62 | 0.59 | 0.6 | 0.63 | 0.59 | 0.61 | 0.6 | 0.58 | 0.58 |
| 0.57 | 0.56 | 0.56 | 0.57 | 0.57 | 0.57 | 0.6 | 0.59 | 0.6 | 0.62 | 0.59 | 0.62 | 0.59 | 0.59 | 0.59 |
| 0.57 | 0.56 | 0.56 | 0.58 | 0.56 | 0.59 | 0.6 | 0.59 | 0.61 | 0.6 | 0.61 | 0.62 | 0.59 | 0.59 | 0.59 |
| 0.58 | 0.57 | 0.57 | 0.59 | 0.57 | 0.58 | 0.62 | 0.59 | 0.61 | 0.62 | 0.6 | 0.62 | 0.59 | 0.58 | 0.59 |
| 0.57 | 0.57 | 0.56 | 0.58 | 0.58 | 0.59 | 0.62 | 0.6 | 0.61 | 0.63 | 0.6 | 0.62 | 0.58 | 0.58 | 0.59 |
| 0.57 | 0.57 | 0.56 | 0.57 | 0.56 | 0.59 | 0.6 | 0.6 | 0.6 | 0.6 | 0.61 | 0.61 | 0.58 | 0.58 | 0.59 |
| 0.57 | 0.56 | 0.57 | 0.58 | 0.58 | 0.59 | 0.61 | 0.59 | 0.61 | 0.62 | 0.61 | 0.62 | 0.61 | 0.59 | 0.59 |

| 10HzN314 | 30HzN314 | 60HzN314 | 10HzT317 | 30HzT317 | 60HzT317 | 10HzG318 | 30HzG318 | 60HzG318 | 10HzQ319 | 30HzQ319 | 60HzQ319 | 10HzM321 | 30HzM321 | 60HzM321 |
| --- | --- | --- | --- | --- | --- | --- | --- | --- | --- | --- | --- | --- | --- | --- |
| 0.61 | 0.61 | 0.6 | 0.6 | 0.54 | 0.42 | 0.59 | 0.57 | 0.56 | 0.6 | 0.59 | 0.6 | 0.65 | 0.61 | 0.63 |
| 0.6 | 0.6 | 0.58 | 0.61 | 0.53 | 0.36 | 0.6 | 0.57 | 0.56 | 0.6 | 0.58 | 0.59 | 0.62 | 0.61 | 0.61 |
| 0.6 | 0.6 | 0.58 | 0.6 | 0.5 | 0.3 | 0.58 | 0.58 | 0.57 | 0.6 | 0.59 | 0.59 | 0.62 | 0.63 | 0.61 |
| 0.6 | 0.61 | 0.6 | 0.6 | 0.46 | 0.22 | 0.6 | 0.57 | 0.56 | 0.6 | 0.59 | 0.59 | 0.62 | 0.62 | 0.63 |
| 0.6 | 0.61 | 0.6 | 0.59 | 0.37 | 0.13 | 0.59 | 0.58 | 0.56 | 0.6 | 0.58 | 0.6 | 0.65 | 0.62 | 0.64 |
| 0.59 | 0.6 | 0.59 | 0.55 | 0.2 | 0.03 | 0.6 | 0.56 | 0.55 | 0.59 | 0.59 | 0.59 | 0.64 | 0.65 | 0.61 |
| 0.6 | 0.61 | 0.6 | 0.36 | 0.04 | 0 | 0.6 | 0.57 | 0.55 | 0.61 | 0.59 | 0.59 | 0.65 | 0.63 | 0.64 |
| 0.61 | 0.61 | 0.6 | 0.02 | 0 | 0 | 0.61 | 0.58 | 0.53 | 0.59 | 0.58 | 0.59 | 0.63 | 0.66 | 0.62 |
| 0.59 | 0.59 | 0.6 | 0.43 | 0.06 | 0.01 | 0.6 | 0.56 | 0.51 | 0.6 | 0.58 | 0.59 | 0.63 | 0.59 | 0.62 |
| 0.61 | 0.59 | 0.6 | 0.55 | 0.25 | 0.05 | 0.59 | 0.56 | 0.49 | 0.6 | 0.58 | 0.59 | 0.63 | 0.6 | 0.63 |
| 0.61 | 0.61 | 0.6 | 0.59 | 0.4 | 0.14 | 0.6 | 0.54 | 0.48 | 0.6 | 0.59 | 0.59 | 0.64 | 0.63 | 0.63 |
| 0.62 | 0.61 | 0.59 | 0.6 | 0.47 | 0.23 | 0.58 | 0.54 | 0.45 | 0.61 | 0.59 | 0.58 | 0.65 | 0.64 | 0.62 |
| 0.59 | 0.61 | 0.6 | 0.6 | 0.51 | 0.3 | 0.6 | 0.53 | 0.42 | 0.6 | 0.58 | 0.58 | 0.63 | 0.63 | 0.63 |
| 0.62 | 0.6 | 0.59 | 0.59 | 0.53 | 0.37 | 0.59 | 0.51 | 0.37 | 0.6 | 0.59 | 0.58 | 0.67 | 0.65 | 0.62 |
| 0.61 | 0.61 | 0.6 | 0.6 | 0.55 | 0.4 | 0.59 | 0.48 | 0.3 | 0.6 | 0.58 | 0.58 | 0.64 | 0.62 | 0.61 |
| 0.6 | 0.62 | 0.59 | 0.61 | 0.55 | 0.44 | 0.59 | 0.44 | 0.21 | 0.6 | 0.58 | 0.58 | 0.65 | 0.63 | 0.61 |
| 0.61 | 0.6 | 0.59 | 0.59 | 0.57 | 0.47 | 0.58 | 0.37 | 0.12 | 0.61 | 0.58 | 0.58 | 0.66 | 0.6 | 0.62 |
| 0.6 | 0.6 | 0.59 | 0.61 | 0.57 | 0.49 | 0.55 | 0.24 | 0.04 | 0.6 | 0.57 | 0.57 | 0.62 | 0.59 | 0.62 |
| 0.61 | 0.61 | 0.59 | 0.6 | 0.58 | 0.49 | 0.42 | 0.04 | 0 | 0.6 | 0.59 | 0.56 | 0.65 | 0.62 | 0.62 |
| 0.6 | 0.59 | 0.58 | 0.61 | 0.57 | 0.5 | 0.03 | 0.01 | 0 | 0.6 | 0.58 | 0.56 | 0.61 | 0.63 | 0.6 |
| 0.61 | 0.6 | 0.58 | 0.62 | 0.56 | 0.5 | 0.37 | 0.04 | 0.01 | 0.6 | 0.57 | 0.55 | 0.67 | 0.64 | 0.61 |
| 0.6 | 0.6 | 0.59 | 0.59 | 0.56 | 0.52 | 0.54 | 0.2 | 0.04 | 0.6 | 0.58 | 0.54 | 0.66 | 0.64 | 0.61 |
| 0.62 | 0.61 | 0.59 | 0.58 | 0.56 | 0.52 | 0.57 | 0.36 | 0.11 | 0.6 | 0.58 | 0.53 | 0.66 | 0.63 | 0.6 |
| 0.61 | 0.6 | 0.58 | 0.6 | 0.56 | 0.53 | 0.57 | 0.45 | 0.2 | 0.6 | 0.57 | 0.52 | 0.66 | 0.63 | 0.61 |
| 0.6 | 0.61 | 0.57 | 0.61 | 0.58 | 0.54 | 0.59 | 0.49 | 0.29 | 0.6 | 0.56 | 0.5 | 0.67 | 0.6 | 0.6 |
| 0.59 | 0.6 | 0.58 | 0.59 | 0.6 | 0.56 | 0.6 | 0.5 | 0.36 | 0.6 | 0.56 | 0.49 | 0.64 | 0.61 | 0.6 |
| 0.6 | 0.61 | 0.57 | 0.59 | 0.59 | 0.56 | 0.58 | 0.54 | 0.4 | 0.59 | 0.55 | 0.46 | 0.66 | 0.64 | 0.59 |
| 0.62 | 0.59 | 0.57 | 0.6 | 0.6 | 0.57 | 0.59 | 0.54 | 0.44 | 0.59 | 0.54 | 0.41 | 0.67 | 0.65 | 0.6 |
| 0.61 | 0.6 | 0.56 | 0.61 | 0.6 | 0.57 | 0.61 | 0.55 | 0.47 | 0.59 | 0.52 | 0.37 | 0.63 | 0.61 | 0.57 |
| 0.62 | 0.59 | 0.55 | 0.59 | 0.59 | 0.57 | 0.59 | 0.56 | 0.48 | 0.59 | 0.48 | 0.3 | 0.65 | 0.62 | 0.57 |
| 0.62 | 0.59 | 0.56 | 0.61 | 0.59 | 0.58 | 0.6 | 0.56 | 0.51 | 0.58 | 0.44 | 0.22 | 0.64 | 0.58 | 0.6 |
| 0.6 | 0.6 | 0.54 | 0.6 | 0.6 | 0.59 | 0.6 | 0.55 | 0.51 | 0.58 | 0.36 | 0.12 | 0.66 | 0.62 | 0.55 |
| 0.6 | 0.59 | 0.52 | 0.62 | 0.6 | 0.59 | 0.6 | 0.57 | 0.53 | 0.52 | 0.2 | 0.04 | 0.65 | 0.63 | 0.55 |
| 0.6 | 0.59 | 0.53 | 0.61 | 0.61 | 0.6 | 0.59 | 0.57 | 0.55 | 0.3 | 0.02 | 0 | 0.66 | 0.61 | 0.54 |
| 0.59 | 0.58 | 0.5 | 0.6 | 0.62 | 0.59 | 0.6 | 0.57 | 0.56 | 0.04 | 0 | 0 | 0.65 | 0.6 | 0.53 |
| 0.62 | 0.58 | 0.47 | 0.6 | 0.61 | 0.59 | 0.59 | 0.57 | 0.56 | 0.46 | 0.1 | 0.02 | 0.63 | 0.59 | 0.49 |
| 0.61 | 0.57 | 0.46 | 0.62 | 0.6 | 0.59 | 0.61 | 0.58 | 0.56 | 0.55 | 0.26 | 0.07 | 0.66 | 0.6 | 0.48 |
| 0.59 | 0.56 | 0.42 | 0.6 | 0.6 | 0.59 | 0.59 | 0.58 | 0.57 | 0.57 | 0.38 | 0.16 | 0.62 | 0.58 | 0.42 |
| 0.59 | 0.52 | 0.36 | 0.61 | 0.6 | 0.6 | 0.6 | 0.59 | 0.56 | 0.58 | 0.46 | 0.26 | 0.64 | 0.54 | 0.4 |
| 0.58 | 0.49 | 0.3 | 0.6 | 0.6 | 0.6 | 0.59 | 0.58 | 0.57 | 0.59 | 0.5 | 0.34 | 0.62 | 0.51 | 0.33 |
| 0.58 | 0.44 | 0.23 | 0.6 | 0.6 | 0.59 | 0.6 | 0.58 | 0.57 | 0.59 | 0.53 | 0.4 | 0.61 | 0.47 | 0.24 |
| 0.58 | 0.39 | 0.15 | 0.63 | 0.61 | 0.6 | 0.6 | 0.57 | 0.58 | 0.59 | 0.54 | 0.44 | 0.57 | 0.41 | 0.15 |
| 0.54 | 0.28 | 0.06 | 0.61 | 0.6 | 0.6 | 0.61 | 0.58 | 0.58 | 0.6 | 0.54 | 0.47 | 0.57 | 0.27 | 0.05 |
| 0.49 | 0.11 | 0.01 | 0.62 | 0.61 | 0.61 | 0.6 | 0.57 | 0.59 | 0.59 | 0.56 | 0.5 | 0.53 | 0.1 | 0.01 |
| 0.09 | 0 | 0 | 0.6 | 0.61 | 0.6 | 0.6 | 0.57 | 0.58 | 0.6 | 0.57 | 0.51 | 0.12 | 0.01 | -0.01 |
| 0.19 | 0.01 | 0 | 0.6 | 0.6 | 0.6 | 0.62 | 0.57 | 0.59 | 0.59 | 0.57 | 0.52 | 0.21 | 0.01 | -0.01 |
| 0.5 | 0.14 | 0.01 | 0.62 | 0.6 | 0.6 | 0.59 | 0.58 | 0.58 | 0.59 | 0.57 | 0.54 | 0.55 | 0.14 | 0.02 |
| 0.56 | 0.31 | 0.07 | 0.6 | 0.61 | 0.59 | 0.6 | 0.59 | 0.58 | 0.6 | 0.57 | 0.55 | 0.59 | 0.34 | 0.07 |
| 0.57 | 0.42 | 0.17 | 0.61 | 0.6 | 0.6 | 0.61 | 0.58 | 0.59 | 0.6 | 0.58 | 0.55 | 0.6 | 0.43 | 0.19 |
| 0.59 | 0.48 | 0.25 | 0.63 | 0.6 | 0.58 | 0.59 | 0.57 | 0.59 | 0.6 | 0.58 | 0.56 | 0.65 | 0.48 | 0.27 |
| 0.59 | 0.52 | 0.33 | 0.62 | 0.6 | 0.58 | 0.58 | 0.58 | 0.58 | 0.6 | 0.58 | 0.56 | 0.62 | 0.56 | 0.34 |
| 0.61 | 0.55 | 0.38 | 0.6 | 0.6 | 0.6 | 0.6 | 0.58 | 0.59 | 0.59 | 0.58 | 0.57 | 0.62 | 0.57 | 0.4 |
| 0.59 | 0.56 | 0.43 | 0.62 | 0.6 | 0.6 | 0.61 | 0.58 | 0.59 | 0.6 | 0.58 | 0.57 | 0.61 | 0.59 | 0.43 |
| 0.59 | 0.57 | 0.46 | 0.61 | 0.6 | 0.6 | 0.58 | 0.59 | 0.58 | 0.6 | 0.57 | 0.58 | 0.62 | 0.61 | 0.48 |
| 0.6 | 0.59 | 0.47 | 0.62 | 0.61 | 0.6 | 0.6 | 0.57 | 0.59 | 0.6 | 0.59 | 0.58 | 0.62 | 0.61 | 0.49 |
| 0.6 | 0.57 | 0.52 | 0.61 | 0.6 | 0.61 | 0.6 | 0.58 | 0.6 | 0.6 | 0.58 | 0.58 | 0.64 | 0.6 | 0.53 |
| 0.6 | 0.59 | 0.52 | 0.61 | 0.61 | 0.6 | 0.59 | 0.58 | 0.59 | 0.59 | 0.59 | 0.59 | 0.61 | 0.62 | 0.54 |
| 0.6 | 0.59 | 0.53 | 0.6 | 0.6 | 0.6 | 0.6 | 0.58 | 0.59 | 0.61 | 0.58 | 0.59 | 0.63 | 0.62 | 0.56 |
| 0.62 | 0.59 | 0.55 | 0.6 | 0.61 | 0.6 | 0.59 | 0.58 | 0.6 | 0.61 | 0.58 | 0.58 | 0.64 | 0.59 | 0.57 |
| 0.6 | 0.61 | 0.54 | 0.61 | 0.61 | 0.6 | 0.59 | 0.57 | 0.59 | 0.6 | 0.58 | 0.59 | 0.64 | 0.63 | 0.58 |
| 0.6 | 0.61 | 0.56 | 0.62 | 0.61 | 0.6 | 0.6 | 0.58 | 0.58 | 0.61 | 0.59 | 0.59 | 0.66 | 0.61 | 0.57 |
| 0.58 | 0.59 | 0.57 | 0.61 | 0.61 | 0.61 | 0.58 | 0.58 | 0.57 | 0.62 | 0.59 | 0.6 | 0.64 | 0.61 | 0.57 |
| 0.59 | 0.59 | 0.57 | 0.61 | 0.61 | 0.61 | 0.6 | 0.57 | 0.57 | 0.59 | 0.59 | 0.59 | 0.62 | 0.6 | 0.57 |

| 10Hz I322 | 30Hz I322 | 60Hz I322 | 10HzN323 | 30HzN323 | 60HzN323 | 10HzL324 | 30HzL324 | 60HzL324 | 10HzY325 | 30HzY325 | 60HzY325 | 10HzT326 | 30HzT326 | 60HzT326 |
| --- | --- | --- | --- | --- | --- | --- | --- | --- | --- | --- | --- | --- | --- | --- |
| 0.58 | 0.6 | 0.61 | 0.59 | 0.59 | 0.58 | 0.58 | 0.59 | 0.59 | 0.55 | 0.58 | 0.61 | 0.57 | 0.58 | 0.55 |
| 0.61 | 0.6 | 0.61 | 0.59 | 0.59 | 0.58 | 0.59 | 0.59 | 0.6 | 0.56 | 0.58 | 0.6 | 0.54 | 0.56 | 0.55 |
| 0.59 | 0.59 | 0.6 | 0.59 | 0.57 | 0.57 | 0.6 | 0.58 | 0.59 | 0.56 | 0.58 | 0.6 | 0.54 | 0.55 | 0.53 |
| 0.61 | 0.59 | 0.59 | 0.59 | 0.57 | 0.57 | 0.59 | 0.6 | 0.59 | 0.55 | 0.58 | 0.61 | 0.53 | 0.6 | 0.56 |
| 0.6 | 0.6 | 0.59 | 0.59 | 0.58 | 0.58 | 0.59 | 0.6 | 0.6 | 0.54 | 0.58 | 0.6 | 0.53 | 0.56 | 0.55 |
| 0.61 | 0.6 | 0.6 | 0.58 | 0.59 | 0.59 | 0.58 | 0.58 | 0.61 | 0.55 | 0.57 | 0.6 | 0.58 | 0.59 | 0.56 |
| 0.63 | 0.61 | 0.58 | 0.59 | 0.58 | 0.59 | 0.61 | 0.59 | 0.6 | 0.55 | 0.58 | 0.6 | 0.6 | 0.57 | 0.54 |
| 0.59 | 0.6 | 0.6 | 0.59 | 0.58 | 0.59 | 0.59 | 0.58 | 0.61 | 0.54 | 0.57 | 0.58 | 0.53 | 0.57 | 0.56 |
| 0.6 | 0.6 | 0.61 | 0.57 | 0.58 | 0.59 | 0.6 | 0.56 | 0.59 | 0.52 | 0.59 | 0.57 | 0.54 | 0.56 | 0.56 |
| 0.58 | 0.59 | 0.6 | 0.6 | 0.56 | 0.58 | 0.57 | 0.57 | 0.59 | 0.55 | 0.58 | 0.6 | 0.55 | 0.57 | 0.53 |
| 0.58 | 0.59 | 0.6 | 0.6 | 0.57 | 0.58 | 0.58 | 0.58 | 0.59 | 0.57 | 0.6 | 0.6 | 0.53 | 0.54 | 0.55 |
| 0.6 | 0.61 | 0.6 | 0.59 | 0.59 | 0.58 | 0.57 | 0.59 | 0.59 | 0.54 | 0.58 | 0.59 | 0.53 | 0.56 | 0.53 |
| 0.59 | 0.61 | 0.59 | 0.59 | 0.59 | 0.58 | 0.59 | 0.58 | 0.58 | 0.54 | 0.59 | 0.6 | 0.54 | 0.58 | 0.53 |
| 0.6 | 0.62 | 0.6 | 0.58 | 0.58 | 0.59 | 0.59 | 0.59 | 0.6 | 0.54 | 0.56 | 0.58 | 0.56 | 0.54 | 0.53 |
| 0.59 | 0.61 | 0.59 | 0.58 | 0.57 | 0.58 | 0.57 | 0.58 | 0.6 | 0.59 | 0.59 | 0.59 | 0.54 | 0.57 | 0.52 |
| 0.6 | 0.59 | 0.59 | 0.58 | 0.57 | 0.58 | 0.6 | 0.59 | 0.59 | 0.52 | 0.57 | 0.58 | 0.56 | 0.56 | 0.54 |
| 0.6 | 0.58 | 0.59 | 0.6 | 0.57 | 0.57 | 0.59 | 0.59 | 0.6 | 0.55 | 0.57 | 0.58 | 0.57 | 0.57 | 0.51 |
| 0.58 | 0.6 | 0.59 | 0.59 | 0.58 | 0.58 | 0.59 | 0.59 | 0.59 | 0.57 | 0.57 | 0.57 | 0.56 | 0.53 | 0.51 |
| 0.6 | 0.61 | 0.6 | 0.59 | 0.57 | 0.58 | 0.59 | 0.58 | 0.58 | 0.54 | 0.57 | 0.58 | 0.59 | 0.55 | 0.48 |
| 0.59 | 0.61 | 0.59 | 0.59 | 0.59 | 0.57 | 0.6 | 0.58 | 0.58 | 0.59 | 0.56 | 0.58 | 0.56 | 0.53 | 0.48 |
| 0.59 | 0.61 | 0.58 | 0.58 | 0.58 | 0.57 | 0.58 | 0.6 | 0.58 | 0.56 | 0.55 | 0.57 | 0.54 | 0.54 | 0.45 |
| 0.59 | 0.57 | 0.59 | 0.6 | 0.57 | 0.58 | 0.58 | 0.58 | 0.57 | 0.57 | 0.56 | 0.57 | 0.58 | 0.54 | 0.42 |
| 0.57 | 0.6 | 0.6 | 0.59 | 0.57 | 0.57 | 0.59 | 0.58 | 0.57 | 0.53 | 0.56 | 0.55 | 0.52 | 0.52 | 0.39 |
| 0.59 | 0.59 | 0.61 | 0.6 | 0.58 | 0.58 | 0.6 | 0.59 | 0.58 | 0.56 | 0.57 | 0.54 | 0.51 | 0.49 | 0.35 |
| 0.58 | 0.59 | 0.6 | 0.57 | 0.57 | 0.57 | 0.59 | 0.58 | 0.57 | 0.55 | 0.58 | 0.53 | 0.53 | 0.46 | 0.3 |
| 0.57 | 0.62 | 0.59 | 0.58 | 0.58 | 0.57 | 0.57 | 0.56 | 0.56 | 0.55 | 0.58 | 0.54 | 0.53 | 0.41 | 0.2 |
| 0.6 | 0.59 | 0.58 | 0.58 | 0.58 | 0.56 | 0.58 | 0.57 | 0.56 | 0.55 | 0.56 | 0.54 | 0.51 | 0.33 | 0.12 |
| 0.6 | 0.6 | 0.58 | 0.58 | 0.57 | 0.56 | 0.58 | 0.58 | 0.55 | 0.54 | 0.57 | 0.54 | 0.46 | 0.22 | 0.06 |
| 0.58 | 0.6 | 0.57 | 0.58 | 0.58 | 0.56 | 0.59 | 0.57 | 0.55 | 0.58 | 0.53 | 0.5 | 0.31 | 0.06 | 0.02 |
| 0.59 | 0.6 | 0.57 | 0.59 | 0.57 | 0.55 | 0.59 | 0.57 | 0.52 | 0.55 | 0.54 | 0.49 | 0.05 | 0.02 | 0.01 |
| 0.61 | 0.58 | 0.57 | 0.57 | 0.57 | 0.55 | 0.6 | 0.57 | 0.51 | 0.58 | 0.55 | 0.49 | 0.29 | -0.01 | 0.02 |
| 0.59 | 0.6 | 0.57 | 0.59 | 0.55 | 0.54 | 0.59 | 0.56 | 0.5 | 0.53 | 0.55 | 0.46 | 0.47 | 0.15 | 0.04 |
| 0.6 | 0.58 | 0.55 | 0.58 | 0.57 | 0.52 | 0.59 | 0.55 | 0.48 | 0.52 | 0.53 | 0.41 | 0.56 | 0.33 | 0.1 |
| 0.56 | 0.59 | 0.56 | 0.59 | 0.56 | 0.51 | 0.58 | 0.53 | 0.44 | 0.53 | 0.52 | 0.38 | 0.54 | 0.41 | 0.19 |
| 0.59 | 0.58 | 0.55 | 0.59 | 0.56 | 0.5 | 0.57 | 0.53 | 0.41 | 0.54 | 0.46 | 0.31 | 0.55 | 0.46 | 0.25 |
| 0.62 | 0.59 | 0.52 | 0.57 | 0.55 | 0.49 | 0.58 | 0.53 | 0.35 | 0.53 | 0.43 | 0.22 | 0.52 | 0.48 | 0.33 |
| 0.59 | 0.59 | 0.51 | 0.58 | 0.55 | 0.47 | 0.57 | 0.48 | 0.29 | 0.52 | 0.37 | 0.15 | 0.56 | 0.51 | 0.37 |
| 0.6 | 0.56 | 0.5 | 0.59 | 0.54 | 0.45 | 0.57 | 0.44 | 0.22 | 0.5 | 0.25 | 0.06 | 0.54 | 0.53 | 0.42 |
| 0.62 | 0.53 | 0.48 | 0.59 | 0.52 | 0.42 | 0.54 | 0.38 | 0.12 | 0.44 | 0.09 | 0.01 | 0.57 | 0.54 | 0.46 |
| 0.57 | 0.53 | 0.45 | 0.57 | 0.52 | 0.37 | 0.53 | 0.21 | 0.02 | 0.07 | 0.01 | 0 | 0.54 | 0.57 | 0.46 |
| 0.58 | 0.53 | 0.43 | 0.57 | 0.5 | 0.31 | 0.34 | 0.04 | 0 | 0.2 | 0.01 | 0.01 | 0.54 | 0.54 | 0.48 |
| 0.6 | 0.55 | 0.4 | 0.56 | 0.44 | 0.23 | 0.01 | 0.01 | 0.01 | 0.45 | 0.13 | 0.01 | 0.56 | 0.58 | 0.47 |
| 0.6 | 0.53 | 0.37 | 0.55 | 0.37 | 0.14 | 0.42 | 0.06 | 0.01 | 0.52 | 0.3 | 0.08 | 0.54 | 0.55 | 0.51 |
| 0.6 | 0.53 | 0.31 | 0.53 | 0.26 | 0.06 | 0.55 | 0.25 | 0.06 | 0.52 | 0.4 | 0.18 | 0.5 | 0.55 | 0.51 |
| 0.6 | 0.47 | 0.26 | 0.44 | 0.09 | 0.01 | 0.55 | 0.39 | 0.13 | 0.52 | 0.47 | 0.26 | 0.57 | 0.56 | 0.52 |
| 0.57 | 0.43 | 0.16 | 0.05 | 0.01 | 0.01 | 0.57 | 0.47 | 0.24 | 0.53 | 0.5 | 0.33 | 0.54 | 0.55 | 0.53 |
| 0.54 | 0.32 | 0.07 | 0.26 | 0.02 | 0 | 0.57 | 0.49 | 0.31 | 0.54 | 0.54 | 0.38 | 0.55 | 0.53 | 0.54 |
| 0.51 | 0.16 | 0.01 | 0.5 | 0.17 | 0.02 | 0.59 | 0.51 | 0.39 | 0.55 | 0.53 | 0.41 | 0.53 | 0.53 | 0.54 |
| 0.21 | 0.01 | 0 | 0.55 | 0.33 | 0.09 | 0.59 | 0.54 | 0.42 | 0.52 | 0.54 | 0.49 | 0.55 | 0.55 | 0.56 |
| 0.09 | 0 | 0 | 0.56 | 0.43 | 0.19 | 0.59 | 0.54 | 0.46 | 0.54 | 0.54 | 0.49 | 0.54 | 0.55 | 0.56 |
| 0.45 | 0.11 | 0.01 | 0.56 | 0.47 | 0.27 | 0.57 | 0.56 | 0.47 | 0.56 | 0.55 | 0.5 | 0.56 | 0.56 | 0.53 |
| 0.55 | 0.3 | 0.05 | 0.57 | 0.51 | 0.34 | 0.57 | 0.57 | 0.52 | 0.56 | 0.56 | 0.51 | 0.52 | 0.55 | 0.52 |
| 0.59 | 0.4 | 0.15 | 0.58 | 0.53 | 0.39 | 0.59 | 0.57 | 0.51 | 0.55 | 0.58 | 0.55 | 0.56 | 0.56 | 0.54 |
| 0.6 | 0.47 | 0.26 | 0.59 | 0.54 | 0.43 | 0.59 | 0.55 | 0.53 | 0.54 | 0.58 | 0.52 | 0.58 | 0.56 | 0.56 |
| 0.57 | 0.53 | 0.33 | 0.58 | 0.54 | 0.47 | 0.6 | 0.55 | 0.54 | 0.57 | 0.58 | 0.53 | 0.55 | 0.56 | 0.56 |
| 0.59 | 0.53 | 0.38 | 0.59 | 0.55 | 0.48 | 0.6 | 0.57 | 0.55 | 0.56 | 0.57 | 0.55 | 0.58 | 0.58 | 0.55 |
| 0.62 | 0.58 | 0.43 | 0.6 | 0.56 | 0.5 | 0.58 | 0.57 | 0.55 | 0.53 | 0.58 | 0.56 | 0.58 | 0.58 | 0.56 |
| 0.6 | 0.55 | 0.46 | 0.58 | 0.55 | 0.52 | 0.57 | 0.57 | 0.56 | 0.56 | 0.56 | 0.56 | 0.56 | 0.57 | 0.57 |
| 0.58 | 0.57 | 0.49 | 0.59 | 0.55 | 0.52 | 0.57 | 0.58 | 0.57 | 0.56 | 0.57 | 0.57 | 0.57 | 0.56 | 0.55 |
| 0.57 | 0.57 | 0.51 | 0.59 | 0.57 | 0.53 | 0.57 | 0.57 | 0.57 | 0.55 | 0.57 | 0.57 | 0.54 | 0.59 | 0.53 |
| 0.56 | 0.58 | 0.54 | 0.58 | 0.57 | 0.54 | 0.59 | 0.57 | 0.57 | 0.54 | 0.58 | 0.59 | 0.55 | 0.57 | 0.56 |
| 0.61 | 0.59 | 0.53 | 0.61 | 0.57 | 0.55 | 0.58 | 0.57 | 0.59 | 0.54 | 0.58 | 0.58 | 0.56 | 0.56 | 0.55 |
| 0.61 | 0.57 | 0.54 | 0.59 | 0.56 | 0.55 | 0.57 | 0.58 | 0.58 | 0.55 | 0.57 | 0.57 | 0.58 | 0.55 | 0.57 |

| 10HzD327 | 30HzD327 | 60HzD327 | 10HzE329 | 30HzE329 | 60HzE329 | 10HzT330 | 30HzT330 | 60HzT330 | 10HzG331 | 30HzG331 | 60HzG331 | 10HzK332 | 30HzK332 | 60HzK332 |
| --- | --- | --- | --- | --- | --- | --- | --- | --- | --- | --- | --- | --- | --- | --- |
| 0.6 | 0.58 | 0.59 | 0.6 | 0.6 | 0.6 | 0.61 | 0.58 | 0.48 | 0.59 | 0.55 | 0.55 | 0.58 | 0.56 | 0.55 |
| 0.6 | 0.58 | 0.58 | 0.59 | 0.61 | 0.59 | 0.61 | 0.57 | 0.44 | 0.59 | 0.56 | 0.55 | 0.57 | 0.55 | 0.55 |
| 0.6 | 0.58 | 0.59 | 0.59 | 0.59 | 0.59 | 0.61 | 0.55 | 0.39 | 0.58 | 0.57 | 0.54 | 0.57 | 0.55 | 0.56 |
| 0.6 | 0.59 | 0.6 | 0.6 | 0.6 | 0.59 | 0.61 | 0.53 | 0.36 | 0.59 | 0.56 | 0.54 | 0.58 | 0.56 | 0.56 |
| 0.61 | 0.58 | 0.6 | 0.6 | 0.6 | 0.59 | 0.59 | 0.51 | 0.29 | 0.58 | 0.54 | 0.53 | 0.57 | 0.56 | 0.56 |
| 0.6 | 0.58 | 0.6 | 0.6 | 0.59 | 0.59 | 0.6 | 0.47 | 0.22 | 0.57 | 0.55 | 0.53 | 0.57 | 0.56 | 0.55 |
| 0.59 | 0.58 | 0.6 | 0.59 | 0.59 | 0.59 | 0.58 | 0.4 | 0.13 | 0.59 | 0.54 | 0.52 | 0.57 | 0.56 | 0.56 |
| 0.6 | 0.58 | 0.6 | 0.6 | 0.6 | 0.59 | 0.57 | 0.28 | 0.05 | 0.59 | 0.54 | 0.5 | 0.57 | 0.55 | 0.55 |
| 0.61 | 0.59 | 0.6 | 0.59 | 0.6 | 0.59 | 0.5 | 0.12 | 0.01 | 0.59 | 0.54 | 0.49 | 0.56 | 0.56 | 0.55 |
| 0.61 | 0.58 | 0.6 | 0.6 | 0.6 | 0.58 | 0.23 | 0 | 0 | 0.58 | 0.54 | 0.48 | 0.57 | 0.56 | 0.56 |
| 0.61 | 0.58 | 0.61 | 0.59 | 0.59 | 0.59 | 0.02 | 0.01 | 0 | 0.59 | 0.52 | 0.45 | 0.58 | 0.55 | 0.55 |
| 0.6 | 0.57 | 0.59 | 0.6 | 0.59 | 0.59 | 0.44 | 0.04 | 0.01 | 0.58 | 0.51 | 0.42 | 0.57 | 0.56 | 0.55 |
| 0.6 | 0.59 | 0.6 | 0.6 | 0.59 | 0.58 | 0.54 | 0.21 | 0.03 | 0.58 | 0.51 | 0.39 | 0.58 | 0.55 | 0.56 |
| 0.61 | 0.57 | 0.6 | 0.59 | 0.59 | 0.57 | 0.58 | 0.36 | 0.09 | 0.6 | 0.49 | 0.32 | 0.58 | 0.55 | 0.55 |
| 0.6 | 0.59 | 0.6 | 0.59 | 0.6 | 0.57 | 0.59 | 0.43 | 0.19 | 0.58 | 0.46 | 0.26 | 0.58 | 0.55 | 0.55 |
| 0.6 | 0.57 | 0.59 | 0.6 | 0.59 | 0.56 | 0.62 | 0.48 | 0.27 | 0.58 | 0.4 | 0.18 | 0.57 | 0.56 | 0.55 |
| 0.61 | 0.58 | 0.59 | 0.59 | 0.59 | 0.56 | 0.62 | 0.51 | 0.32 | 0.56 | 0.3 | 0.08 | 0.57 | 0.56 | 0.54 |
| 0.59 | 0.58 | 0.59 | 0.6 | 0.59 | 0.55 | 0.6 | 0.53 | 0.37 | 0.49 | 0.14 | 0.02 | 0.57 | 0.55 | 0.55 |
| 0.6 | 0.58 | 0.59 | 0.59 | 0.59 | 0.54 | 0.62 | 0.56 | 0.41 | 0.16 | 0.01 | 0 | 0.57 | 0.54 | 0.55 |
| 0.59 | 0.57 | 0.59 | 0.59 | 0.58 | 0.54 | 0.62 | 0.56 | 0.43 | 0.15 | 0 | 0 | 0.56 | 0.55 | 0.54 |
| 0.59 | 0.58 | 0.59 | 0.6 | 0.58 | 0.52 | 0.61 | 0.56 | 0.46 | 0.49 | 0.13 | 0.02 | 0.57 | 0.56 | 0.54 |
| 0.58 | 0.58 | 0.58 | 0.6 | 0.57 | 0.51 | 0.61 | 0.55 | 0.47 | 0.54 | 0.3 | 0.08 | 0.57 | 0.55 | 0.54 |
| 0.6 | 0.58 | 0.58 | 0.59 | 0.56 | 0.48 | 0.61 | 0.54 | 0.48 | 0.57 | 0.4 | 0.17 | 0.57 | 0.55 | 0.53 |
| 0.6 | 0.58 | 0.57 | 0.6 | 0.56 | 0.46 | 0.61 | 0.55 | 0.51 | 0.56 | 0.46 | 0.25 | 0.57 | 0.56 | 0.53 |
| 0.6 | 0.58 | 0.58 | 0.6 | 0.55 | 0.42 | 0.63 | 0.58 | 0.52 | 0.58 | 0.48 | 0.33 | 0.57 | 0.54 | 0.53 |
| 0.6 | 0.57 | 0.57 | 0.59 | 0.54 | 0.38 | 0.62 | 0.59 | 0.54 | 0.58 | 0.5 | 0.37 | 0.57 | 0.54 | 0.52 |
| 0.61 | 0.57 | 0.57 | 0.59 | 0.5 | 0.31 | 0.62 | 0.59 | 0.55 | 0.58 | 0.52 | 0.41 | 0.57 | 0.54 | 0.51 |
| 0.6 | 0.58 | 0.57 | 0.58 | 0.45 | 0.23 | 0.61 | 0.6 | 0.56 | 0.59 | 0.53 | 0.45 | 0.56 | 0.54 | 0.5 |
| 0.58 | 0.57 | 0.56 | 0.57 | 0.38 | 0.13 | 0.61 | 0.6 | 0.56 | 0.6 | 0.53 | 0.47 | 0.56 | 0.54 | 0.48 |
| 0.59 | 0.58 | 0.55 | 0.53 | 0.22 | 0.04 | 0.63 | 0.6 | 0.58 | 0.58 | 0.54 | 0.48 | 0.57 | 0.53 | 0.45 |
| 0.61 | 0.57 | 0.54 | 0.37 | 0.04 | 0.01 | 0.63 | 0.6 | 0.58 | 0.59 | 0.54 | 0.51 | 0.57 | 0.52 | 0.42 |
| 0.6 | 0.56 | 0.53 | 0.02 | 0.01 | 0 | 0.62 | 0.61 | 0.58 | 0.59 | 0.56 | 0.52 | 0.56 | 0.5 | 0.36 |
| 0.59 | 0.56 | 0.52 | 0.43 | 0.07 | 0.01 | 0.62 | 0.61 | 0.59 | 0.58 | 0.56 | 0.52 | 0.55 | 0.46 | 0.27 |
| 0.6 | 0.55 | 0.5 | 0.54 | 0.27 | 0.06 | 0.62 | 0.6 | 0.59 | 0.6 | 0.56 | 0.53 | 0.54 | 0.34 | 0.16 |
| 0.6 | 0.55 | 0.49 | 0.57 | 0.39 | 0.15 | 0.61 | 0.6 | 0.59 | 0.59 | 0.56 | 0.53 | 0.43 | 0.13 | 0.03 |
| 0.59 | 0.55 | 0.45 | 0.59 | 0.45 | 0.23 | 0.63 | 0.6 | 0.59 | 0.59 | 0.56 | 0.55 | 0.15 | 0.03 | 0.02 |
| 0.6 | 0.53 | 0.42 | 0.57 | 0.48 | 0.3 | 0.62 | 0.61 | 0.6 | 0.59 | 0.56 | 0.54 | 0.5 | 0.25 | 0.1 |
| 0.6 | 0.52 | 0.38 | 0.58 | 0.5 | 0.37 | 0.63 | 0.62 | 0.6 | 0.59 | 0.56 | 0.56 | 0.54 | 0.41 | 0.22 |
| 0.6 | 0.49 | 0.32 | 0.58 | 0.53 | 0.41 | 0.62 | 0.62 | 0.6 | 0.59 | 0.57 | 0.56 | 0.56 | 0.47 | 0.33 |
| 0.59 | 0.47 | 0.24 | 0.59 | 0.55 | 0.45 | 0.61 | 0.62 | 0.61 | 0.59 | 0.55 | 0.56 | 0.56 | 0.51 | 0.4 |
| 0.58 | 0.4 | 0.16 | 0.6 | 0.57 | 0.48 | 0.61 | 0.61 | 0.61 | 0.58 | 0.55 | 0.56 | 0.57 | 0.52 | 0.44 |
| 0.55 | 0.28 | 0.07 | 0.6 | 0.57 | 0.51 | 0.62 | 0.61 | 0.6 | 0.58 | 0.56 | 0.56 | 0.57 | 0.53 | 0.47 |
| 0.49 | 0.13 | 0.01 | 0.59 | 0.58 | 0.52 | 0.61 | 0.61 | 0.61 | 0.59 | 0.57 | 0.56 | 0.57 | 0.54 | 0.49 |
| 0.21 | 0 | 0 | 0.59 | 0.59 | 0.54 | 0.61 | 0.6 | 0.6 | 0.58 | 0.56 | 0.57 | 0.57 | 0.54 | 0.5 |
| 0.06 | 0.01 | 0.01 | 0.6 | 0.59 | 0.54 | 0.63 | 0.61 | 0.61 | 0.58 | 0.56 | 0.56 | 0.56 | 0.55 | 0.51 |
| 0.46 | 0.08 | 0.01 | 0.59 | 0.59 | 0.56 | 0.6 | 0.6 | 0.61 | 0.59 | 0.57 | 0.57 | 0.58 | 0.54 | 0.52 |
| 0.55 | 0.25 | 0.05 | 0.6 | 0.58 | 0.56 | 0.61 | 0.61 | 0.61 | 0.58 | 0.55 | 0.58 | 0.57 | 0.54 | 0.52 |
| 0.57 | 0.37 | 0.13 | 0.6 | 0.59 | 0.56 | 0.62 | 0.61 | 0.61 | 0.58 | 0.56 | 0.57 | 0.58 | 0.55 | 0.53 |
| 0.59 | 0.44 | 0.22 | 0.59 | 0.59 | 0.57 | 0.61 | 0.61 | 0.61 | 0.6 | 0.56 | 0.57 | 0.57 | 0.55 | 0.54 |
| 0.58 | 0.49 | 0.3 | 0.6 | 0.59 | 0.57 | 0.63 | 0.62 | 0.62 | 0.58 | 0.57 | 0.57 | 0.57 | 0.56 | 0.54 |
| 0.59 | 0.51 | 0.35 | 0.61 | 0.59 | 0.58 | 0.61 | 0.61 | 0.61 | 0.57 | 0.57 | 0.57 | 0.56 | 0.55 | 0.54 |
| 0.58 | 0.52 | 0.41 | 0.6 | 0.6 | 0.58 | 0.61 | 0.6 | 0.6 | 0.6 | 0.57 | 0.57 | 0.57 | 0.54 | 0.55 |
| 0.6 | 0.54 | 0.44 | 0.6 | 0.59 | 0.58 | 0.62 | 0.61 | 0.59 | 0.6 | 0.57 | 0.58 | 0.58 | 0.55 | 0.55 |
| 0.59 | 0.54 | 0.46 | 0.6 | 0.59 | 0.59 | 0.62 | 0.6 | 0.6 | 0.59 | 0.56 | 0.58 | 0.57 | 0.55 | 0.54 |
| 0.6 | 0.55 | 0.49 | 0.6 | 0.6 | 0.59 | 0.62 | 0.62 | 0.61 | 0.59 | 0.56 | 0.58 | 0.58 | 0.55 | 0.55 |
| 0.6 | 0.56 | 0.5 | 0.59 | 0.6 | 0.59 | 0.61 | 0.61 | 0.61 | 0.58 | 0.56 | 0.57 | 0.58 | 0.55 | 0.55 |
| 0.6 | 0.56 | 0.53 | 0.6 | 0.59 | 0.59 | 0.62 | 0.61 | 0.62 | 0.59 | 0.55 | 0.57 | 0.57 | 0.55 | 0.55 |
| 0.59 | 0.57 | 0.54 | 0.6 | 0.6 | 0.59 | 0.61 | 0.62 | 0.61 | 0.6 | 0.56 | 0.58 | 0.58 | 0.56 | 0.56 |
| 0.61 | 0.56 | 0.54 | 0.59 | 0.59 | 0.59 | 0.61 | 0.62 | 0.61 | 0.59 | 0.57 | 0.58 | 0.56 | 0.55 | 0.56 |
| 0.59 | 0.57 | 0.56 | 0.59 | 0.59 | 0.59 | 0.62 | 0.63 | 0.62 | 0.59 | 0.56 | 0.58 | 0.57 | 0.55 | 0.55 |
| 0.6 | 0.56 | 0.57 | 0.6 | 0.59 | 0.6 | 0.61 | 0.61 | 0.62 | 0.6 | 0.56 | 0.57 | 0.57 | 0.55 | 0.56 |
| 0.6 | 0.57 | 0.56 | 0.59 | 0.6 | 0.6 | 0.62 | 0.6 | 0.62 | 0.59 | 0.57 | 0.56 | 0.58 | 0.55 | 0.56 |
| 0.6 | 0.57 | 0.57 | 0.6 | 0.6 | 0.6 | 0.62 | 0.6 | 0.62 | 0.59 | 0.56 | 0.56 | 0.56 | 0.56 | 0.55 |

| 10HzL333 | 30HzL333 | 60HzL333 | 10HzG335 | 30HzG335 | 60HzG335 | 10HzT338 | 30HzT338 | 60HzT338 | 10Hz S340 | 30Hz S340 | 60Hz S340 | 10Hz F341 | 30Hz F341 | 60Hz F341 |
| --- | --- | --- | --- | --- | --- | --- | --- | --- | --- | --- | --- | --- | --- | --- |
| 0.58 | 0.59 | 0.59 | 0.58 | 0.51 | 0.47 | 0.59 | 0.54 | 0.52 | 0.58 | 0.6 | 0.61 | 0.56 | 0.58 | 0.58 |
| 0.59 | 0.59 | 0.6 | 0.56 | 0.52 | 0.48 | 0.6 | 0.54 | 0.51 | 0.59 | 0.59 | 0.62 | 0.59 | 0.6 | 0.59 |
| 0.59 | 0.58 | 0.59 | 0.57 | 0.51 | 0.45 | 0.59 | 0.56 | 0.49 | 0.59 | 0.6 | 0.6 | 0.58 | 0.59 | 0.59 |
| 0.57 | 0.58 | 0.6 | 0.57 | 0.52 | 0.44 | 0.58 | 0.55 | 0.49 | 0.58 | 0.59 | 0.61 | 0.62 | 0.58 | 0.6 |
| 0.59 | 0.6 | 0.6 | 0.56 | 0.5 | 0.41 | 0.58 | 0.53 | 0.45 | 0.58 | 0.6 | 0.6 | 0.6 | 0.58 | 0.59 |
| 0.58 | 0.58 | 0.6 | 0.55 | 0.48 | 0.37 | 0.57 | 0.52 | 0.42 | 0.59 | 0.6 | 0.61 | 0.57 | 0.58 | 0.58 |
| 0.59 | 0.58 | 0.6 | 0.57 | 0.48 | 0.32 | 0.58 | 0.51 | 0.39 | 0.58 | 0.59 | 0.61 | 0.61 | 0.57 | 0.59 |
| 0.58 | 0.59 | 0.6 | 0.54 | 0.45 | 0.27 | 0.57 | 0.5 | 0.33 | 0.58 | 0.58 | 0.6 | 0.59 | 0.57 | 0.6 |
| 0.58 | 0.58 | 0.59 | 0.58 | 0.41 | 0.2 | 0.57 | 0.47 | 0.25 | 0.57 | 0.6 | 0.6 | 0.58 | 0.59 | 0.59 |
| 0.59 | 0.59 | 0.59 | 0.53 | 0.35 | 0.12 | 0.57 | 0.41 | 0.18 | 0.6 | 0.6 | 0.61 | 0.6 | 0.6 | 0.59 |
| 0.59 | 0.59 | 0.59 | 0.53 | 0.27 | 0.06 | 0.55 | 0.3 | 0.08 | 0.58 | 0.61 | 0.61 | 0.6 | 0.58 | 0.58 |
| 0.58 | 0.58 | 0.58 | 0.46 | 0.1 | 0.01 | 0.49 | 0.15 | 0.02 | 0.57 | 0.59 | 0.6 | 0.57 | 0.59 | 0.59 |
| 0.58 | 0.58 | 0.59 | 0.2 | 0 | -0.01 | 0.28 | 0.02 | 0.01 | 0.58 | 0.59 | 0.6 | 0.59 | 0.58 | 0.57 |
| 0.59 | 0.58 | 0.59 | 0.04 | 0 | 0 | 0.04 | 0 | 0 | 0.58 | 0.58 | 0.6 | 0.62 | 0.57 | 0.57 |
| 0.58 | 0.59 | 0.59 | 0.41 | 0.04 | 0 | 0.43 | 0.06 | 0.01 | 0.59 | 0.59 | 0.6 | 0.59 | 0.57 | 0.59 |
| 0.58 | 0.58 | 0.59 | 0.52 | 0.17 | 0.02 | 0.53 | 0.23 | 0.04 | 0.57 | 0.59 | 0.58 | 0.59 | 0.59 | 0.59 |
| 0.58 | 0.57 | 0.59 | 0.56 | 0.31 | 0.09 | 0.56 | 0.36 | 0.11 | 0.58 | 0.59 | 0.58 | 0.58 | 0.58 | 0.58 |
| 0.58 | 0.58 | 0.59 | 0.54 | 0.4 | 0.17 | 0.57 | 0.43 | 0.21 | 0.58 | 0.59 | 0.59 | 0.59 | 0.56 | 0.59 |
| 0.59 | 0.58 | 0.58 | 0.56 | 0.45 | 0.26 | 0.57 | 0.46 | 0.28 | 0.58 | 0.58 | 0.59 | 0.58 | 0.58 | 0.59 |
| 0.6 | 0.58 | 0.58 | 0.56 | 0.48 | 0.32 | 0.57 | 0.49 | 0.34 | 0.61 | 0.57 | 0.59 | 0.6 | 0.6 | 0.57 |
| 0.59 | 0.58 | 0.57 | 0.57 | 0.49 | 0.36 | 0.58 | 0.51 | 0.38 | 0.57 | 0.58 | 0.58 | 0.59 | 0.59 | 0.57 |
| 0.59 | 0.58 | 0.57 | 0.57 | 0.5 | 0.39 | 0.58 | 0.52 | 0.41 | 0.6 | 0.58 | 0.58 | 0.59 | 0.58 | 0.58 |
| 0.59 | 0.58 | 0.57 | 0.56 | 0.5 | 0.41 | 0.58 | 0.53 | 0.43 | 0.58 | 0.58 | 0.56 | 0.58 | 0.56 | 0.57 |
| 0.59 | 0.58 | 0.56 | 0.56 | 0.52 | 0.45 | 0.58 | 0.52 | 0.44 | 0.57 | 0.59 | 0.57 | 0.56 | 0.57 | 0.58 |
| 0.59 | 0.58 | 0.56 | 0.59 | 0.51 | 0.47 | 0.59 | 0.51 | 0.46 | 0.58 | 0.59 | 0.56 | 0.59 | 0.58 | 0.56 |
| 0.58 | 0.57 | 0.56 | 0.59 | 0.52 | 0.48 | 0.58 | 0.52 | 0.48 | 0.59 | 0.59 | 0.55 | 0.61 | 0.57 | 0.56 |
| 0.58 | 0.58 | 0.55 | 0.57 | 0.51 | 0.49 | 0.58 | 0.54 | 0.49 | 0.59 | 0.57 | 0.55 | 0.6 | 0.58 | 0.54 |
| 0.58 | 0.57 | 0.55 | 0.56 | 0.51 | 0.5 | 0.57 | 0.56 | 0.5 | 0.58 | 0.57 | 0.54 | 0.58 | 0.58 | 0.54 |
| 0.59 | 0.56 | 0.53 | 0.58 | 0.55 | 0.52 | 0.58 | 0.56 | 0.52 | 0.59 | 0.56 | 0.53 | 0.59 | 0.56 | 0.54 |
| 0.58 | 0.57 | 0.51 | 0.57 | 0.54 | 0.51 | 0.58 | 0.55 | 0.53 | 0.6 | 0.56 | 0.5 | 0.58 | 0.56 | 0.52 |
| 0.59 | 0.56 | 0.5 | 0.59 | 0.55 | 0.53 | 0.58 | 0.57 | 0.54 | 0.59 | 0.57 | 0.49 | 0.58 | 0.57 | 0.52 |
| 0.57 | 0.55 | 0.48 | 0.58 | 0.54 | 0.51 | 0.58 | 0.56 | 0.53 | 0.57 | 0.55 | 0.46 | 0.58 | 0.56 | 0.5 |
| 0.58 | 0.55 | 0.45 | 0.56 | 0.54 | 0.52 | 0.59 | 0.55 | 0.55 | 0.58 | 0.55 | 0.41 | 0.6 | 0.56 | 0.48 |
| 0.58 | 0.54 | 0.42 | 0.57 | 0.54 | 0.53 | 0.57 | 0.55 | 0.55 | 0.57 | 0.53 | 0.37 | 0.57 | 0.55 | 0.47 |
| 0.58 | 0.52 | 0.38 | 0.58 | 0.55 | 0.54 | 0.59 | 0.56 | 0.57 | 0.57 | 0.47 | 0.29 | 0.58 | 0.54 | 0.42 |
| 0.57 | 0.5 | 0.32 | 0.56 | 0.54 | 0.55 | 0.58 | 0.56 | 0.57 | 0.56 | 0.43 | 0.22 | 0.58 | 0.52 | 0.38 |
| 0.58 | 0.47 | 0.25 | 0.57 | 0.54 | 0.55 | 0.58 | 0.57 | 0.56 | 0.57 | 0.37 | 0.13 | 0.57 | 0.49 | 0.33 |
| 0.56 | 0.41 | 0.16 | 0.57 | 0.55 | 0.53 | 0.58 | 0.56 | 0.57 | 0.52 | 0.25 | 0.05 | 0.55 | 0.48 | 0.25 |
| 0.54 | 0.3 | 0.08 | 0.57 | 0.54 | 0.53 | 0.57 | 0.57 | 0.58 | 0.46 | 0.09 | 0.01 | 0.54 | 0.39 | 0.15 |
| 0.49 | 0.14 | 0.01 | 0.57 | 0.55 | 0.53 | 0.58 | 0.56 | 0.57 | 0.05 | 0 | 0 | 0.54 | 0.27 | 0.06 |
| 0.18 | 0.01 | 0 | 0.57 | 0.55 | 0.55 | 0.58 | 0.58 | 0.57 | 0.23 | 0.02 | 0.01 | 0.45 | 0.1 | 0.01 |
| 0.1 | 0.01 | 0 | 0.54 | 0.56 | 0.54 | 0.59 | 0.57 | 0.58 | 0.49 | 0.14 | 0.01 | 0.04 | 0.01 | 0 |
| 0.45 | 0.09 | 0.01 | 0.58 | 0.56 | 0.55 | 0.58 | 0.57 | 0.58 | 0.54 | 0.32 | 0.07 | 0.26 | 0.02 | 0 |
| 0.53 | 0.27 | 0.06 | 0.57 | 0.56 | 0.56 | 0.58 | 0.58 | 0.59 | 0.56 | 0.42 | 0.18 | 0.51 | 0.17 | 0.02 |
| 0.56 | 0.38 | 0.14 | 0.55 | 0.53 | 0.56 | 0.58 | 0.57 | 0.58 | 0.56 | 0.49 | 0.27 | 0.56 | 0.34 | 0.09 |
| 0.56 | 0.44 | 0.23 | 0.56 | 0.55 | 0.57 | 0.58 | 0.58 | 0.58 | 0.59 | 0.52 | 0.34 | 0.55 | 0.45 | 0.17 |
| 0.58 | 0.49 | 0.31 | 0.56 | 0.55 | 0.57 | 0.58 | 0.57 | 0.58 | 0.59 | 0.53 | 0.4 | 0.58 | 0.48 | 0.26 |
| 0.57 | 0.51 | 0.37 | 0.57 | 0.54 | 0.54 | 0.58 | 0.57 | 0.58 | 0.58 | 0.54 | 0.42 | 0.57 | 0.51 | 0.32 |
| 0.59 | 0.53 | 0.41 | 0.58 | 0.54 | 0.56 | 0.58 | 0.56 | 0.58 | 0.58 | 0.56 | 0.48 | 0.57 | 0.54 | 0.38 |
| 0.58 | 0.54 | 0.45 | 0.57 | 0.54 | 0.56 | 0.58 | 0.57 | 0.59 | 0.59 | 0.56 | 0.5 | 0.59 | 0.54 | 0.43 |
| 0.59 | 0.55 | 0.48 | 0.57 | 0.56 | 0.56 | 0.58 | 0.57 | 0.58 | 0.59 | 0.57 | 0.52 | 0.58 | 0.57 | 0.45 |
| 0.59 | 0.56 | 0.5 | 0.57 | 0.54 | 0.57 | 0.58 | 0.56 | 0.58 | 0.59 | 0.58 | 0.53 | 0.59 | 0.57 | 0.47 |
| 0.59 | 0.57 | 0.52 | 0.57 | 0.54 | 0.56 | 0.58 | 0.57 | 0.58 | 0.59 | 0.58 | 0.54 | 0.59 | 0.55 | 0.49 |
| 0.58 | 0.56 | 0.53 | 0.57 | 0.54 | 0.56 | 0.58 | 0.57 | 0.58 | 0.58 | 0.59 | 0.54 | 0.58 | 0.56 | 0.52 |
| 0.59 | 0.58 | 0.53 | 0.58 | 0.55 | 0.55 | 0.58 | 0.56 | 0.58 | 0.59 | 0.58 | 0.55 | 0.59 | 0.56 | 0.53 |
| 0.59 | 0.56 | 0.54 | 0.58 | 0.54 | 0.54 | 0.58 | 0.56 | 0.57 | 0.58 | 0.59 | 0.57 | 0.58 | 0.56 | 0.53 |
| 0.58 | 0.57 | 0.55 | 0.57 | 0.54 | 0.55 | 0.58 | 0.57 | 0.57 | 0.57 | 0.59 | 0.56 | 0.58 | 0.57 | 0.54 |
| 0.58 | 0.58 | 0.56 | 0.56 | 0.54 | 0.55 | 0.58 | 0.57 | 0.59 | 0.59 | 0.58 | 0.58 | 0.57 | 0.56 | 0.54 |
| 0.58 | 0.56 | 0.57 | 0.58 | 0.56 | 0.56 | 0.59 | 0.57 | 0.59 | 0.58 | 0.58 | 0.59 | 0.56 | 0.59 | 0.56 |
| 0.58 | 0.58 | 0.56 | 0.56 | 0.55 | 0.55 | 0.57 | 0.57 | 0.59 | 0.58 | 0.59 | 0.59 | 0.59 | 0.6 | 0.56 |
| 0.58 | 0.58 | 0.58 | 0.55 | 0.54 | 0.56 | 0.58 | 0.57 | 0.59 | 0.56 | 0.59 | 0.59 | 0.6 | 0.58 | 0.55 |
| 0.58 | 0.58 | 0.58 | 0.55 | 0.55 | 0.56 | 0.58 | 0.55 | 0.59 | 0.59 | 0.6 | 0.58 | 0.6 | 0.58 | 0.57 |
| 0.59 | 0.57 | 0.57 | 0.57 | 0.55 | 0.57 | 0.57 | 0.57 | 0.59 | 0.58 | 0.59 | 0.58 | 0.61 | 0.59 | 0.57 |

| 10HzD343 | 30HzD343 | 60HzD343 | 10HzS346 | 30HzS346 | 60HzS346 | 10HzA347 | 30HzA347 | 60HzA347 | 10HzA349 | 30Hz  A349 | 60HzA349 | 10HzA350 | 30HzA350 | 60HzA350 |
| --- | --- | --- | --- | --- | --- | --- | --- | --- | --- | --- | --- | --- | --- | --- |
| 0.55 | 0.54 | 0.55 | 0.59 | 0.58 | 0.57 | 0.57 | 0.57 | 0.57 | 0.55 | 0.57 | 0.57 | 0.59 | 0.56 | 0.59 |
| 0.58 | 0.55 | 0.57 | 0.57 | 0.58 | 0.58 | 0.58 | 0.55 | 0.57 | 0.56 | 0.54 | 0.59 | 0.56 | 0.56 | 0.57 |
| 0.56 | 0.56 | 0.55 | 0.59 | 0.58 | 0.58 | 0.55 | 0.58 | 0.56 | 0.53 | 0.54 | 0.59 | 0.59 | 0.56 | 0.58 |
| 0.6 | 0.55 | 0.55 | 0.6 | 0.57 | 0.57 | 0.6 | 0.54 | 0.57 | 0.56 | 0.53 | 0.58 | 0.6 | 0.55 | 0.56 |
| 0.58 | 0.56 | 0.57 | 0.59 | 0.58 | 0.56 | 0.6 | 0.53 | 0.59 | 0.55 | 0.55 | 0.58 | 0.54 | 0.56 | 0.57 |
| 0.59 | 0.55 | 0.55 | 0.59 | 0.58 | 0.57 | 0.58 | 0.55 | 0.56 | 0.55 | 0.55 | 0.57 | 0.61 | 0.56 | 0.58 |
| 0.56 | 0.56 | 0.56 | 0.6 | 0.57 | 0.57 | 0.56 | 0.55 | 0.59 | 0.59 | 0.57 | 0.57 | 0.58 | 0.55 | 0.57 |
| 0.57 | 0.55 | 0.56 | 0.6 | 0.58 | 0.56 | 0.56 | 0.59 | 0.6 | 0.55 | 0.54 | 0.57 | 0.58 | 0.6 | 0.57 |
| 0.6 | 0.57 | 0.53 | 0.59 | 0.56 | 0.56 | 0.59 | 0.57 | 0.58 | 0.54 | 0.55 | 0.57 | 0.54 | 0.55 | 0.57 |
| 0.65 | 0.54 | 0.55 | 0.59 | 0.57 | 0.55 | 0.59 | 0.52 | 0.57 | 0.57 | 0.55 | 0.58 | 0.6 | 0.57 | 0.58 |
| 0.61 | 0.55 | 0.55 | 0.6 | 0.57 | 0.55 | 0.61 | 0.6 | 0.58 | 0.53 | 0.55 | 0.58 | 0.58 | 0.56 | 0.56 |
| 0.59 | 0.55 | 0.56 | 0.59 | 0.56 | 0.54 | 0.59 | 0.54 | 0.56 | 0.56 | 0.54 | 0.55 | 0.59 | 0.57 | 0.57 |
| 0.59 | 0.56 | 0.55 | 0.58 | 0.57 | 0.53 | 0.57 | 0.56 | 0.56 | 0.57 | 0.54 | 0.56 | 0.56 | 0.56 | 0.56 |
| 0.56 | 0.53 | 0.54 | 0.59 | 0.56 | 0.52 | 0.6 | 0.58 | 0.57 | 0.55 | 0.54 | 0.54 | 0.58 | 0.59 | 0.55 |
| 0.6 | 0.57 | 0.55 | 0.58 | 0.56 | 0.51 | 0.6 | 0.53 | 0.6 | 0.54 | 0.54 | 0.55 | 0.55 | 0.54 | 0.54 |
| 0.6 | 0.54 | 0.52 | 0.59 | 0.56 | 0.48 | 0.55 | 0.54 | 0.56 | 0.54 | 0.56 | 0.57 | 0.57 | 0.53 | 0.56 |
| 0.57 | 0.53 | 0.52 | 0.58 | 0.55 | 0.47 | 0.58 | 0.57 | 0.56 | 0.58 | 0.55 | 0.55 | 0.59 | 0.56 | 0.55 |
| 0.56 | 0.56 | 0.51 | 0.59 | 0.53 | 0.44 | 0.6 | 0.55 | 0.56 | 0.55 | 0.55 | 0.54 | 0.57 | 0.56 | 0.55 |
| 0.58 | 0.52 | 0.52 | 0.59 | 0.53 | 0.4 | 0.56 | 0.56 | 0.54 | 0.57 | 0.54 | 0.55 | 0.59 | 0.55 | 0.55 |
| 0.6 | 0.54 | 0.51 | 0.59 | 0.5 | 0.36 | 0.55 | 0.58 | 0.54 | 0.54 | 0.54 | 0.53 | 0.58 | 0.54 | 0.55 |
| 0.59 | 0.53 | 0.49 | 0.58 | 0.47 | 0.29 | 0.59 | 0.57 | 0.55 | 0.57 | 0.54 | 0.53 | 0.6 | 0.54 | 0.54 |
| 0.6 | 0.53 | 0.46 | 0.58 | 0.43 | 0.21 | 0.64 | 0.59 | 0.53 | 0.59 | 0.51 | 0.52 | 0.6 | 0.57 | 0.54 |
| 0.6 | 0.53 | 0.45 | 0.56 | 0.36 | 0.12 | 0.57 | 0.54 | 0.54 | 0.54 | 0.51 | 0.54 | 0.6 | 0.54 | 0.54 |
| 0.59 | 0.52 | 0.41 | 0.54 | 0.22 | 0.04 | 0.58 | 0.56 | 0.55 | 0.55 | 0.53 | 0.51 | 0.6 | 0.56 | 0.52 |
| 0.56 | 0.49 | 0.32 | 0.39 | 0.06 | 0 | 0.54 | 0.51 | 0.54 | 0.55 | 0.52 | 0.5 | 0.58 | 0.56 | 0.51 |
| 0.57 | 0.42 | 0.24 | 0.04 | 0 | 0 | 0.56 | 0.53 | 0.53 | 0.55 | 0.55 | 0.48 | 0.6 | 0.55 | 0.52 |
| 0.58 | 0.29 | 0.13 | 0.35 | 0.03 | 0 | 0.59 | 0.54 | 0.55 | 0.55 | 0.53 | 0.46 | 0.58 | 0.51 | 0.49 |
| 0.28 | 0.13 | 0.04 | 0.51 | 0.19 | 0.03 | 0.56 | 0.54 | 0.51 | 0.58 | 0.53 | 0.45 | 0.58 | 0.56 | 0.49 |
| 0.1 | 0.03 | -0.01 | 0.55 | 0.35 | 0.11 | 0.59 | 0.55 | 0.47 | 0.56 | 0.49 | 0.42 | 0.59 | 0.53 | 0.45 |
| 0.27 | 0.04 | -0.01 | 0.57 | 0.43 | 0.21 | 0.59 | 0.57 | 0.5 | 0.53 | 0.49 | 0.37 | 0.58 | 0.54 | 0.42 |
| 0.52 | 0.24 | 0.05 | 0.58 | 0.48 | 0.28 | 0.61 | 0.54 | 0.44 | 0.55 | 0.46 | 0.33 | 0.6 | 0.5 | 0.37 |
| 0.59 | 0.37 | 0.2 | 0.57 | 0.5 | 0.35 | 0.62 | 0.49 | 0.45 | 0.55 | 0.44 | 0.25 | 0.57 | 0.48 | 0.32 |
| 0.61 | 0.46 | 0.26 | 0.58 | 0.53 | 0.4 | 0.6 | 0.52 | 0.43 | 0.51 | 0.37 | 0.17 | 0.57 | 0.46 | 0.24 |
| 0.61 | 0.5 | 0.36 | 0.58 | 0.54 | 0.44 | 0.56 | 0.52 | 0.39 | 0.54 | 0.31 | 0.07 | 0.55 | 0.38 | 0.17 |
| 0.61 | 0.52 | 0.42 | 0.59 | 0.55 | 0.47 | 0.57 | 0.48 | 0.36 | 0.45 | 0.14 | 0.01 | 0.55 | 0.28 | 0.06 |
| 0.58 | 0.54 | 0.45 | 0.59 | 0.55 | 0.48 | 0.58 | 0.47 | 0.29 | 0.21 | 0.00 | 0 | 0.47 | 0.11 | 0.01 |
| 0.6 | 0.52 | 0.49 | 0.6 | 0.56 | 0.51 | 0.58 | 0.46 | 0.23 | 0.07 | -0.01 | 0.02 | 0.2 | 0.01 | 0 |
| 0.57 | 0.54 | 0.49 | 0.58 | 0.57 | 0.52 | 0.57 | 0.37 | 0.16 | 0.44 | 0.07 | 0.02 | 0.09 | 0 | -0.01 |
| 0.59 | 0.56 | 0.49 | 0.59 | 0.57 | 0.52 | 0.53 | 0.27 | 0.08 | 0.5 | 0.25 | 0.07 | 0.46 | 0.1 | 0 |
| 0.61 | 0.55 | 0.52 | 0.6 | 0.57 | 0.54 | 0.55 | 0.16 | 0.03 | 0.53 | 0.34 | 0.15 | 0.54 | 0.24 | 0.04 |
| 0.6 | 0.55 | 0.52 | 0.58 | 0.58 | 0.55 | 0.27 | 0.01 | 0.01 | 0.53 | 0.42 | 0.23 | 0.55 | 0.37 | 0.14 |
| 0.59 | 0.55 | 0.53 | 0.59 | 0.56 | 0.55 | 0.01 | 0 | 0.01 | 0.53 | 0.46 | 0.3 | 0.57 | 0.43 | 0.2 |
| 0.59 | 0.54 | 0.54 | 0.59 | 0.57 | 0.56 | 0.33 | 0.01 | 0.03 | 0.57 | 0.47 | 0.33 | 0.57 | 0.43 | 0.27 |
| 0.56 | 0.53 | 0.54 | 0.59 | 0.58 | 0.56 | 0.53 | 0.18 | 0.02 | 0.59 | 0.49 | 0.38 | 0.58 | 0.49 | 0.35 |
| 0.58 | 0.56 | 0.56 | 0.58 | 0.58 | 0.56 | 0.54 | 0.32 | 0.07 | 0.53 | 0.48 | 0.4 | 0.59 | 0.5 | 0.39 |
| 0.61 | 0.56 | 0.56 | 0.59 | 0.58 | 0.57 | 0.6 | 0.41 | 0.16 | 0.52 | 0.47 | 0.44 | 0.57 | 0.53 | 0.42 |
| 0.62 | 0.58 | 0.56 | 0.59 | 0.57 | 0.57 | 0.53 | 0.41 | 0.25 | 0.55 | 0.51 | 0.45 | 0.57 | 0.53 | 0.45 |
| 0.59 | 0.55 | 0.56 | 0.59 | 0.58 | 0.58 | 0.56 | 0.43 | 0.3 | 0.54 | 0.52 | 0.49 | 0.59 | 0.51 | 0.48 |
| 0.59 | 0.55 | 0.55 | 0.57 | 0.57 | 0.57 | 0.54 | 0.47 | 0.34 | 0.56 | 0.53 | 0.49 | 0.59 | 0.54 | 0.5 |
| 0.59 | 0.54 | 0.56 | 0.6 | 0.57 | 0.58 | 0.59 | 0.5 | 0.39 | 0.54 | 0.53 | 0.51 | 0.57 | 0.52 | 0.49 |
| 0.57 | 0.54 | 0.56 | 0.6 | 0.58 | 0.57 | 0.59 | 0.53 | 0.42 | 0.57 | 0.54 | 0.53 | 0.57 | 0.58 | 0.5 |
| 0.63 | 0.56 | 0.56 | 0.58 | 0.58 | 0.58 | 0.58 | 0.53 | 0.45 | 0.57 | 0.55 | 0.53 | 0.61 | 0.55 | 0.51 |
| 0.59 | 0.54 | 0.56 | 0.59 | 0.58 | 0.59 | 0.62 | 0.55 | 0.48 | 0.55 | 0.56 | 0.54 | 0.6 | 0.55 | 0.53 |
| 0.58 | 0.55 | 0.58 | 0.59 | 0.57 | 0.58 | 0.6 | 0.52 | 0.49 | 0.57 | 0.54 | 0.52 | 0.55 | 0.56 | 0.54 |
| 0.59 | 0.53 | 0.55 | 0.59 | 0.57 | 0.59 | 0.6 | 0.53 | 0.5 | 0.57 | 0.53 | 0.55 | 0.57 | 0.56 | 0.54 |
| 0.59 | 0.55 | 0.59 | 0.58 | 0.58 | 0.58 | 0.58 | 0.56 | 0.52 | 0.56 | 0.53 | 0.56 | 0.6 | 0.55 | 0.56 |
| 0.62 | 0.53 | 0.56 | 0.59 | 0.57 | 0.59 | 0.58 | 0.55 | 0.52 | 0.57 | 0.54 | 0.55 | 0.57 | 0.56 | 0.54 |
| 0.59 | 0.57 | 0.59 | 0.58 | 0.58 | 0.58 | 0.58 | 0.56 | 0.54 | 0.55 | 0.55 | 0.56 | 0.58 | 0.57 | 0.57 |
| 0.59 | 0.55 | 0.57 | 0.58 | 0.58 | 0.58 | 0.6 | 0.54 | 0.55 | 0.57 | 0.54 | 0.55 | 0.58 | 0.57 | 0.54 |
| 0.58 | 0.55 | 0.56 | 0.59 | 0.57 | 0.58 | 0.6 | 0.54 | 0.53 | 0.57 | 0.55 | 0.56 | 0.58 | 0.56 | 0.56 |
| 0.57 | 0.58 | 0.57 | 0.59 | 0.58 | 0.59 | 0.56 | 0.55 | 0.54 | 0.54 | 0.54 | 0.57 | 0.59 | 0.56 | 0.56 |
| 0.6 | 0.54 | 0.57 | 0.6 | 0.58 | 0.59 | 0.55 | 0.53 | 0.56 | 0.56 | 0.55 | 0.56 | 0.6 | 0.55 | 0.57 |
| 0.55 | 0.54 | 0.57 | 0.59 | 0.58 | 0.59 | 0.56 | 0.56 | 0.55 | 0.53 | 0.55 | 0.56 | 0.59 | 0.55 | 0.57 |

| 10Hz I351 | 30Hz I351 | 60Hz I351 | 10HzD352 | 30HzD352 | 60HzD352 | 10Hz F354 | 30Hz F354 | 60Hz F354 | 10HzD355 | 30HzD355 | 60HzD355 | 10HzG356 | 30HzG356 | 60HzG356 |
| --- | --- | --- | --- | --- | --- | --- | --- | --- | --- | --- | --- | --- | --- | --- |
| 0.59 | 0.61 | 0.6 | 0.6 | 0.58 | 0.6 | 0.6 | 0.59 | 0.57 | 0.58 | 0.58 | 0.59 | 0.59 | 0.57 | 0.57 |
| 0.6 | 0.6 | 0.61 | 0.58 | 0.58 | 0.59 | 0.6 | 0.59 | 0.56 | 0.6 | 0.59 | 0.58 | 0.58 | 0.57 | 0.58 |
| 0.62 | 0.6 | 0.61 | 0.6 | 0.57 | 0.59 | 0.63 | 0.59 | 0.56 | 0.6 | 0.59 | 0.59 | 0.58 | 0.56 | 0.58 |
| 0.62 | 0.6 | 0.6 | 0.59 | 0.58 | 0.59 | 0.63 | 0.58 | 0.57 | 0.61 | 0.59 | 0.59 | 0.57 | 0.57 | 0.56 |
| 0.6 | 0.61 | 0.61 | 0.6 | 0.56 | 0.59 | 0.62 | 0.59 | 0.55 | 0.61 | 0.59 | 0.6 | 0.59 | 0.56 | 0.56 |
| 0.6 | 0.6 | 0.59 | 0.6 | 0.57 | 0.58 | 0.63 | 0.58 | 0.54 | 0.6 | 0.59 | 0.59 | 0.6 | 0.56 | 0.55 |
| 0.59 | 0.58 | 0.6 | 0.59 | 0.58 | 0.58 | 0.61 | 0.58 | 0.53 | 0.59 | 0.59 | 0.59 | 0.59 | 0.56 | 0.56 |
| 0.58 | 0.61 | 0.61 | 0.59 | 0.59 | 0.58 | 0.62 | 0.57 | 0.53 | 0.6 | 0.58 | 0.58 | 0.58 | 0.57 | 0.55 |
| 0.63 | 0.59 | 0.58 | 0.59 | 0.58 | 0.59 | 0.62 | 0.57 | 0.5 | 0.6 | 0.58 | 0.57 | 0.6 | 0.57 | 0.56 |
| 0.59 | 0.62 | 0.61 | 0.59 | 0.57 | 0.58 | 0.61 | 0.56 | 0.5 | 0.61 | 0.59 | 0.58 | 0.6 | 0.57 | 0.54 |
| 0.59 | 0.61 | 0.58 | 0.59 | 0.58 | 0.58 | 0.61 | 0.56 | 0.47 | 0.6 | 0.58 | 0.58 | 0.59 | 0.56 | 0.53 |
| 0.62 | 0.6 | 0.59 | 0.6 | 0.58 | 0.58 | 0.6 | 0.56 | 0.46 | 0.61 | 0.59 | 0.58 | 0.58 | 0.54 | 0.52 |
| 0.61 | 0.59 | 0.6 | 0.59 | 0.57 | 0.58 | 0.6 | 0.55 | 0.43 | 0.6 | 0.59 | 0.58 | 0.61 | 0.55 | 0.51 |
| 0.6 | 0.59 | 0.6 | 0.59 | 0.58 | 0.57 | 0.58 | 0.54 | 0.39 | 0.61 | 0.59 | 0.56 | 0.58 | 0.55 | 0.49 |
| 0.63 | 0.59 | 0.57 | 0.59 | 0.57 | 0.57 | 0.6 | 0.52 | 0.33 | 0.6 | 0.58 | 0.56 | 0.57 | 0.51 | 0.47 |
| 0.58 | 0.61 | 0.57 | 0.59 | 0.58 | 0.56 | 0.63 | 0.49 | 0.26 | 0.6 | 0.57 | 0.55 | 0.57 | 0.51 | 0.46 |
| 0.61 | 0.59 | 0.56 | 0.58 | 0.57 | 0.56 | 0.59 | 0.43 | 0.19 | 0.6 | 0.59 | 0.56 | 0.58 | 0.52 | 0.46 |
| 0.61 | 0.57 | 0.56 | 0.59 | 0.56 | 0.55 | 0.58 | 0.33 | 0.09 | 0.61 | 0.57 | 0.55 | 0.57 | 0.55 | 0.44 |
| 0.57 | 0.6 | 0.55 | 0.58 | 0.56 | 0.54 | 0.53 | 0.19 | 0.03 | 0.61 | 0.57 | 0.55 | 0.59 | 0.53 | 0.43 |
| 0.58 | 0.58 | 0.57 | 0.59 | 0.58 | 0.53 | 0.4 | 0.03 | 0.01 | 0.61 | 0.57 | 0.53 | 0.58 | 0.53 | 0.38 |
| 0.58 | 0.58 | 0.56 | 0.59 | 0.57 | 0.52 | 0.02 | 0 | 0.01 | 0.61 | 0.58 | 0.52 | 0.59 | 0.5 | 0.36 |
| 0.58 | 0.59 | 0.57 | 0.58 | 0.55 | 0.51 | 0.33 | 0.03 | 0 | 0.6 | 0.57 | 0.52 | 0.61 | 0.49 | 0.29 |
| 0.59 | 0.58 | 0.55 | 0.57 | 0.55 | 0.49 | 0.52 | 0.16 | 0.02 | 0.61 | 0.56 | 0.5 | 0.59 | 0.45 | 0.23 |
| 0.59 | 0.6 | 0.53 | 0.6 | 0.53 | 0.46 | 0.57 | 0.3 | 0.06 | 0.6 | 0.56 | 0.47 | 0.56 | 0.4 | 0.16 |
| 0.59 | 0.6 | 0.52 | 0.59 | 0.53 | 0.42 | 0.6 | 0.41 | 0.15 | 0.59 | 0.55 | 0.45 | 0.57 | 0.32 | 0.07 |
| 0.59 | 0.58 | 0.48 | 0.58 | 0.52 | 0.38 | 0.6 | 0.45 | 0.23 | 0.6 | 0.54 | 0.41 | 0.51 | 0.16 | 0.02 |
| 0.59 | 0.56 | 0.47 | 0.58 | 0.5 | 0.32 | 0.6 | 0.5 | 0.29 | 0.61 | 0.53 | 0.37 | 0.33 | 0.01 | 0 |
| 0.6 | 0.55 | 0.44 | 0.57 | 0.45 | 0.26 | 0.62 | 0.52 | 0.36 | 0.59 | 0.5 | 0.33 | 0.02 | 0 | 0 |
| 0.61 | 0.52 | 0.4 | 0.58 | 0.41 | 0.17 | 0.6 | 0.53 | 0.39 | 0.59 | 0.46 | 0.25 | 0.36 | 0.04 | 0 |
| 0.57 | 0.56 | 0.35 | 0.55 | 0.31 | 0.08 | 0.6 | 0.53 | 0.42 | 0.58 | 0.41 | 0.17 | 0.5 | 0.19 | 0.02 |
| 0.6 | 0.51 | 0.3 | 0.49 | 0.14 | 0.02 | 0.63 | 0.55 | 0.45 | 0.57 | 0.32 | 0.08 | 0.57 | 0.32 | 0.08 |
| 0.59 | 0.45 | 0.21 | 0.25 | 0.02 | 0 | 0.62 | 0.56 | 0.46 | 0.51 | 0.15 | 0.02 | 0.56 | 0.4 | 0.18 |
| 0.57 | 0.39 | 0.13 | 0.05 | 0 | 0 | 0.61 | 0.54 | 0.48 | 0.3 | 0.02 | 0 | 0.59 | 0.48 | 0.26 |
| 0.54 | 0.26 | 0.04 | 0.43 | 0.08 | 0 | 0.61 | 0.57 | 0.49 | 0 | 0.01 | 0 | 0.59 | 0.49 | 0.34 |
| 0.44 | 0.08 | 0.03 | 0.52 | 0.25 | 0.04 | 0.61 | 0.55 | 0.5 | 0.43 | 0.06 | 0 | 0.59 | 0.51 | 0.38 |
| 0.11 | 0.02 | 0.00 | 0.56 | 0.37 | 0.11 | 0.6 | 0.56 | 0.5 | 0.54 | 0.23 | 0.03 | 0.58 | 0.53 | 0.42 |
| 0.16 | 0.01 | 0 | 0.58 | 0.44 | 0.2 | 0.62 | 0.56 | 0.5 | 0.58 | 0.35 | 0.1 | 0.6 | 0.53 | 0.46 |
| 0.48 | 0.16 | 0.02 | 0.58 | 0.47 | 0.28 | 0.59 | 0.54 | 0.51 | 0.58 | 0.43 | 0.19 | 0.59 | 0.55 | 0.47 |
| 0.56 | 0.29 | 0.08 | 0.6 | 0.5 | 0.34 | 0.6 | 0.56 | 0.52 | 0.59 | 0.47 | 0.27 | 0.58 | 0.53 | 0.49 |
| 0.62 | 0.4 | 0.15 | 0.58 | 0.52 | 0.38 | 0.6 | 0.57 | 0.54 | 0.6 | 0.48 | 0.32 | 0.6 | 0.54 | 0.51 |
| 0.58 | 0.47 | 0.23 | 0.59 | 0.54 | 0.42 | 0.62 | 0.58 | 0.55 | 0.58 | 0.5 | 0.38 | 0.6 | 0.54 | 0.53 |
| 0.59 | 0.52 | 0.33 | 0.6 | 0.53 | 0.44 | 0.62 | 0.57 | 0.55 | 0.61 | 0.54 | 0.44 | 0.6 | 0.56 | 0.54 |
| 0.57 | 0.53 | 0.36 | 0.58 | 0.52 | 0.46 | 0.63 | 0.57 | 0.58 | 0.59 | 0.55 | 0.46 | 0.61 | 0.56 | 0.53 |
| 0.6 | 0.54 | 0.43 | 0.59 | 0.54 | 0.48 | 0.62 | 0.59 | 0.57 | 0.6 | 0.56 | 0.48 | 0.59 | 0.56 | 0.54 |
| 0.62 | 0.57 | 0.46 | 0.59 | 0.55 | 0.5 | 0.62 | 0.59 | 0.57 | 0.59 | 0.57 | 0.51 | 0.58 | 0.56 | 0.55 |
| 0.6 | 0.56 | 0.48 | 0.58 | 0.56 | 0.53 | 0.6 | 0.6 | 0.58 | 0.6 | 0.57 | 0.52 | 0.58 | 0.56 | 0.55 |
| 0.62 | 0.6 | 0.5 | 0.6 | 0.56 | 0.53 | 0.64 | 0.6 | 0.58 | 0.59 | 0.57 | 0.52 | 0.62 | 0.58 | 0.56 |
| 0.61 | 0.6 | 0.51 | 0.58 | 0.57 | 0.54 | 0.61 | 0.57 | 0.58 | 0.6 | 0.58 | 0.54 | 0.59 | 0.56 | 0.56 |
| 0.61 | 0.59 | 0.54 | 0.6 | 0.58 | 0.57 | 0.6 | 0.59 | 0.59 | 0.59 | 0.58 | 0.54 | 0.59 | 0.55 | 0.57 |
| 0.56 | 0.59 | 0.54 | 0.58 | 0.58 | 0.56 | 0.62 | 0.58 | 0.58 | 0.6 | 0.58 | 0.56 | 0.59 | 0.58 | 0.56 |
| 0.6 | 0.6 | 0.54 | 0.59 | 0.58 | 0.57 | 0.61 | 0.58 | 0.59 | 0.61 | 0.58 | 0.57 | 0.6 | 0.59 | 0.57 |
| 0.59 | 0.61 | 0.56 | 0.58 | 0.59 | 0.57 | 0.61 | 0.59 | 0.59 | 0.6 | 0.57 | 0.56 | 0.6 | 0.57 | 0.58 |
| 0.63 | 0.59 | 0.57 | 0.6 | 0.58 | 0.57 | 0.61 | 0.58 | 0.59 | 0.6 | 0.57 | 0.57 | 0.61 | 0.56 | 0.58 |
| 0.6 | 0.61 | 0.57 | 0.59 | 0.59 | 0.57 | 0.6 | 0.59 | 0.59 | 0.61 | 0.57 | 0.57 | 0.6 | 0.57 | 0.58 |
| 0.62 | 0.6 | 0.57 | 0.59 | 0.58 | 0.58 | 0.61 | 0.6 | 0.61 | 0.62 | 0.59 | 0.58 | 0.6 | 0.56 | 0.56 |
| 0.6 | 0.59 | 0.57 | 0.6 | 0.58 | 0.58 | 0.62 | 0.6 | 0.59 | 0.59 | 0.58 | 0.57 | 0.59 | 0.57 | 0.58 |
| 0.58 | 0.59 | 0.59 | 0.59 | 0.59 | 0.59 | 0.61 | 0.59 | 0.59 | 0.61 | 0.58 | 0.58 | 0.57 | 0.56 | 0.58 |
| 0.61 | 0.62 | 0.59 | 0.59 | 0.59 | 0.58 | 0.6 | 0.59 | 0.6 | 0.6 | 0.57 | 0.59 | 0.58 | 0.56 | 0.59 |
| 0.63 | 0.57 | 0.58 | 0.59 | 0.59 | 0.59 | 0.63 | 0.59 | 0.59 | 0.6 | 0.57 | 0.58 | 0.6 | 0.57 | 0.57 |
| 0.59 | 0.61 | 0.59 | 0.59 | 0.57 | 0.59 | 0.62 | 0.59 | 0.6 | 0.62 | 0.58 | 0.59 | 0.57 | 0.57 | 0.58 |
| 0.63 | 0.61 | 0.6 | 0.6 | 0.57 | 0.59 | 0.62 | 0.59 | 0.59 | 0.61 | 0.58 | 0.6 | 0.6 | 0.59 | 0.58 |
| 0.59 | 0.61 | 0.59 | 0.59 | 0.57 | 0.58 | 0.62 | 0.6 | 0.59 | 0.61 | 0.59 | 0.59 | 0.59 | 0.56 | 0.59 |
| 0.59 | 0.59 | 0.6 | 0.59 | 0.58 | 0.58 | 0.61 | 0.58 | 0.58 | 0.61 | 0.59 | 0.59 | 0.6 | 0.55 | 0.58 |

| 10HzE358 | 30HzE358 | 60HzE358 | 10Hz F359 | 30Hz F359 | 60Hz F359 | 10Hz S360 | 30Hz S360 | 60Hz S360 | 10HzN362 | 30HzN362 | 60HzN362 | 10Hz S367 | 30Hz S367 | 60Hz S367 |
| --- | --- | --- | --- | --- | --- | --- | --- | --- | --- | --- | --- | --- | --- | --- |
| 0.59 | 0.58 | 0.59 | 0.59 | 0.58 | 0.6 | 0.59 | 0.54 | 0.58 | 0.59 | 0.59 | 0.59 | 0.59 | 0.6 | 0.59 |
| 0.6 | 0.59 | 0.59 | 0.57 | 0.58 | 0.61 | 0.63 | 0.55 | 0.56 | 0.6 | 0.59 | 0.58 | 0.58 | 0.58 | 0.59 |
| 0.61 | 0.59 | 0.59 | 0.58 | 0.58 | 0.61 | 0.6 | 0.56 | 0.52 | 0.59 | 0.58 | 0.58 | 0.56 | 0.57 | 0.59 |
| 0.6 | 0.59 | 0.59 | 0.58 | 0.59 | 0.61 | 0.61 | 0.57 | 0.57 | 0.6 | 0.59 | 0.59 | 0.58 | 0.58 | 0.58 |
| 0.59 | 0.59 | 0.59 | 0.57 | 0.59 | 0.59 | 0.62 | 0.6 | 0.57 | 0.6 | 0.58 | 0.59 | 0.58 | 0.58 | 0.59 |
| 0.59 | 0.58 | 0.58 | 0.59 | 0.59 | 0.59 | 0.62 | 0.58 | 0.53 | 0.59 | 0.59 | 0.59 | 0.56 | 0.56 | 0.59 |
| 0.59 | 0.58 | 0.58 | 0.58 | 0.56 | 0.59 | 0.6 | 0.55 | 0.62 | 0.59 | 0.58 | 0.59 | 0.59 | 0.58 | 0.59 |
| 0.59 | 0.58 | 0.58 | 0.59 | 0.58 | 0.57 | 0.58 | 0.54 | 0.57 | 0.6 | 0.59 | 0.59 | 0.57 | 0.58 | 0.59 |
| 0.6 | 0.58 | 0.58 | 0.6 | 0.57 | 0.58 | 0.61 | 0.58 | 0.55 | 0.59 | 0.59 | 0.58 | 0.56 | 0.58 | 0.59 |
| 0.6 | 0.58 | 0.57 | 0.58 | 0.58 | 0.59 | 0.57 | 0.6 | 0.56 | 0.59 | 0.59 | 0.58 | 0.57 | 0.57 | 0.59 |
| 0.59 | 0.58 | 0.57 | 0.58 | 0.59 | 0.6 | 0.62 | 0.58 | 0.55 | 0.6 | 0.59 | 0.58 | 0.59 | 0.58 | 0.58 |
| 0.59 | 0.59 | 0.57 | 0.58 | 0.58 | 0.6 | 0.62 | 0.56 | 0.56 | 0.6 | 0.58 | 0.57 | 0.59 | 0.57 | 0.58 |
| 0.6 | 0.58 | 0.56 | 0.6 | 0.58 | 0.6 | 0.59 | 0.57 | 0.57 | 0.6 | 0.59 | 0.58 | 0.6 | 0.58 | 0.58 |
| 0.59 | 0.58 | 0.55 | 0.59 | 0.57 | 0.61 | 0.6 | 0.56 | 0.6 | 0.59 | 0.58 | 0.57 | 0.59 | 0.58 | 0.58 |
| 0.6 | 0.57 | 0.55 | 0.58 | 0.59 | 0.61 | 0.54 | 0.53 | 0.58 | 0.61 | 0.58 | 0.57 | 0.58 | 0.57 | 0.57 |
| 0.6 | 0.58 | 0.53 | 0.59 | 0.59 | 0.6 | 0.62 | 0.6 | 0.55 | 0.6 | 0.58 | 0.57 | 0.59 | 0.57 | 0.57 |
| 0.6 | 0.57 | 0.52 | 0.57 | 0.59 | 0.6 | 0.63 | 0.57 | 0.58 | 0.6 | 0.59 | 0.57 | 0.59 | 0.57 | 0.57 |
| 0.6 | 0.56 | 0.51 | 0.58 | 0.6 | 0.6 | 0.59 | 0.59 | 0.56 | 0.59 | 0.58 | 0.56 | 0.57 | 0.56 | 0.57 |
| 0.6 | 0.56 | 0.49 | 0.58 | 0.57 | 0.61 | 0.6 | 0.6 | 0.6 | 0.6 | 0.58 | 0.56 | 0.57 | 0.57 | 0.57 |
| 0.59 | 0.55 | 0.46 | 0.6 | 0.58 | 0.59 | 0.61 | 0.58 | 0.58 | 0.59 | 0.58 | 0.55 | 0.6 | 0.57 | 0.55 |
| 0.59 | 0.54 | 0.43 | 0.59 | 0.57 | 0.6 | 0.58 | 0.55 | 0.57 | 0.6 | 0.58 | 0.54 | 0.57 | 0.57 | 0.55 |
| 0.6 | 0.53 | 0.39 | 0.58 | 0.57 | 0.59 | 0.58 | 0.58 | 0.59 | 0.59 | 0.58 | 0.54 | 0.59 | 0.57 | 0.54 |
| 0.58 | 0.5 | 0.33 | 0.57 | 0.57 | 0.58 | 0.63 | 0.63 | 0.54 | 0.59 | 0.57 | 0.53 | 0.58 | 0.57 | 0.53 |
| 0.59 | 0.48 | 0.26 | 0.6 | 0.6 | 0.58 | 0.62 | 0.58 | 0.57 | 0.59 | 0.57 | 0.51 | 0.59 | 0.57 | 0.53 |
| 0.58 | 0.41 | 0.17 | 0.56 | 0.56 | 0.58 | 0.63 | 0.55 | 0.59 | 0.58 | 0.57 | 0.5 | 0.59 | 0.57 | 0.51 |
| 0.56 | 0.3 | 0.08 | 0.58 | 0.58 | 0.6 | 0.58 | 0.6 | 0.55 | 0.59 | 0.56 | 0.48 | 0.58 | 0.56 | 0.49 |
| 0.49 | 0.13 | 0.02 | 0.57 | 0.58 | 0.58 | 0.59 | 0.54 | 0.54 | 0.59 | 0.55 | 0.45 | 0.59 | 0.55 | 0.48 |
| 0.19 | 0.01 | 0 | 0.58 | 0.56 | 0.59 | 0.62 | 0.58 | 0.54 | 0.59 | 0.53 | 0.42 | 0.58 | 0.54 | 0.44 |
| 0.11 | 0 | 0 | 0.59 | 0.59 | 0.58 | 0.6 | 0.54 | 0.54 | 0.59 | 0.52 | 0.37 | 0.59 | 0.54 | 0.41 |
| 0.48 | 0.11 | 0.02 | 0.59 | 0.58 | 0.58 | 0.61 | 0.55 | 0.56 | 0.58 | 0.49 | 0.31 | 0.56 | 0.53 | 0.36 |
| 0.56 | 0.28 | 0.07 | 0.58 | 0.58 | 0.57 | 0.58 | 0.51 | 0.54 | 0.57 | 0.46 | 0.23 | 0.57 | 0.51 | 0.3 |
| 0.57 | 0.39 | 0.15 | 0.57 | 0.58 | 0.58 | 0.57 | 0.56 | 0.51 | 0.57 | 0.39 | 0.14 | 0.56 | 0.48 | 0.21 |
| 0.58 | 0.46 | 0.24 | 0.58 | 0.57 | 0.57 | 0.61 | 0.51 | 0.45 | 0.53 | 0.25 | 0.05 | 0.56 | 0.42 | 0.12 |
| 0.59 | 0.5 | 0.31 | 0.6 | 0.56 | 0.56 | 0.62 | 0.54 | 0.44 | 0.41 | 0.07 | 0.01 | 0.56 | 0.34 | 0.05 |
| 0.59 | 0.52 | 0.37 | 0.57 | 0.55 | 0.53 | 0.65 | 0.55 | 0.46 | 0.05 | 0.01 | 0 | 0.49 | 0.19 | 0.01 |
| 0.59 | 0.53 | 0.41 | 0.59 | 0.56 | 0.53 | 0.61 | 0.53 | 0.48 | 0.37 | 0.05 | 0 | 0.3 | 0.02 | 0.01 |
| 0.59 | 0.54 | 0.44 | 0.57 | 0.55 | 0.52 | 0.63 | 0.55 | 0.45 | 0.53 | 0.23 | 0.05 | 0.07 | 0.01 | 0.01 |
| 0.59 | 0.54 | 0.46 | 0.58 | 0.54 | 0.51 | 0.59 | 0.55 | 0.42 | 0.58 | 0.37 | 0.13 | 0.42 | 0.06 | 0.04 |
| 0.59 | 0.54 | 0.48 | 0.57 | 0.52 | 0.49 | 0.57 | 0.55 | 0.4 | 0.57 | 0.46 | 0.22 | 0.52 | 0.22 | 0.1 |
| 0.6 | 0.56 | 0.5 | 0.56 | 0.52 | 0.47 | 0.62 | 0.51 | 0.37 | 0.58 | 0.49 | 0.3 | 0.55 | 0.35 | 0.18 |
| 0.59 | 0.57 | 0.52 | 0.58 | 0.53 | 0.46 | 0.64 | 0.45 | 0.3 | 0.58 | 0.52 | 0.37 | 0.55 | 0.4 | 0.27 |
| 0.59 | 0.57 | 0.53 | 0.58 | 0.54 | 0.43 | 0.51 | 0.46 | 0.29 | 0.6 | 0.54 | 0.42 | 0.57 | 0.48 | 0.34 |
| 0.6 | 0.57 | 0.55 | 0.58 | 0.53 | 0.4 | 0.55 | 0.45 | 0.18 | 0.6 | 0.55 | 0.45 | 0.58 | 0.5 | 0.39 |
| 0.6 | 0.57 | 0.56 | 0.58 | 0.51 | 0.37 | 0.5 | 0.28 | 0.09 | 0.6 | 0.56 | 0.48 | 0.58 | 0.53 | 0.44 |
| 0.6 | 0.58 | 0.56 | 0.57 | 0.49 | 0.33 | 0.36 | 0.1 | 0.01 | 0.58 | 0.56 | 0.49 | 0.57 | 0.54 | 0.46 |
| 0.6 | 0.58 | 0.57 | 0.56 | 0.45 | 0.28 | 0.21 | 0 | -0.03 | 0.6 | 0.58 | 0.51 | 0.58 | 0.54 | 0.49 |
| 0.59 | 0.58 | 0.57 | 0.59 | 0.42 | 0.17 | 0.04 | 0.03 | 0.04 | 0.59 | 0.58 | 0.53 | 0.58 | 0.54 | 0.5 |
| 0.6 | 0.58 | 0.57 | 0.56 | 0.33 | 0.09 | 0.4 | 0 | 0.03 | 0.59 | 0.58 | 0.53 | 0.58 | 0.55 | 0.52 |
| 0.6 | 0.58 | 0.58 | 0.49 | 0.19 | 0.03 | 0.45 | 0.2 | 0.03 | 0.59 | 0.58 | 0.54 | 0.59 | 0.57 | 0.53 |
| 0.59 | 0.59 | 0.58 | 0.34 | 0.02 | 0.01 | 0.58 | 0.3 | 0.14 | 0.6 | 0.58 | 0.55 | 0.59 | 0.56 | 0.54 |
| 0.59 | 0.58 | 0.58 | 0.05 | 0.01 | 0 | 0.57 | 0.46 | 0.23 | 0.59 | 0.58 | 0.56 | 0.59 | 0.57 | 0.55 |
| 0.6 | 0.58 | 0.59 | 0.37 | 0.04 | 0.01 | 0.57 | 0.47 | 0.23 | 0.59 | 0.58 | 0.57 | 0.59 | 0.57 | 0.56 |
| 0.6 | 0.58 | 0.59 | 0.51 | 0.2 | 0.04 | 0.68 | 0.53 | 0.35 | 0.59 | 0.58 | 0.56 | 0.58 | 0.56 | 0.56 |
| 0.6 | 0.58 | 0.59 | 0.56 | 0.34 | 0.1 | 0.58 | 0.53 | 0.38 | 0.59 | 0.58 | 0.57 | 0.58 | 0.57 | 0.56 |
| 0.6 | 0.59 | 0.59 | 0.56 | 0.43 | 0.2 | 0.62 | 0.57 | 0.44 | 0.6 | 0.58 | 0.57 | 0.59 | 0.56 | 0.57 |
| 0.6 | 0.59 | 0.59 | 0.58 | 0.46 | 0.27 | 0.64 | 0.57 | 0.48 | 0.61 | 0.59 | 0.57 | 0.58 | 0.58 | 0.58 |
| 0.59 | 0.59 | 0.59 | 0.58 | 0.51 | 0.35 | 0.56 | 0.6 | 0.44 | 0.59 | 0.58 | 0.57 | 0.59 | 0.58 | 0.57 |
| 0.59 | 0.59 | 0.6 | 0.57 | 0.52 | 0.41 | 0.6 | 0.53 | 0.48 | 0.59 | 0.58 | 0.57 | 0.58 | 0.58 | 0.58 |
| 0.6 | 0.58 | 0.6 | 0.58 | 0.52 | 0.45 | 0.57 | 0.54 | 0.52 | 0.59 | 0.59 | 0.58 | 0.58 | 0.57 | 0.58 |
| 0.59 | 0.59 | 0.6 | 0.6 | 0.55 | 0.47 | 0.6 | 0.55 | 0.52 | 0.6 | 0.58 | 0.58 | 0.59 | 0.58 | 0.58 |
| 0.6 | 0.58 | 0.59 | 0.56 | 0.54 | 0.49 | 0.53 | 0.54 | 0.53 | 0.61 | 0.59 | 0.58 | 0.59 | 0.58 | 0.6 |
| 0.59 | 0.59 | 0.6 | 0.59 | 0.54 | 0.51 | 0.62 | 0.57 | 0.53 | 0.58 | 0.59 | 0.58 | 0.59 | 0.58 | 0.58 |
| 0.59 | 0.59 | 0.6 | 0.59 | 0.57 | 0.52 | 0.6 | 0.56 | 0.55 | 0.61 | 0.58 | 0.58 | 0.58 | 0.58 | 0.59 |

| 10Hz F368 | 30Hz F368 | 60Hz F368 | 10Hz A369 | 30Hz A369 | 60Hz A369 | 10Hz T370 | 30Hz T370 | 60Hz T370 | 10Hz A373 | 30Hz A373 | 60Hz A373 |
| --- | --- | --- | --- | --- | --- | --- | --- | --- | --- | --- | --- |
| 0.59 | 0.6 | 0.59 | 0.58 | 0.58 | 0.53 | 0.54 | 0.54 | 0.57 | 0.59 | 0.59 | 0.59 |
| 0.58 | 0.58 | 0.59 | 0.63 | 0.54 | 0.54 | 0.57 | 0.55 | 0.57 | 0.59 | 0.58 | 0.58 |
| 0.56 | 0.57 | 0.59 | 0.55 | 0.58 | 0.55 | 0.56 | 0.54 | 0.55 | 0.59 | 0.57 | 0.56 |
| 0.58 | 0.58 | 0.59 | 0.59 | 0.6 | 0.56 | 0.57 | 0.55 | 0.56 | 0.59 | 0.57 | 0.57 |
| 0.58 | 0.58 | 0.6 | 0.55 | 0.59 | 0.58 | 0.57 | 0.56 | 0.55 | 0.6 | 0.57 | 0.58 |
| 0.56 | 0.56 | 0.59 | 0.58 | 0.59 | 0.55 | 0.58 | 0.55 | 0.57 | 0.58 | 0.58 | 0.59 |
| 0.59 | 0.58 | 0.61 | 0.6 | 0.6 | 0.58 | 0.57 | 0.56 | 0.56 | 0.59 | 0.58 | 0.59 |
| 0.57 | 0.58 | 0.6 | 0.59 | 0.58 | 0.55 | 0.56 | 0.54 | 0.57 | 0.59 | 0.58 | 0.59 |
| 0.56 | 0.58 | 0.58 | 0.58 | 0.57 | 0.56 | 0.58 | 0.55 | 0.55 | 0.58 | 0.58 | 0.59 |
| 0.57 | 0.57 | 0.59 | 0.6 | 0.57 | 0.56 | 0.57 | 0.55 | 0.54 | 0.6 | 0.56 | 0.58 |
| 0.59 | 0.58 | 0.58 | 0.61 | 0.6 | 0.54 | 0.57 | 0.55 | 0.56 | 0.61 | 0.57 | 0.58 |
| 0.59 | 0.57 | 0.58 | 0.59 | 0.57 | 0.55 | 0.55 | 0.55 | 0.54 | 0.59 | 0.58 | 0.58 |
| 0.6 | 0.58 | 0.59 | 0.6 | 0.59 | 0.59 | 0.57 | 0.54 | 0.52 | 0.6 | 0.58 | 0.58 |
| 0.59 | 0.58 | 0.58 | 0.6 | 0.63 | 0.56 | 0.54 | 0.55 | 0.53 | 0.59 | 0.57 | 0.59 |
| 0.58 | 0.57 | 0.57 | 0.58 | 0.58 | 0.56 | 0.58 | 0.53 | 0.51 | 0.58 | 0.57 | 0.58 |
| 0.59 | 0.57 | 0.58 | 0.58 | 0.57 | 0.56 | 0.55 | 0.53 | 0.5 | 0.58 | 0.57 | 0.58 |
| 0.59 | 0.57 | 0.57 | 0.56 | 0.59 | 0.57 | 0.57 | 0.54 | 0.49 | 0.6 | 0.57 | 0.57 |
| 0.57 | 0.56 | 0.57 | 0.57 | 0.57 | 0.56 | 0.58 | 0.52 | 0.46 | 0.59 | 0.58 | 0.58 |
| 0.57 | 0.57 | 0.57 | 0.59 | 0.62 | 0.54 | 0.57 | 0.51 | 0.43 | 0.59 | 0.57 | 0.58 |
| 0.6 | 0.57 | 0.56 | 0.59 | 0.58 | 0.55 | 0.58 | 0.51 | 0.4 | 0.58 | 0.59 | 0.57 |
| 0.57 | 0.57 | 0.56 | 0.59 | 0.61 | 0.54 | 0.57 | 0.46 | 0.35 | 0.59 | 0.58 | 0.57 |
| 0.59 | 0.57 | 0.55 | 0.58 | 0.57 | 0.54 | 0.54 | 0.44 | 0.28 | 0.6 | 0.57 | 0.58 |
| 0.58 | 0.57 | 0.55 | 0.57 | 0.57 | 0.56 | 0.56 | 0.38 | 0.17 | 0.59 | 0.56 | 0.57 |
| 0.59 | 0.57 | 0.53 | 0.57 | 0.57 | 0.55 | 0.51 | 0.27 | 0.08 | 0.6 | 0.58 | 0.58 |
| 0.59 | 0.57 | 0.52 | 0.61 | 0.6 | 0.54 | 0.38 | 0.13 | 0.02 | 0.57 | 0.58 | 0.57 |
| 0.58 | 0.56 | 0.5 | 0.59 | 0.59 | 0.55 | 0.17 | 0.02 | 0 | 0.58 | 0.57 | 0.57 |
| 0.59 | 0.55 | 0.49 | 0.63 | 0.58 | 0.53 | 0.1 | -0.01 | 0 | 0.59 | 0.58 | 0.56 |
| 0.58 | 0.54 | 0.46 | 0.6 | 0.61 | 0.52 | 0.48 | 0.12 | 0.02 | 0.59 | 0.57 | 0.56 |
| 0.59 | 0.54 | 0.44 | 0.57 | 0.59 | 0.54 | 0.5 | 0.3 | 0.09 | 0.59 | 0.57 | 0.56 |
| 0.56 | 0.53 | 0.39 | 0.61 | 0.55 | 0.52 | 0.54 | 0.38 | 0.18 | 0.59 | 0.57 | 0.55 |
| 0.57 | 0.51 | 0.34 | 0.61 | 0.59 | 0.53 | 0.54 | 0.44 | 0.27 | 0.57 | 0.57 | 0.55 |
| 0.56 | 0.48 | 0.27 | 0.63 | 0.56 | 0.52 | 0.57 | 0.49 | 0.33 | 0.6 | 0.55 | 0.54 |
| 0.56 | 0.42 | 0.18 | 0.6 | 0.56 | 0.5 | 0.56 | 0.51 | 0.38 | 0.57 | 0.56 | 0.52 |
| 0.56 | 0.34 | 0.09 | 0.57 | 0.57 | 0.47 | 0.55 | 0.51 | 0.43 | 0.58 | 0.56 | 0.51 |
| 0.49 | 0.19 | 0.03 | 0.59 | 0.58 | 0.48 | 0.56 | 0.5 | 0.48 | 0.59 | 0.56 | 0.5 |
| 0.3 | 0.02 | 0.01 | 0.63 | 0.59 | 0.44 | 0.57 | 0.53 | 0.49 | 0.57 | 0.56 | 0.48 |
| 0.07 | 0.01 | 0.01 | 0.59 | 0.54 | 0.42 | 0.58 | 0.51 | 0.5 | 0.58 | 0.55 | 0.47 |
| 0.42 | 0.06 | 0.01 | 0.55 | 0.54 | 0.42 | 0.58 | 0.52 | 0.52 | 0.59 | 0.54 | 0.45 |
| 0.52 | 0.22 | 0.06 | 0.55 | 0.55 | 0.35 | 0.57 | 0.55 | 0.5 | 0.59 | 0.53 | 0.42 |
| 0.55 | 0.35 | 0.13 | 0.58 | 0.49 | 0.31 | 0.56 | 0.55 | 0.54 | 0.58 | 0.51 | 0.37 |
| 0.55 | 0.4 | 0.22 | 0.59 | 0.47 | 0.23 | 0.55 | 0.54 | 0.53 | 0.57 | 0.5 | 0.31 |
| 0.57 | 0.48 | 0.29 | 0.6 | 0.42 | 0.14 | 0.56 | 0.56 | 0.55 | 0.56 | 0.44 | 0.23 |
| 0.58 | 0.5 | 0.36 | 0.54 | 0.32 | 0.06 | 0.58 | 0.55 | 0.56 | 0.55 | 0.37 | 0.14 |
| 0.58 | 0.53 | 0.42 | 0.47 | 0.13 | 0.02 | 0.58 | 0.55 | 0.56 | 0.53 | 0.26 | 0.06 |
| 0.57 | 0.54 | 0.44 | 0.08 | 0.02 | -0.01 | 0.57 | 0.54 | 0.54 | 0.44 | 0.09 | 0.01 |
| 0.58 | 0.54 | 0.48 | 0.2 | 0.02 | 0.01 | 0.56 | 0.55 | 0.56 | 0.05 | 0.01 | 0.01 |
| 0.58 | 0.54 | 0.5 | 0.51 | 0.14 | 0.02 | 0.55 | 0.56 | 0.58 | 0.27 | 0.02 | 0 |
| 0.58 | 0.55 | 0.52 | 0.56 | 0.33 | 0.09 | 0.57 | 0.56 | 0.56 | 0.5 | 0.17 | 0.02 |
| 0.59 | 0.57 | 0.53 | 0.56 | 0.43 | 0.16 | 0.61 | 0.56 | 0.56 | 0.55 | 0.33 | 0.09 |
| 0.59 | 0.56 | 0.54 | 0.56 | 0.46 | 0.25 | 0.58 | 0.54 | 0.56 | 0.56 | 0.43 | 0.18 |
| 0.59 | 0.57 | 0.55 | 0.57 | 0.51 | 0.31 | 0.58 | 0.55 | 0.57 | 0.56 | 0.47 | 0.27 |
| 0.59 | 0.57 | 0.55 | 0.59 | 0.55 | 0.36 | 0.57 | 0.54 | 0.56 | 0.58 | 0.51 | 0.33 |
| 0.58 | 0.56 | 0.56 | 0.57 | 0.53 | 0.42 | 0.56 | 0.55 | 0.57 | 0.59 | 0.53 | 0.39 |
| 0.58 | 0.57 | 0.56 | 0.61 | 0.54 | 0.44 | 0.57 | 0.54 | 0.58 | 0.59 | 0.54 | 0.43 |
| 0.59 | 0.56 | 0.57 | 0.6 | 0.56 | 0.46 | 0.59 | 0.54 | 0.57 | 0.58 | 0.54 | 0.47 |
| 0.58 | 0.58 | 0.58 | 0.56 | 0.56 | 0.47 | 0.55 | 0.54 | 0.57 | 0.59 | 0.55 | 0.48 |
| 0.59 | 0.58 | 0.57 | 0.59 | 0.59 | 0.49 | 0.57 | 0.54 | 0.57 | 0.61 | 0.56 | 0.49 |
| 0.58 | 0.58 | 0.57 | 0.6 | 0.56 | 0.52 | 0.59 | 0.55 | 0.57 | 0.59 | 0.55 | 0.52 |
| 0.58 | 0.57 | 0.58 | 0.61 | 0.57 | 0.5 | 0.56 | 0.55 | 0.58 | 0.59 | 0.55 | 0.52 |
| 0.59 | 0.58 | 0.58 | 0.6 | 0.58 | 0.5 | 0.58 | 0.54 | 0.58 | 0.59 | 0.56 | 0.53 |
| 0.59 | 0.58 | 0.58 | 0.57 | 0.62 | 0.53 | 0.56 | 0.55 | 0.56 | 0.59 | 0.57 | 0.53 |
| 0.59 | 0.58 | 0.58 | 0.58 | 0.6 | 0.54 | 0.58 | 0.56 | 0.58 | 0.61 | 0.57 | 0.55 |
| 0.58 | 0.58 | 0.59 | 0.6 | 0.6 | 0.52 | 0.56 | 0.56 | 0.58 | 0.59 | 0.56 | 0.54 |

**Table S2.** ^15^N CEST I/I_0_ data measured for FUS-RRM at pH 6.4.

|  | **I/I_0_** | | | | | | | | | | | | | | | |
| --- | --- | --- | --- | --- | --- | --- | --- | --- | --- | --- | --- | --- | --- | --- | --- | --- |
| **^15^N Offset (ppm)** | **N285** | **G293** | **E294** | **T297** | **E299** | **S300** | **D303** | **F305** | **Q307** | **T317** | **T330** | **T338** | **D352** | **D355** | **I351** | **G356** |
| 102.0 | 0.58 | 0.51 | 0.60 | 0.58 | 0.58 | 0.59 | 0.58 | 0.59 | 0.56 | 0.54 | 0.58 | 0.54 | 0.58 | 0.58 | 0.61 | 0.57 |
| 102.5 | 0.58 | 0.47 | 0.60 | 0.59 | 0.58 | 0.59 | 0.58 | 0.58 | 0.57 | 0.53 | 0.57 | 0.54 | 0.58 | 0.59 | 0.60 | 0.57 |
| 103.0 | 0.57 | 0.42 | 0.59 | 0.59 | 0.58 | 0.59 | 0.57 | 0.58 | 0.57 | 0.50 | 0.55 | 0.56 | 0.57 | 0.59 | 0.60 | 0.56 |
| 103.5 | 0.56 | 0.35 | 0.61 | 0.59 | 0.58 | 0.60 | 0.58 | 0.58 | 0.57 | 0.46 | 0.53 | 0.55 | 0.58 | 0.59 | 0.60 | 0.57 |
| 104.0 | 0.58 | 0.19 | 0.60 | 0.58 | 0.58 | 0.59 | 0.58 | 0.58 | 0.56 | 0.37 | 0.51 | 0.53 | 0.56 | 0.59 | 0.61 | 0.56 |
| 104.5 | 0.56 | 0.03 | 0.60 | 0.58 | 0.58 | 0.59 | 0.58 | 0.57 | 0.57 | 0.20 | 0.47 | 0.52 | 0.57 | 0.59 | 0.60 | 0.56 |
| 105.0 | 0.57 | 0.01 | 0.59 | 0.58 | 0.58 | 0.58 | 0.58 | 0.56 | 0.57 | 0.04 | 0.40 | 0.51 | 0.58 | 0.59 | 0.58 | 0.56 |
| 105.5 | 0.57 | 0.07 | 0.59 | 0.58 | 0.58 | 0.59 | 0.58 | 0.57 | 0.56 | 0.00 | 0.28 | 0.50 | 0.59 | 0.58 | 0.61 | 0.57 |
| 106.0 | 0.58 | 0.23 | 0.60 | 0.58 | 0.58 | 0.59 | 0.58 | 0.58 | 0.56 | 0.06 | 0.12 | 0.47 | 0.58 | 0.58 | 0.59 | 0.57 |
| 106.5 | 0.57 | 0.37 | 0.59 | 0.57 | 0.58 | 0.59 | 0.58 | 0.57 | 0.56 | 0.25 | 0.00 | 0.41 | 0.57 | 0.59 | 0.62 | 0.57 |
| 107.0 | 0.58 | 0.43 | 0.59 | 0.56 | 0.57 | 0.59 | 0.58 | 0.57 | 0.56 | 0.40 | 0.01 | 0.30 | 0.58 | 0.58 | 0.61 | 0.56 |
| 107.5 | 0.56 | 0.49 | 0.60 | 0.56 | 0.57 | 0.59 | 0.57 | 0.57 | 0.56 | 0.47 | 0.04 | 0.15 | 0.58 | 0.59 | 0.60 | 0.54 |
| 108.0 | 0.58 | 0.52 | 0.59 | 0.55 | 0.59 | 0.58 | 0.58 | 0.55 | 0.56 | 0.51 | 0.21 | 0.02 | 0.57 | 0.59 | 0.59 | 0.55 |
| 108.5 | 0.57 | 0.51 | 0.57 | 0.52 | 0.57 | 0.58 | 0.57 | 0.54 | 0.56 | 0.53 | 0.36 | 0.00 | 0.58 | 0.59 | 0.59 | 0.55 |
| 109.0 | 0.57 | 0.50 | 0.56 | 0.50 | 0.58 | 0.57 | 0.57 | 0.54 | 0.56 | 0.55 | 0.43 | 0.06 | 0.57 | 0.58 | 0.59 | 0.51 |
| 109.5 | 0.56 | 0.53 | 0.56 | 0.46 | 0.58 | 0.58 | 0.57 | 0.52 | 0.55 | 0.55 | 0.48 | 0.23 | 0.58 | 0.57 | 0.61 | 0.51 |
| 110.0 | 0.56 | 0.54 | 0.55 | 0.38 | 0.57 | 0.57 | 0.57 | 0.48 | 0.55 | 0.57 | 0.51 | 0.36 | 0.57 | 0.59 | 0.59 | 0.52 |
| 110.5 | 0.56 | 0.57 | 0.54 | 0.24 | 0.57 | 0.56 | 0.56 | 0.45 | 0.55 | 0.57 | 0.53 | 0.43 | 0.56 | 0.57 | 0.57 | 0.55 |
| 111.0 | 0.55 | 0.58 | 0.51 | 0.06 | 0.57 | 0.56 | 0.55 | 0.37 | 0.55 | 0.58 | 0.56 | 0.46 | 0.56 | 0.57 | 0.60 | 0.53 |
| 111.5 | 0.57 | 0.57 | 0.47 | 0.00 | 0.57 | 0.55 | 0.54 | 0.22 | 0.55 | 0.57 | 0.56 | 0.49 | 0.58 | 0.57 | 0.58 | 0.53 |
| 112.0 | 0.56 | 0.58 | 0.39 | 0.04 | 0.57 | 0.55 | 0.53 | 0.05 | 0.53 | 0.56 | 0.56 | 0.51 | 0.57 | 0.58 | 0.58 | 0.50 |
| 112.5 | 0.55 | 0.56 | 0.26 | 0.22 | 0.56 | 0.53 | 0.51 | 0.01 | 0.52 | 0.56 | 0.55 | 0.52 | 0.55 | 0.57 | 0.59 | 0.49 |
| 113.0 | 0.55 | 0.57 | 0.07 | 0.36 | 0.56 | 0.51 | 0.48 | 0.03 | 0.51 | 0.56 | 0.54 | 0.53 | 0.55 | 0.56 | 0.58 | 0.45 |
| 113.5 | 0.57 | 0.57 | 0.00 | 0.45 | 0.56 | 0.48 | 0.43 | 0.20 | 0.47 | 0.56 | 0.55 | 0.52 | 0.53 | 0.56 | 0.60 | 0.40 |
| 114.0 | 0.56 | 0.57 | 0.01 | 0.50 | 0.55 | 0.41 | 0.34 | 0.35 | 0.44 | 0.58 | 0.58 | 0.51 | 0.53 | 0.55 | 0.60 | 0.32 |
| 114.5 | 0.56 | 0.57 | 0.14 | 0.52 | 0.52 | 0.30 | 0.19 | 0.44 | 0.38 | 0.60 | 0.59 | 0.52 | 0.52 | 0.54 | 0.58 | 0.16 |
| 115.0 | 0.54 | 0.58 | 0.31 | 0.55 | 0.51 | 0.13 | 0.03 | 0.48 | 0.27 | 0.59 | 0.59 | 0.54 | 0.50 | 0.53 | 0.56 | 0.01 |
| 115.5 | 0.55 | 0.58 | 0.42 | 0.55 | 0.48 | 0.01 | 0.00 | 0.50 | 0.09 | 0.60 | 0.60 | 0.56 | 0.45 | 0.50 | 0.55 | 0.00 |
| 116.0 | 0.53 | 0.58 | 0.48 | 0.56 | 0.44 | 0.01 | 0.05 | 0.52 | -0.01 | 0.60 | 0.60 | 0.56 | 0.41 | 0.46 | 0.52 | 0.04 |
| 116.5 | 0.53 | 0.58 | 0.51 | 0.55 | 0.38 | 0.11 | 0.22 | 0.53 | 0.01 | 0.59 | 0.60 | 0.55 | 0.31 | 0.41 | 0.56 | 0.19 |
| 117.0 | 0.51 | 0.56 | 0.53 | 0.55 | 0.24 | 0.28 | 0.35 | 0.53 | 0.14 | 0.59 | 0.60 | 0.57 | 0.14 | 0.32 | 0.51 | 0.32 |
| 117.5 | 0.49 | 0.58 | 0.54 | 0.54 | 0.07 | 0.39 | 0.42 | 0.53 | 0.29 | 0.60 | 0.61 | 0.56 | 0.02 | 0.15 | 0.45 | 0.40 |
| 118.0 | 0.45 | 0.57 | 0.56 | 0.55 | 0.01 | 0.46 | 0.45 | 0.54 | 0.39 | 0.60 | 0.61 | 0.55 | 0.00 | 0.02 | 0.39 | 0.48 |
| 118.5 | 0.37 | 0.57 | 0.56 | 0.57 | 0.01 | 0.50 | 0.49 | 0.56 | 0.44 | 0.61 | 0.60 | 0.55 | 0.08 | 0.01 | 0.26 | 0.49 |
| 119.0 | 0.25 | 0.58 | 0.56 | 0.58 | 0.17 | 0.53 | 0.53 | 0.56 | 0.45 | 0.62 | 0.60 | 0.56 | 0.25 | 0.06 | 0.08 | 0.51 |
| 119.5 | 0.08 | 0.57 | 0.55 | 0.59 | 0.32 | 0.54 | 0.54 | 0.57 | 0.48 | 0.61 | 0.60 | 0.56 | 0.37 | 0.23 | 0.02 | 0.53 |
| 120.0 | 0.00 | 0.57 | 0.56 | 0.57 | 0.42 | 0.54 | 0.55 | 0.57 | 0.51 | 0.60 | 0.61 | 0.57 | 0.44 | 0.35 | 0.01 | 0.53 |
| 120.5 | 0.02 | 0.58 | 0.58 | 0.59 | 0.46 | 0.54 | 0.56 | 0.58 | 0.53 | 0.60 | 0.62 | 0.56 | 0.47 | 0.43 | 0.16 | 0.55 |
| 121.0 | 0.15 | 0.58 | 0.59 | 0.59 | 0.50 | 0.55 | 0.55 | 0.56 | 0.54 | 0.60 | 0.62 | 0.57 | 0.50 | 0.47 | 0.29 | 0.53 |
| 121.5 | 0.31 | 0.57 | 0.60 | 0.59 | 0.51 | 0.56 | 0.57 | 0.57 | 0.54 | 0.60 | 0.62 | 0.56 | 0.52 | 0.48 | 0.40 | 0.54 |
| 122.0 | 0.42 | 0.58 | 0.60 | 0.59 | 0.53 | 0.58 | 0.57 | 0.58 | 0.55 | 0.60 | 0.61 | 0.58 | 0.54 | 0.50 | 0.47 | 0.54 |
| 122.5 | 0.47 | 0.58 | 0.59 | 0.58 | 0.52 | 0.58 | 0.58 | 0.58 | 0.55 | 0.61 | 0.61 | 0.57 | 0.53 | 0.54 | 0.52 | 0.56 |
| 123.0 | 0.49 | 0.57 | 0.60 | 0.59 | 0.52 | 0.58 | 0.58 | 0.58 | 0.55 | 0.60 | 0.61 | 0.57 | 0.52 | 0.55 | 0.53 | 0.56 |
| 123.5 | 0.52 | 0.58 | 0.60 | 0.58 | 0.52 | 0.58 | 0.58 | 0.56 | 0.55 | 0.61 | 0.60 | 0.58 | 0.54 | 0.56 | 0.54 | 0.56 |
| 124.0 | 0.54 | 0.59 | 0.60 | 0.59 | 0.55 | 0.58 | 0.58 | 0.59 | 0.56 | 0.61 | 0.61 | 0.57 | 0.55 | 0.57 | 0.57 | 0.56 |
| 124.5 | 0.54 | 0.59 | 0.61 | 0.59 | 0.56 | 0.58 | 0.58 | 0.58 | 0.56 | 0.60 | 0.60 | 0.58 | 0.56 | 0.57 | 0.56 | 0.56 |
| 125.0 | 0.55 | 0.58 | 0.60 | 0.60 | 0.56 | 0.58 | 0.58 | 0.59 | 0.56 | 0.60 | 0.61 | 0.57 | 0.56 | 0.57 | 0.60 | 0.58 |
| 125.5 | 0.57 | 0.59 | 0.60 | 0.60 | 0.57 | 0.59 | 0.58 | 0.58 | 0.56 | 0.61 | 0.61 | 0.57 | 0.57 | 0.58 | 0.60 | 0.56 |
| 126.0 | 0.56 | 0.58 | 0.60 | 0.60 | 0.57 | 0.59 | 0.59 | 0.58 | 0.55 | 0.60 | 0.61 | 0.56 | 0.58 | 0.58 | 0.59 | 0.55 |
| 126.5 | 0.56 | 0.57 | 0.60 | 0.60 | 0.57 | 0.59 | 0.58 | 0.59 | 0.56 | 0.60 | 0.62 | 0.57 | 0.58 | 0.58 | 0.59 | 0.58 |
| 127.0 | 0.56 | 0.58 | 0.60 | 0.59 | 0.57 | 0.58 | 0.59 | 0.58 | 0.56 | 0.60 | 0.61 | 0.57 | 0.58 | 0.58 | 0.60 | 0.59 |
| 127.5 | 0.55 | 0.58 | 0.61 | 0.59 | 0.58 | 0.59 | 0.59 | 0.58 | 0.56 | 0.60 | 0.60 | 0.56 | 0.59 | 0.57 | 0.61 | 0.57 |
| 128.0 | 0.57 | 0.58 | 0.60 | 0.60 | 0.57 | 0.58 | 0.59 | 0.59 | 0.56 | 0.60 | 0.61 | 0.57 | 0.58 | 0.57 | 0.59 | 0.56 |
| 128.5 | 0.56 | 0.59 | 0.61 | 0.60 | 0.58 | 0.59 | 0.58 | 0.59 | 0.56 | 0.60 | 0.60 | 0.57 | 0.59 | 0.57 | 0.61 | 0.57 |
| 129.0 | 0.58 | 0.59 | 0.60 | 0.60 | 0.57 | 0.59 | 0.58 | 0.58 | 0.57 | 0.61 | 0.62 | 0.56 | 0.58 | 0.59 | 0.60 | 0.56 |
| 129.5 | 0.57 | 0.58 | 0.60 | 0.60 | 0.58 | 0.59 | 0.59 | 0.60 | 0.57 | 0.60 | 0.61 | 0.56 | 0.58 | 0.58 | 0.59 | 0.57 |
| 130.0 | 0.56 | 0.58 | 0.61 | 0.59 | 0.56 | 0.59 | 0.58 | 0.58 | 0.56 | 0.61 | 0.61 | 0.57 | 0.59 | 0.58 | 0.59 | 0.56 |
| 130.5 | 0.56 | 0.58 | 0.60 | 0.59 | 0.58 | 0.59 | 0.59 | 0.58 | 0.56 | 0.60 | 0.62 | 0.57 | 0.59 | 0.57 | 0.62 | 0.56 |
| 131.0 | 0.56 | 0.57 | 0.60 | 0.60 | 0.58 | 0.59 | 0.58 | 0.58 | 0.56 | 0.61 | 0.62 | 0.57 | 0.59 | 0.57 | 0.57 | 0.57 |
| 131.5 | 0.58 | 0.59 | 0.60 | 0.60 | 0.58 | 0.60 | 0.59 | 0.58 | 0.57 | 0.61 | 0.63 | 0.57 | 0.57 | 0.58 | 0.61 | 0.57 |
| 132.0 | 0.58 | 0.58 | 0.60 | 0.60 | 0.58 | 0.59 | 0.58 | 0.60 | 0.57 | 0.61 | 0.61 | 0.57 | 0.57 | 0.58 | 0.61 | 0.59 |
| 132.5 | 0.58 | 0.58 | 0.60 | 0.59 | 0.58 | 0.59 | 0.58 | 0.58 | 0.57 | 0.61 | 0.60 | 0.55 | 0.57 | 0.59 | 0.61 | 0.56 |
| 133.0 | 0.57 | 0.58 | 0.61 | 0.59 | 0.58 | 0.59 | 0.59 | 0.59 | 0.56 | 0.61 | 0.60 | 0.57 | 0.58 | 0.59 | 0.59 | 0.55 |

**Table S3.** ^15^N CEST I/I_0_ data measured for FUS-RRM at pH 4.6.

|  | **I/I_0_** | | | | | | | | | |
| --- | --- | --- | --- | --- | --- | --- | --- | --- | --- | --- |
| **^15^N offset (ppm)** | **E294** | **T297** | **E299** | **F305** | **Q307** | **T317** | **T330** | **T338** | **D352** | **S360** |
| 102.0 | 0.56 | 0.58 | 0.59 | 0.58 | 0.58 | 0.56 | 0.52 | 0.55 | 0.55 | 0.57 |
| 102.5 | 0.59 | 0.56 | 0.57 | 0.60 | 0.55 | 0.55 | 0.60 | 0.56 | 0.58 | 0.54 |
| 103.0 | 0.60 | 0.60 | 0.59 | 0.60 | 0.58 | 0.50 | 0.54 | 0.54 | 0.60 | 0.59 |
| 103.5 | 0.60 | 0.61 | 0.58 | 0.56 | 0.56 | 0.47 | 0.53 | 0.53 | 0.59 | 0.62 |
| 104.0 | 0.60 | 0.59 | 0.58 | 0.61 | 0.54 | 0.38 | 0.48 | 0.55 | 0.58 | 0.65 |
| 104.5 | 0.59 | 0.58 | 0.60 | 0.58 | 0.55 | 0.21 | 0.46 | 0.51 | 0.57 | 0.57 |
| 105.0 | 0.58 | 0.59 | 0.59 | 0.58 | 0.55 | 0.06 | 0.37 | 0.51 | 0.58 | 0.59 |
| 105.5 | 0.62 | 0.60 | 0.59 | 0.58 | 0.56 | -0.02 | 0.29 | 0.51 | 0.58 | 0.62 |
| 106.0 | 0.58 | 0.54 | 0.58 | 0.62 | 0.55 | 0.04 | 0.14 | 0.48 | 0.58 | 0.62 |
| 106.5 | 0.61 | 0.55 | 0.59 | 0.57 | 0.57 | 0.20 | 0.03 | 0.39 | 0.58 | 0.59 |
| 107.0 | 0.57 | 0.57 | 0.58 | 0.57 | 0.55 | 0.37 | 0.00 | 0.31 | 0.60 | 0.62 |
| 107.5 | 0.58 | 0.56 | 0.58 | 0.55 | 0.56 | 0.45 | 0.04 | 0.15 | 0.58 | 0.57 |
| 108.0 | 0.59 | 0.54 | 0.59 | 0.56 | 0.54 | 0.49 | 0.01 | 0.02 | 0.59 | 0.58 |
| 108.5 | 0.58 | 0.54 | 0.58 | 0.53 | 0.56 | 0.51 | 0.09 | -0.01 | 0.54 | 0.60 |
| 109.0 | 0.57 | 0.50 | 0.57 | 0.56 | 0.55 | 0.54 | 0.18 | 0.04 | 0.56 | 0.60 |
| 109.5 | 0.57 | 0.46 | 0.57 | 0.51 | 0.55 | 0.54 | 0.29 | 0.15 | 0.57 | 0.61 |
| 110.0 | 0.53 | 0.35 | 0.57 | 0.51 | 0.53 | 0.53 | 0.37 | 0.32 | 0.56 | 0.62 |
| 110.5 | 0.52 | 0.23 | 0.56 | 0.45 | 0.55 | 0.55 | 0.47 | 0.43 | 0.58 | 0.62 |
| 111.0 | 0.48 | 0.09 | 0.55 | 0.35 | 0.54 | 0.55 | 0.50 | 0.47 | 0.57 | 0.58 |
| 111.5 | 0.46 | -0.02 | 0.57 | 0.24 | 0.54 | 0.54 | 0.52 | 0.47 | 0.58 | 0.57 |
| 112.0 | 0.40 | -0.03 | 0.55 | 0.05 | 0.55 | 0.54 | 0.49 | 0.52 | 0.57 | 0.61 |
| 112.5 | 0.25 | 0.10 | 0.57 | -0.01 | 0.51 | 0.50 | 0.44 | 0.51 | 0.55 | 0.62 |
| 113.0 | 0.11 | 0.26 | 0.54 | 0.02 | 0.51 | 0.52 | 0.48 | 0.50 | 0.54 | 0.58 |
| 113.5 | 0.02 | 0.39 | 0.55 | 0.13 | 0.48 | 0.56 | 0.52 | 0.49 | 0.52 | 0.65 |
| 114.0 | -0.01 | 0.46 | 0.54 | 0.30 | 0.43 | 0.57 | 0.57 | 0.44 | 0.52 | 0.56 |
| 114.5 | 0.01 | 0.46 | 0.54 | 0.41 | 0.37 | 0.58 | 0.59 | 0.46 | 0.50 | 0.58 |
| 115.0 | 0.15 | 0.54 | 0.50 | 0.46 | 0.24 | 0.59 | 0.60 | 0.50 | 0.49 | 0.61 |
| 115.5 | 0.28 | 0.52 | 0.47 | 0.51 | 0.09 | 0.62 | 0.62 | 0.55 | 0.42 | 0.59 |
| 116.0 | 0.36 | 0.56 | 0.40 | 0.53 | 0.04 | 0.61 | 0.58 | 0.55 | 0.34 | 0.56 |
| 116.5 | 0.44 | 0.50 | 0.30 | 0.52 | 0.01 | 0.60 | 0.61 | 0.58 | 0.22 | 0.60 |
| 117.0 | 0.45 | 0.55 | 0.14 | 0.52 | 0.12 | 0.62 | 0.60 | 0.58 | 0.05 | 0.54 |
| 117.5 | 0.49 | 0.48 | 0.02 | 0.45 | 0.26 | 0.59 | 0.55 | 0.57 | -0.01 | 0.50 |
| 118.0 | 0.49 | 0.50 | 0.01 | 0.53 | 0.37 | 0.62 | 0.60 | 0.58 | -0.02 | 0.49 |
| 118.5 | 0.50 | 0.52 | 0.05 | 0.55 | 0.39 | 0.61 | 0.65 | 0.59 | 0.10 | 0.51 |
| 119.0 | 0.49 | 0.56 | 0.24 | 0.54 | 0.41 | 0.62 | 0.60 | 0.56 | 0.25 | 0.53 |
| 119.5 | 0.48 | 0.58 | 0.36 | 0.55 | 0.42 | 0.58 | 0.61 | 0.57 | 0.38 | 0.53 |
| 120.0 | 0.52 | 0.60 | 0.42 | 0.55 | 0.50 | 0.60 | 0.62 | 0.58 | 0.43 | 0.55 |
| 120.5 | 0.54 | 0.59 | 0.44 | 0.58 | 0.51 | 0.60 | 0.60 | 0.54 | 0.45 | 0.53 |
| 121.0 | 0.59 | 0.57 | 0.47 | 0.57 | 0.53 | 0.62 | 0.57 | 0.60 | 0.48 | 0.50 |
| 121.5 | 0.60 | 0.60 | 0.48 | 0.58 | 0.56 | 0.62 | 0.60 | 0.55 | 0.48 | 0.55 |
| 122.0 | 0.57 | 0.59 | 0.51 | 0.58 | 0.53 | 0.62 | 0.59 | 0.56 | 0.47 | 0.47 |
| 122.5 | 0.57 | 0.56 | 0.47 | 0.60 | 0.53 | 0.58 | 0.61 | 0.60 | 0.46 | 0.44 |
| 123.0 | 0.60 | 0.60 | 0.46 | 0.61 | 0.56 | 0.58 | 0.61 | 0.59 | 0.50 | 0.34 |
| 123.5 | 0.61 | 0.58 | 0.50 | 0.59 | 0.57 | 0.61 | 0.62 | 0.58 | 0.53 | 0.26 |
| 124.0 | 0.60 | 0.60 | 0.52 | 0.60 | 0.55 | 0.61 | 0.61 | 0.56 | 0.55 | 0.13 |
| 124.5 | 0.60 | 0.62 | 0.54 | 0.60 | 0.56 | 0.61 | 0.62 | 0.57 | 0.56 | 0.00 |
| 125.0 | 0.59 | 0.60 | 0.57 | 0.56 | 0.57 | 0.60 | 0.60 | 0.55 | 0.54 | 0.00 |
| 125.5 | 0.60 | 0.61 | 0.58 | 0.59 | 0.55 | 0.61 | 0.57 | 0.61 | 0.59 | 0.03 |
| 126.0 | 0.58 | 0.60 | 0.58 | 0.59 | 0.55 | 0.60 | 0.65 | 0.54 | 0.55 | 0.13 |
| 126.5 | 0.60 | 0.60 | 0.58 | 0.61 | 0.56 | 0.59 | 0.63 | 0.58 | 0.56 | 0.38 |
| 127.0 | 0.60 | 0.63 | 0.55 | 0.57 | 0.55 | 0.63 | 0.63 | 0.56 | 0.57 | 0.42 |
| 127.5 | 0.62 | 0.56 | 0.57 | 0.63 | 0.56 | 0.62 | 0.64 | 0.59 | 0.57 | 0.50 |
| 128.0 | 0.61 | 0.60 | 0.58 | 0.59 | 0.53 | 0.60 | 0.61 | 0.60 | 0.57 | 0.48 |
| 128.5 | 0.62 | 0.59 | 0.58 | 0.59 | 0.57 | 0.59 | 0.63 | 0.58 | 0.58 | 0.54 |
| 129.0 | 0.62 | 0.60 | 0.57 | 0.57 | 0.54 | 0.62 | 0.62 | 0.57 | 0.57 | 0.56 |
| 129.5 | 0.60 | 0.60 | 0.57 | 0.59 | 0.57 | 0.63 | 0.62 | 0.55 | 0.59 | 0.56 |
| 130.0 | 0.64 | 0.60 | 0.59 | 0.58 | 0.55 | 0.61 | 0.65 | 0.57 | 0.60 | 0.59 |
| 130.5 | 0.60 | 0.58 | 0.58 | 0.60 | 0.55 | 0.59 | 0.63 | 0.51 | 0.59 | 0.56 |
| 131.0 | 0.62 | 0.61 | 0.59 | 0.60 | 0.55 | 0.60 | 0.66 | 0.59 | 0.54 | 0.55 |
| 131.5 | 0.59 | 0.56 | 0.58 | 0.59 | 0.58 | 0.62 | 0.62 | 0.58 | 0.56 | 0.60 |
| 132.0 | 0.61 | 0.58 | 0.58 | 0.64 | 0.53 | 0.60 | 0.62 | 0.59 | 0.57 | 0.63 |
| 132.5 | 0.57 | 0.59 | 0.57 | 0.59 | 0.58 | 0.60 | 0.63 | 0.58 | 0.59 | 0.59 |
| 133.0 | 0.59 | 0.59 | 0.57 | 0.59 | 0.56 | 0.61 | 0.61 | 0.57 | 0.59 | 0.64 |
| 133.5 | 0.60 | 0.56 | 0.58 | 0.58 | 0.54 | 0.61 | 0.61 | 0.58 | 0.58 | 0.62 |
| 134.0 | 0.59 | 0.60 | 0.59 | 0.57 | 0.55 | 0.62 | 0.59 | 0.57 | 0.59 | 0.63 |

**Table S4.** ^15^N CEST I/I_0_ data measured for FUS-RRM in 1M urea at pH 6.4.

|  | **I/I_0_** | | | | | | | | | | | | | |
| --- | --- | --- | --- | --- | --- | --- | --- | --- | --- | --- | --- | --- | --- | --- |
| **^15^N offset (ppm)** | **G293** | **E294** | **T297** | **E299** | **S300** | **D303** | **F305** | **Q307** | **T317** | **T330** | **T338** | **D352** | **D355** | **G356** |
| 102.0 | 0.48 | 0.58 | 0.54 | 0.52 | 0.58 | 0.56 | 0.54 | 0.54 | 0.52 | 0.54 | 0.54 | 0.53 | 0.56 | 0.53 |
| 102.5 | 0.47 | 0.58 | 0.54 | 0.54 | 0.57 | 0.55 | 0.54 | 0.55 | 0.50 | 0.54 | 0.53 | 0.54 | 0.55 | 0.51 |
| 103.0 | 0.42 | 0.58 | 0.52 | 0.54 | 0.56 | 0.56 | 0.53 | 0.54 | 0.49 | 0.54 | 0.50 | 0.55 | 0.56 | 0.54 |
| 103.5 | 0.35 | 0.58 | 0.54 | 0.54 | 0.56 | 0.56 | 0.53 | 0.55 | 0.44 | 0.51 | 0.50 | 0.54 | 0.56 | 0.54 |
| 104.0 | 0.22 | 0.57 | 0.55 | 0.55 | 0.56 | 0.55 | 0.55 | 0.54 | 0.38 | 0.51 | 0.52 | 0.53 | 0.53 | 0.51 |
| 104.5 | 0.05 | 0.55 | 0.54 | 0.54 | 0.56 | 0.55 | 0.54 | 0.53 | 0.22 | 0.45 | 0.52 | 0.55 | 0.54 | 0.53 |
| 105.0 | 0.02 | 0.57 | 0.52 | 0.53 | 0.57 | 0.55 | 0.52 | 0.53 | 0.07 | 0.39 | 0.49 | 0.54 | 0.53 | 0.55 |
| 105.5 | 0.05 | 0.57 | 0.51 | 0.54 | 0.57 | 0.56 | 0.53 | 0.58 | 0.03 | 0.33 | 0.48 | 0.54 | 0.54 | 0.52 |
| 106.0 | 0.19 | 0.58 | 0.53 | 0.55 | 0.57 | 0.55 | 0.54 | 0.54 | 0.07 | 0.14 | 0.44 | 0.54 | 0.54 | 0.52 |
| 106.5 | 0.29 | 0.55 | 0.54 | 0.50 | 0.56 | 0.55 | 0.54 | 0.55 | 0.25 | 0.00 | 0.41 | 0.55 | 0.56 | 0.50 |
| 107.0 | 0.39 | 0.56 | 0.53 | 0.52 | 0.56 | 0.54 | 0.54 | 0.53 | 0.36 | 0.01 | 0.30 | 0.53 | 0.55 | 0.52 |
| 107.5 | 0.42 | 0.57 | 0.52 | 0.54 | 0.56 | 0.55 | 0.56 | 0.52 | 0.46 | 0.08 | 0.20 | 0.55 | 0.54 | 0.51 |
| 108.0 | 0.46 | 0.59 | 0.49 | 0.53 | 0.56 | 0.56 | 0.55 | 0.56 | 0.48 | 0.23 | 0.04 | 0.55 | 0.55 | 0.55 |
| 108.5 | 0.41 | 0.56 | 0.49 | 0.55 | 0.57 | 0.56 | 0.50 | 0.53 | 0.48 | 0.36 | 0.01 | 0.53 | 0.54 | 0.50 |
| 109.0 | 0.38 | 0.55 | 0.45 | 0.54 | 0.56 | 0.54 | 0.49 | 0.53 | 0.50 | 0.44 | 0.08 | 0.54 | 0.56 | 0.38 |
| 109.5 | 0.42 | 0.55 | 0.43 | 0.56 | 0.55 | 0.55 | 0.49 | 0.54 | 0.54 | 0.48 | 0.20 | 0.53 | 0.55 | 0.40 |
| 110.0 | 0.48 | 0.53 | 0.36 | 0.55 | 0.56 | 0.54 | 0.46 | 0.54 | 0.54 | 0.51 | 0.33 | 0.53 | 0.55 | 0.47 |
| 110.5 | 0.48 | 0.52 | 0.24 | 0.54 | 0.55 | 0.54 | 0.41 | 0.52 | 0.53 | 0.49 | 0.35 | 0.52 | 0.55 | 0.46 |
| 111.0 | 0.53 | 0.49 | 0.07 | 0.51 | 0.55 | 0.52 | 0.34 | 0.54 | 0.52 | 0.54 | 0.44 | 0.51 | 0.56 | 0.48 |
| 111.5 | 0.51 | 0.45 | 0.01 | 0.53 | 0.55 | 0.51 | 0.22 | 0.51 | 0.55 | 0.53 | 0.44 | 0.53 | 0.54 | 0.46 |
| 112.0 | 0.51 | 0.40 | 0.05 | 0.51 | 0.54 | 0.51 | 0.07 | 0.49 | 0.54 | 0.51 | 0.48 | 0.53 | 0.52 | 0.47 |
| 112.5 | 0.50 | 0.29 | 0.21 | 0.52 | 0.52 | 0.50 | 0.00 | 0.49 | 0.49 | 0.52 | 0.50 | 0.52 | 0.53 | 0.42 |
| 113.0 | 0.52 | 0.10 | 0.33 | 0.52 | 0.50 | 0.46 | 0.04 | 0.48 | 0.47 | 0.48 | 0.47 | 0.50 | 0.53 | 0.43 |
| 113.5 | 0.52 | 0.01 | 0.39 | 0.52 | 0.47 | 0.44 | 0.19 | 0.45 | 0.46 | 0.44 | 0.46 | 0.52 | 0.55 | 0.39 |
| 114.0 | 0.52 | -0.01 | 0.46 | 0.52 | 0.42 | 0.37 | 0.32 | 0.43 | 0.47 | 0.50 | 0.42 | 0.50 | 0.54 | 0.27 |
| 114.5 | 0.52 | 0.12 | 0.49 | 0.51 | 0.33 | 0.24 | 0.38 | 0.39 | 0.52 | 0.52 | 0.43 | 0.49 | 0.51 | 0.19 |
| 115.0 | 0.55 | 0.28 | 0.50 | 0.49 | 0.17 | 0.06 | 0.45 | 0.27 | 0.51 | 0.56 | 0.46 | 0.48 | 0.51 | 0.03 |
| 115.5 | 0.55 | 0.42 | 0.49 | 0.45 | 0.02 | 0.01 | 0.44 | 0.06 | 0.55 | 0.56 | 0.52 | 0.45 | 0.47 | 0.01 |
| 116.0 | 0.53 | 0.46 | 0.52 | 0.42 | 0.01 | 0.05 | 0.46 | 0.00 | 0.56 | 0.57 | 0.53 | 0.42 | 0.47 | 0.03 |
| 116.5 | 0.55 | 0.48 | 0.50 | 0.38 | 0.10 | 0.21 | 0.49 | 0.01 | 0.54 | 0.57 | 0.54 | 0.31 | 0.41 | 0.14 |
| 117.0 | 0.50 | 0.52 | 0.50 | 0.27 | 0.27 | 0.31 | 0.49 | 0.14 | 0.55 | 0.57 | 0.55 | 0.18 | 0.34 | 0.30 |
| 117.5 | 0.56 | 0.49 | 0.45 | 0.11 | 0.38 | 0.39 | 0.46 | 0.29 | 0.54 | 0.59 | 0.53 | 0.02 | 0.17 | 0.41 |
| 118.0 | 0.52 | 0.51 | 0.41 | 0.00 | 0.46 | 0.40 | 0.40 | 0.37 | 0.57 | 0.57 | 0.50 | 0.02 | 0.03 | 0.44 |
| 118.5 | 0.54 | 0.53 | 0.44 | 0.03 | 0.48 | 0.39 | 0.45 | 0.40 | 0.54 | 0.58 | 0.53 | 0.05 | 0.01 | 0.45 |
| 119.0 | 0.55 | 0.51 | 0.50 | 0.15 | 0.49 | 0.46 | 0.50 | 0.40 | 0.59 | 0.57 | 0.54 | 0.20 | 0.04 | 0.48 |
| 119.5 | 0.54 | 0.46 | 0.52 | 0.30 | 0.51 | 0.50 | 0.50 | 0.40 | 0.52 | 0.58 | 0.56 | 0.35 | 0.19 | 0.51 |
| 120.0 | 0.53 | 0.45 | 0.53 | 0.39 | 0.51 | 0.52 | 0.53 | 0.42 | 0.55 | 0.60 | 0.50 | 0.40 | 0.33 | 0.49 |
| 120.5 | 0.53 | 0.50 | 0.54 | 0.41 | 0.50 | 0.52 | 0.52 | 0.49 | 0.56 | 0.60 | 0.50 | 0.43 | 0.37 | 0.48 |
| 121.0 | 0.54 | 0.53 | 0.52 | 0.45 | 0.49 | 0.54 | 0.57 | 0.52 | 0.57 | 0.57 | 0.53 | 0.45 | 0.41 | 0.53 |
| 121.5 | 0.52 | 0.55 | 0.52 | 0.48 | 0.51 | 0.54 | 0.52 | 0.50 | 0.57 | 0.58 | 0.54 | 0.47 | 0.42 | 0.53 |
| 122.0 | 0.51 | 0.55 | 0.55 | 0.49 | 0.54 | 0.54 | 0.57 | 0.52 | 0.56 | 0.58 | 0.56 | 0.48 | 0.40 | 0.51 |
| 122.5 | 0.53 | 0.57 | 0.55 | 0.46 | 0.55 | 0.54 | 0.55 | 0.53 | 0.56 | 0.58 | 0.54 | 0.47 | 0.46 | 0.51 |
| 123.0 | 0.53 | 0.58 | 0.52 | 0.46 | 0.55 | 0.54 | 0.56 | 0.54 | 0.58 | 0.58 | 0.57 | 0.44 | 0.50 | 0.49 |
| 123.5 | 0.55 | 0.56 | 0.54 | 0.45 | 0.56 | 0.55 | 0.53 | 0.51 | 0.58 | 0.59 | 0.53 | 0.43 | 0.52 | 0.51 |
| 124.0 | 0.56 | 0.56 | 0.55 | 0.42 | 0.56 | 0.55 | 0.53 | 0.56 | 0.58 | 0.59 | 0.54 | 0.50 | 0.52 | 0.52 |
| 124.5 | 0.54 | 0.56 | 0.55 | 0.46 | 0.56 | 0.56 | 0.56 | 0.52 | 0.55 | 0.62 | 0.53 | 0.50 | 0.53 | 0.52 |
| 125.0 | 0.52 | 0.56 | 0.54 | 0.51 | 0.56 | 0.56 | 0.53 | 0.53 | 0.57 | 0.60 | 0.54 | 0.54 | 0.56 | 0.50 |
| 125.5 | 0.53 | 0.58 | 0.53 | 0.51 | 0.58 | 0.55 | 0.55 | 0.55 | 0.58 | 0.59 | 0.55 | 0.54 | 0.55 | 0.51 |
| 126.0 | 0.55 | 0.57 | 0.54 | 0.53 | 0.58 | 0.55 | 0.54 | 0.55 | 0.57 | 0.61 | 0.51 | 0.53 | 0.55 | 0.52 |
| 126.5 | 0.53 | 0.59 | 0.55 | 0.52 | 0.56 | 0.56 | 0.57 | 0.51 | 0.56 | 0.59 | 0.54 | 0.52 | 0.54 | 0.49 |
| 127.0 | 0.50 | 0.57 | 0.56 | 0.52 | 0.56 | 0.55 | 0.53 | 0.53 | 0.56 | 0.59 | 0.54 | 0.54 | 0.54 | 0.50 |
| 127.5 | 0.55 | 0.57 | 0.55 | 0.52 | 0.57 | 0.56 | 0.57 | 0.56 | 0.58 | 0.55 | 0.52 | 0.53 | 0.55 | 0.52 |
| 128.0 | 0.54 | 0.57 | 0.53 | 0.54 | 0.57 | 0.57 | 0.54 | 0.54 | 0.57 | 0.56 | 0.57 | 0.53 | 0.54 | 0.55 |
| 128.5 | 0.55 | 0.57 | 0.54 | 0.53 | 0.56 | 0.57 | 0.54 | 0.54 | 0.58 | 0.59 | 0.53 | 0.53 | 0.55 | 0.53 |
| 129.0 | 0.52 | 0.57 | 0.53 | 0.55 | 0.57 | 0.56 | 0.52 | 0.54 | 0.55 | 0.58 | 0.53 | 0.55 | 0.55 | 0.53 |
| 129.5 | 0.55 | 0.58 | 0.54 | 0.54 | 0.56 | 0.57 | 0.55 | 0.54 | 0.55 | 0.57 | 0.51 | 0.53 | 0.53 | 0.53 |
| 130.0 | 0.55 | 0.58 | 0.54 | 0.53 | 0.58 | 0.54 | 0.56 | 0.54 | 0.55 | 0.58 | 0.52 | 0.56 | 0.55 | 0.50 |
| 130.5 | 0.54 | 0.58 | 0.54 | 0.53 | 0.57 | 0.58 | 0.55 | 0.55 | 0.57 | 0.58 | 0.55 | 0.54 | 0.57 | 0.50 |
| 131.0 | 0.54 | 0.57 | 0.53 | 0.54 | 0.57 | 0.55 | 0.52 | 0.54 | 0.57 | 0.62 | 0.53 | 0.56 | 0.55 | 0.53 |
| 131.5 | 0.56 | 0.57 | 0.53 | 0.52 | 0.57 | 0.55 | 0.51 | 0.53 | 0.58 | 0.60 | 0.53 | 0.55 | 0.55 | 0.55 |
| 132.0 | 0.55 | 0.58 | 0.56 | 0.56 | 0.56 | 0.57 | 0.54 | 0.55 | 0.55 | 0.57 | 0.54 | 0.55 | 0.56 | 0.54 |
| 132.5 | 0.53 | 0.59 | 0.54 | 0.54 | 0.56 | 0.56 | 0.58 | 0.53 | 0.56 | 0.57 | 0.55 | 0.55 | 0.56 | 0.56 |
| 133.0 | 0.53 | 0.57 | 0.54 | 0.56 | 0.57 | 0.55 | 0.55 | 0.53 | 0.56 | 0.57 | 0.53 | 0.54 | 0.56 | 0.54 |

**Table S5.** ^15^N CEST I/I_0_ data measured for FUS-RRM at pH 6.4 in the presence of ATP.

|  | **I/I_0_** | | | | | | | | | | | | |
| --- | --- | --- | --- | --- | --- | --- | --- | --- | --- | --- | --- | --- | --- |
| **^15^N offset (ppm)** | **N285** | **E294** | **T297** | **E299** | **S300** | **F305** | **Q307** | **T317** | **T330** | **T338** | **D352** | **D355** | **G356** |
| 101.17 | 0.58 | 0.61 | 0.63 | 0.56 | 0.59 | 0.60 | 0.57 | 0.57 | 0.60 | 0.55 | 0.60 | 0.59 | 0.61 |
| 102.49 | 0.58 | 0.62 | 0.60 | 0.58 | 0.60 | 0.60 | 0.58 | 0.48 | 0.55 | 0.53 | 0.64 | 0.57 | 0.60 |
| 103.81 | 0.59 | 0.61 | 0.62 | 0.59 | 0.59 | 0.59 | 0.56 | 0.15 | 0.41 | 0.52 | 0.64 | 0.57 | 0.57 |
| 105.13 | 0.55 | 0.62 | 0.60 | 0.60 | 0.60 | 0.57 | 0.57 | 0.06 | 0.12 | 0.47 | 0.62 | 0.60 | 0.61 |
| 106.45 | 0.58 | 0.61 | 0.61 | 0.57 | 0.60 | 0.58 | 0.55 | 0.43 | -0.02 | 0.19 | 0.64 | 0.53 | 0.55 |
| 107.77 | 0.58 | 0.58 | 0.55 | 0.55 | 0.59 | 0.58 | 0.59 | 0.54 | 0.32 | 0.01 | 0.62 | 0.56 | 0.55 |
| 109.09 | 0.56 | 0.55 | 0.40 | 0.59 | 0.58 | 0.50 | 0.56 | 0.59 | 0.51 | 0.36 | 0.62 | 0.58 | 0.54 |
| 110.40 | 0.57 | 0.47 | -0.01 | 0.57 | 0.57 | 0.32 | 0.54 | 0.58 | 0.55 | 0.49 | 0.61 | 0.56 | 0.53 |
| 111.72 | 0.53 | 0.22 | 0.48 | 0.59 | 0.55 | 0.01 | 0.51 | 0.56 | 0.56 | 0.54 | 0.60 | 0.57 | 0.46 |
| 113.04 | 0.53 | 0.01 | 0.57 | 0.50 | 0.43 | 0.51 | 0.46 | 0.59 | 0.58 | 0.53 | 0.56 | 0.58 | 0.30 |
| 114.36 | 0.53 | 0.51 | 0.59 | 0.49 | 0.02 | 0.54 | 0.20 | 0.61 | 0.61 | 0.57 | 0.49 | 0.52 | -0.01 |
| 115.68 | 0.51 | 0.57 | 0.56 | 0.34 | 0.51 | 0.55 | 1.80025652E-4 | 0.61 | 0.62 | 0.55 | 0.27 | 0.40 | 0.44 |
| 117.00 | 0.46 | 0.57 | 0.60 | 0.01 | 0.55 | 0.58 | 0.47 | 0.61 | 0.65 | 0.54 | 0.01 | 0.02 | 0.48 |
| 118.32 | 0.15 | 0.57 | 0.62 | 0.49 | 0.56 | 0.61 | 0.55 | 0.61 | 0.63 | 0.57 | 0.49 | 0.11 | 0.59 |
| 119.64 | 0.03 | 0.60 | 0.62 | 0.50 | 0.59 | 0.62 | 0.55 | 0.62 | 0.64 | 0.57 | 0.54 | 0.43 | 0.56 |
| 120.96 | 0.38 | 0.61 | 0.61 | 0.53 | 0.59 | 0.57 | 0.55 | 0.63 | 0.62 | 0.61 | 0.56 | 0.48 | 0.58 |
| 122.28 | 0.49 | 0.62 | 0.63 | 0.59 | 0.59 | 0.64 | 0.55 | 0.63 | 0.65 | 0.57 | 0.61 | 0.56 | 0.61 |
| 123.60 | 0.54 | 0.61 | 0.61 | 0.55 | 0.60 | 0.59 | 0.57 | 0.60 | 0.64 | 0.57 | 0.60 | 0.57 | 0.58 |
| 124.91 | 0.56 | 0.62 | 0.63 | 0.56 | 0.60 | 0.65 | 0.57 | 0.60 | 0.63 | 0.57 | 0.62 | 0.58 | 0.58 |
| 126.23 | 0.55 | 0.62 | 0.65 | 0.57 | 0.60 | 0.62 | 0.57 | 0.60 | 0.64 | 0.57 | 0.62 | 0.57 | 0.61 |
| 127.55 | 0.57 | 0.60 | 0.60 | 0.58 | 0.60 | 0.61 | 0.56 | 0.63 | 0.63 | 0.61 | 0.63 | 0.58 | 0.57 |
| 128.87 | 0.56 | 0.61 | 0.63 | 0.59 | 0.59 | 0.60 | 0.56 | 0.63 | 0.63 | 0.59 | 0.61 | 0.57 | 0.59 |
| 130.19 | 0.58 | 0.62 | 0.66 | 0.62 | 0.59 | 0.62 | 0.57 | 0.63 | 0.64 | 0.58 | 0.62 | 0.57 | 0.64 |
| 131.51 | 0.54 | 0.63 | 0.60 | 0.58 | 0.61 | 0.60 | 0.57 | 0.62 | 0.64 | 0.58 | 0.62 | 0.58 | 0.61 |
| 132.83 | 0.58 | 0.63 |  | 0.58 |  |  | 0.57 | 0.64 | 0.64 | 0.58 | 0.62 | 0.60 |  |

**Table S6.** ^15^N CEST I/I_0_ data for FUS-RRM at pH 4.6 in the presence of ATP.

|  | **I/I_0_** | | | | | | | | |
| --- | --- | --- | --- | --- | --- | --- | --- | --- | --- |
| **^15^N offset (ppm)** | **E294** | **T297** | **F305** | **Q307** | **T317** | **T330** | **T338** | **D352** | **S360** |
| 101.17 | 0.61 | 0.60 | 0.60 | 0.59 | 0.51 | 0.53 | 0.56 | 0.56 | 0.62 |
| 102.49 | 0.58 | 0.59 | 0.58 | 0.57 | 0.43 | 0.51 | 0.57 | 0.57 | 0.62 |
| 103.81 | 0.61 | 0.58 | 0.55 | 0.58 | 0.14 | 0.36 | 0.51 | 0.57 | 0.61 |
| 105.13 | 0.57 | 0.57 | 0.55 | 0.57 | 0.04 | 0.19 | 0.43 | 0.58 | 0.65 |
| 106.45 | 0.58 | 0.55 | 0.56 | 0.58 | 0.40 | 0.00 | 0.16 | 0.56 | 0.59 |
| 107.77 | 0.56 | 0.52 | 0.53 | 0.59 | 0.51 | 0.04 | -0.02 | 0.58 | 0.61 |
| 109.09 | 0.53 | 0.40 | 0.51 | 0.58 | 0.54 | 0.31 | 0.51 | 0.56 | 0.68 |
| 110.40 | 0.44 | 0.01 | 0.31 | 0.57 | 0.56 | 0.44 | 0.53 | 0.55 | 0.61 |
| 111.72 | 0.19 | 0.12 | -0.02 | 0.54 | 0.51 | 0.47 | 0.50 | 0.54 | 0.63 |
| 113.04 | -0.01 | 0.43 | 0.48 | 0.45 | 0.58 | 0.52 | 0.59 | 0.52 | 0.62 |
| 114.36 | 0.42 | 0.52 | 0.48 | 0.17 | 0.60 | 0.57 | 0.58 | 0.42 | 0.61 |
| 115.68 | 0.49 | 0.55 | 0.49 | -0.01 | 0.59 | 0.60 | 0.58 | 0.16 | 0.60 |
| 117.00 | 0.47 | 0.50 | 0.55 | 0.43 | 0.60 | 0.62 | 0.59 | -0.01 | 0.53 |
| 119.64 | 0.55 | 0.56 | 0.56 | 0.52 | 0.61 | 0.62 | 0.61 | 0.45 | 0.56 |
| 120.96 | 0.60 | 0.62 | 0.58 | 0.56 | 0.61 | 0.65 | 0.59 | 0.47 | 0.56 |
| 122.28 | 0.63 | 0.65 | 0.57 | 0.55 | 0.61 | 0.61 | 0.57 | 0.52 | 0.49 |
| 123.60 | 0.58 | 0.61 | 0.58 | 0.57 | 0.60 | 0.64 | 0.57 | 0.56 | 0.33 |
| 124.91 | 0.61 | 0.60 | 0.55 | 0.59 | 0.61 | 0.62 | 0.60 | 0.57 | 0.04 |
| 126.23 | 0.60 | 0.63 | 0.56 | 0.58 | 0.60 | 0.58 | 0.59 | 0.57 | 0.08 |
| 127.55 | 0.61 | 0.62 | 0.62 | 0.58 | 0.64 | 0.62 | 0.58 | 0.58 | 0.45 |
| 128.87 | 0.61 | 0.63 | 0.60 | 0.59 | 0.61 | 0.61 | 0.58 | 0.59 | 0.53 |
| 130.19 | 0.63 | 0.64 | 0.60 | 0.61 | 0.63 | 0.65 | 0.60 | 0.58 | 0.58 |
| 131.51 | 0.60 | 0.59 | 0.59 | 0.60 | 0.64 | 0.64 | 0.60 | 0.57 | 0.60 |
| 132.83 | 0.60 | 0.61 | 0.60 | 0.60 | 0.59 | 0.64 | 0.63 | 0.59 | 0.61 |

**Table S7.** *R_1_* relaxation rates measured from HARD experiment for FUS-RRM (pH 6.4) with ATP at 600 MHz NMR spectrometer. Only data for which reliable fits were obtained have been included.

| **Residue number** | **R_1_ (s^-1^)** | |
| --- | --- | --- |
|  | **Value** | **Error** |
| 285 | 1.43 | 0.04 |
| 287 | 1.58 | 0.12 |
| 288 | 1.51 | 0.04 |
| 289 | 1.38 | 0.14 |
| 290 | 1.32 | 0.03 |
| 294 | 1.26 | 0.04 |
| 295 | 1.36 | 0.03 |
| 296 | 1.19 | 0.05 |
| 297 | 1.36 | 0.03 |
| 298 | 1.41 | 0.04 |
| 300 | 1.32 | 0.05 |
| 301 | 1.44 | 0.05 |
| 302 | 1.64 | 0.05 |
| 303 | 0.99 | 0.02 |
| 304 | 1.45 | 0.02 |
| 305 | 1.52 | 0.04 |
| 306 | 1.31 | 0.06 |
| 307 | 1.53 | 0.02 |
| 308 | 1.59 | 0.04 |
| 309 | 1.41 | 0.03 |
| 310 | 1.21 | 0.03 |
| 311 | 1.31 | 0.07 |
| 313 | 1.45 | 0.03 |
| 314 | 1.29 | 0.06 |
| 317 | 1.35 | 0.03 |
| 318 | 1.39 | 0.02 |
| 319 | 1.44 | 0.02 |
| 322 | 1.42 | 0.05 |
| 323 | 1.69 | 0.04 |
| 324 | 1.37 | 0.04 |
| 326 | 1.31 | 0.05 |
| 327 | 1.40 | 0.03 |
| 329 | 1.24 | 0.02 |
| 330 | 1.25 | 0.04 |
| 331 | 1.36 | 0.04 |
| 332 | 1.58 | 0.01 |
| 335 | 1.32 | 0.08 |
| 338 | 1.40 | 0.04 |
| 340 | 1.45 | 0.04 |
| 341 | 1.41 | 0.05 |
| 346 | 1.37 | 0.02 |
| 347 | 1.41 | 0.07 |
| 349 | 1.54 | 0.03 |
| 350 | 1.50 | 0.07 |
| 351 | 1.43 | 0.04 |
| 352 | 1.41 | 0.03 |
| 353 | 1.39 | 0.04 |
| 355 | 1.42 | 0.02 |
| 356 | 1.47 | 0.03 |
| 358 | 1.61 | 0.02 |
| 360 | 1.60 | 0.04 |
| 361 | 1.36 | 0.04 |
| 362 | 1.40 | 0.02 |
| 367 | 1.47 | 0.02 |
| 369 | 1.44 | 0.18 |
| 372 | 1.76 | 0.05 |

**Table S8**: *R_1ρ_* relaxation rates measured using HSn pulses (n=1,2,4,6,8) from HARD experiment for FUS-RRM (pH 6.4) with ATP at 600 MHz NMR spectrometer.

| **Residue**  **Number** | ***R_1ρ_* (s^-1^)** | | | | | | | | | |
| --- | --- | --- | --- | --- | --- | --- | --- | --- | --- | --- |
|  | **HS1** | | **HS2** | | **HS4** | | **HS6** | | **HS8** | |
|  | **Value** | **Error** | **Value** | **Error** | **Value** | **Error** | **Value** | **Error** | **Value** | **Error** |
| 285 | 4.23 | 0.10 | 4.91 | 0.22 | 5.59 | 0.30 | 5.28 | 0.14 | 6.25 | 0.23 |
| 287 | 4.18 | 0.49 | 4.97 | 0.28 | 6.42 | 0.56 | 4.78 | 0.38 | 5.46 | 0.39 |
| 288 | 3.87 | 0.17 | 5.23 | 0.54 | 5.84 | 0.43 | 5.52 | 0.27 | 5.81 | 0.11 |
| 289 | 4.62 | 0.45 | 5.40 | 1.56 | 7.69 | 0.55 | 6.67 | 0.33 | 8.26 | 1.13 |
| 290 | 3.53 | 0.06 | 4.08 | 0.13 | 5.51 | 0.11 | 4.95 | 0.10 | 5.13 | 0.10 |
| 294 | 5.52 | 0.34 | 6.18 | 0.26 | 7.34 | 0.18 | 6.93 | 0.07 | 7.54 | 0.22 |
| 295 | 3.58 | 0.11 | 4.62 | 0.53 | 5.87 | 0.20 | 5.20 | 0.30 | 5.17 | 0.24 |
| 296 | 4.20 | 0.22 | 4.37 | 0.12 | 5.25 | 0.18 | 5.11 | 0.26 | 5.33 | 0.23 |
| 297 | 3.74 | 0.10 | 3.96 | 0.27 | 4.27 | 0.43 | 4.20 | 0.22 | 5.23 | 0.09 |
| 298 | 3.97 | 0.23 | 4.70 | 0.17 | 4.73 | 0.71 | 5.24 | 0.18 | 6.01 | 0.17 |
| 300 | 4.54 | 0.43 | 4.62 | 0.28 | 6.49 | 0.45 | 5.59 | 0.30 | 6.00 | 0.28 |
| 301 | 4.84 | 0.19 | 5.67 | 0.39 | 6.43 | 0.43 | 6.27 | 0.34 | 7.31 | 0.41 |
| 302 | 4.99 | 0.43 | 5.86 | 1.16 | 7.21 | 0.46 | 6.66 | 0.35 | 6.87 | 0.42 |
| 303 | 3.81 | 0.14 | 3.29 | 0.68 | 2.31 | 0.55 | 2.39 | 0.37 | 4.36 | 0.11 |
| 304 | 4.77 | 0.28 | 5.00 | 0.50 | 6.16 | 0.30 | 5.88 | 0.26 | 6.11 | 0.26 |
| 305 | 4.61 | 0.22 | 5.17 | 0.47 | 5.78 | 0.33 | 5.74 | 0.15 | 6.08 | 0.18 |
| 306 | 4.24 | 0.30 | 4.45 | 0.51 | 5.33 | 0.54 | 4.78 | 0.14 | 5.37 | 0.20 |
| 307 | 5.03 | 0.18 | 6.15 | 0.37 | 6.81 | 0.50 | 7.18 | 0.33 | 7.34 | 0.33 |
| 308 | 4.46 | 0.23 | 5.31 | 0.42 | 6.56 | 0.47 | 6.56 | 0.16 | 6.33 | 0.25 |
| 309 | 3.92 | 0.07 | 4.85 | 0.48 | 5.68 | 0.17 | 5.46 | 0.10 | 5.38 | 0.17 |
| 310 | 4.61 | 0.16 | 5.17 | 0.56 | 6.33 | 0.34 | 6.12 | 0.25 | 6.60 | 0.18 |
| 311 | 5.02 | 0.19 | 5.64 | 0.56 | 6.70 | 0.39 | 6.64 | 0.26 | 6.90 | 0.24 |
| 313 | 4.12 | 0.26 | 5.00 | 0.25 | 5.67 | 0.29 | 5.65 | 0.33 | 6.05 | 0.33 |
| 314 | 4.41 | 0.16 | 4.78 | 0.44 | 5.95 | 0.08 | 5.83 | 0.17 | 6.05 | 0.20 |
| 317 | 4.58 | 0.21 | 4.90 | 0.45 | 5.82 | 0.23 | 5.62 | 0.12 | 5.95 | 0.21 |
| 318 | 3.85 | 0.19 | 4.31 | 0.30 | 5.46 | 0.11 | 5.20 | 0.15 | 5.72 | 0.25 |
| 319 | 3.49 | 0.08 | 3.68 | 0.06 | 5.08 | 0.22 | 4.48 | 0.11 | 4.69 | 0.12 |
| 322 | 3.70 | 0.14 | 4.62 | 0.34 | 5.53 | 0.10 | 4.76 | 0.27 | 4.92 | 0.13 |
| 323 | 4.68 | 0.13 | 5.60 | 0.31 | 5.70 | 0.19 | 5.44 | 0.29 | 6.14 | 0.39 |
| 324 | 3.41 | 0.15 | 4.00 | 0.19 | 5.19 | 0.23 | 4.55 | 0.23 | 4.47 | 0.15 |
| 326 | 4.49 | 0.10 | 4.68 | 1.00 | 5.58 | 0.23 | 5.87 | 0.17 | 5.85 | 0.13 |
| 327 | 4.43 | 0.08 | 4.94 | 0.38 | 5.93 | 0.06 | 6.14 | 0.11 | 6.52 | 0.12 |
| 329 | 4.82 | 0.22 | 5.44 | 0.56 | 6.23 | 0.34 | 6.24 | 0.22 | 6.53 | 0.16 |
| 330 | 5.32 | 0.16 | 5.86 | 0.21 | 6.86 | 0.15 | 6.97 | 0.19 | 7.62 | 0.19 |
| 331 | 4.04 | 0.07 | 4.30 | 0.14 | 5.58 | 0.16 | 5.31 | 0.17 | 5.70 | 0.24 |
| 332 | 4.06 | 0.26 | 3.95 | 0.32 | 4.71 | 0.16 | 4.29 | 0.21 | 4.24 | 0.22 |
| 335 | 4.19 | 0.12 | 5.16 | 0.21 | 5.90 | 0.44 | 6.57 | 0.36 | 6.48 | 0.61 |
| 338 | 3.76 | 0.16 | 5.10 | 0.32 | 5.11 | 0.31 | 5.57 | 0.10 | 5.32 | 0.21 |
| 340 | 5.06 | 0.32 | 5.99 | 0.40 | 6.35 | 0.11 | 6.29 | 0.08 | 6.62 | 0.25 |
| 341 | 3.43 | 0.20 | 4.16 | 0.11 | 5.15 | 0.15 | 5.18 | 0.25 | 4.75 | 0.16 |
| 346 | 4.51 | 0.17 | 4.71 | 0.35 | 5.92 | 0.19 | 5.89 | 0.21 | 6.24 | 0.26 |
| 347 | 4.69 | 0.42 | 4.80 | 0.41 | 5.55 | 0.47 | 6.43 | 0.28 | 6.23 | 0.31 |
| 349 | 4.48 | 0.17 | 4.83 | 0.31 | 5.84 | 0.10 | 5.55 | 0.23 | 6.45 | 0.10 |
| 350 | 4.08 | 0.18 | 4.75 | 0.35 | 6.34 | 0.14 | 5.74 | 0.45 | 6.19 | 0.16 |
| 351 | 4.47 | 0.45 | 5.37 | 0.92 | 6.84 | 0.15 | 6.18 | 0.37 | 6.30 | 0.21 |
| 352 | 4.55 | 0.30 | 5.26 | 0.70 | 6.55 | 0.35 | 6.38 | 0.19 | 6.43 | 0.20 |
| 353 | 4.46 | 0.16 | 5.67 | 0.16 | 4.17 | 1.28 | 6.67 | 0.15 | 6.41 | 0.18 |
| 355 | 4.34 | 0.14 | 5.03 | 0.23 | 6.01 | 0.33 | 6.20 | 0.22 | 6.36 | 0.18 |
| 356 | 4.07 | 0.29 | 4.48 | 0.25 | 6.27 | 0.09 | 5.89 | 0.33 | 5.47 | 0.36 |
| 358 | 4.09 | 0.24 | 4.19 | 0.23 | 4.50 | 0.15 | 4.39 | 0.14 | 4.74 | 0.21 |
| 360 | 5.08 | 0.20 | 5.98 | 0.36 | 5.97 | 0.38 | 6.32 | 0.34 | 6.80 | 0.17 |
| 361 | 4.19 | 0.26 | 4.63 | 0.22 | 5.16 | 0.44 | 5.69 | 0.27 | 5.82 | 0.13 |
| 362 | 3.70 | 0.11 | 4.05 | 0.16 | 5.46 | 0.20 | 4.84 | 0.09 | 5.22 | 0.19 |
| 367 | 4.36 | 0.18 | 5.25 | 0.22 | 6.45 | 0.04 | 5.98 | 0.11 | 6.48 | 0.14 |
| 369 | 4.77 | 0.39 | 5.13 | 0.80 | 6.66 | 0.52 | 7.05 | 0.65 | 6.39 | 0.54 |
| 372 | 4.03 | 0.31 | 3.76 | 0.81 | 4.90 | 0.39 | 4.53 | 0.26 | 4.52 | 0.20 |

**Table S9**: *R_2ρ_* relaxation rates measured using HSn pulses (n=1,2,4,6,8) from HARD experiment for FUS-RRM (pH 6.4) with ATP at 600 MHz NMR spectrometer.

| **Residue**  **Number** | ***R_2ρ_* (s^-1^)** | | | | | | | | | |
| --- | --- | --- | --- | --- | --- | --- | --- | --- | --- | --- |
|  | **HS1** | | **HS2** | | **HS4** | | **HS6** | | **HS8** | |
|  | **Value** | **Error** | **Value** | **Error** | **Value** | **Error** | **Value** | **Error** | **Value** | **Error** |
| 285 | 13.69 | 0.38 | 12.95 | 0.39 | 12.37 | 1.91 | 13.19 | 0.44 | 13.07 | 0.23 |
| 287 | 14.17 | 1.08 | 13.62 | 0.67 | 12.66 | 1.49 | 13.91 | 1.87 | 12.63 | 0.52 |
| 288 | 13.92 | 0.68 | 12.46 | 0.34 | 11.77 | 1.75 | 11.39 | 0.58 | 13.58 | 0.75 |
| 289 | 16.73 | 0.80 | 19.53 | 1.53 | 19.09 | 2.58 | 14.38 | 1.10 | 16.63 | 0.77 |
| 290 | 13.14 | 0.28 | 12.75 | 0.18 | 12.16 | 2.33 | 12.04 | 0.37 | 12.58 | 0.12 |
| 294 | 16.29 | 0.29 | 15.87 | 0.36 | 16.15 | 2.16 | 14.75 | 1.14 | 15.86 | 0.33 |
| 295 | 11.22 | 0.32 | 11.03 | 0.29 | 11.09 | 0.72 | 11.03 | 0.59 | 10.76 | 0.64 |
| 296 | 10.10 | 0.64 | 9.68 | 0.20 | 9.00 | 0.84 | 9.88 | 0.65 | 9.68 | 0.61 |
| 297 | 12.23 | 0.18 | 11.38 | 0.12 | 11.45 | 0.70 | 11.60 | 0.42 | 11.40 | 0.12 |
| 298 | 13.84 | 0.41 | 13.23 | 0.20 | 12.78 | 2.16 | 13.24 | 0.44 | 12.30 | 0.22 |
| 300 | 8.72 | 0.47 | 8.48 | 0.36 | 10.81 | 0.91 | 10.49 | 1.51 | 8.49 | 0.45 |
| 301 | 15.91 | 0.53 | 14.30 | 0.97 | 13.44 | 4.08 | 9.94 | 3.12 | 14.93 | 0.85 |
| 302 | 15.06 | 0.80 | 13.21 | 1.06 | 14.25 | 1.60 | 13.96 | 1.22 | 12.38 | 0.71 |
| 303 | 12.13 | 0.22 | 11.63 | 0.17 | 11.61 | 0.42 | 10.58 | 0.26 | 11.12 | 0.31 |
| 304 | 13.20 | 0.44 | 11.91 | 0.25 | 12.25 | 1.11 | 12.06 | 0.47 | 12.78 | 0.66 |
| 305 | 13.72 | 0.37 | 12.71 | 0.14 | 13.41 | 0.89 | 13.11 | 0.44 | 12.99 | 0.42 |
| 306 | 12.42 | 0.27 | 12.38 | 0.54 | 11.17 | 1.96 | 12.12 | 0.72 | 11.66 | 0.55 |
| 307 | 13.74 | 0.51 | 13.38 | 0.39 | 12.97 | 1.70 | 12.43 | 0.56 | 13.28 | 0.77 |
| 308 | 13.58 | 0.56 | 12.83 | 0.49 | 12.72 | 1.73 | 12.04 | 0.49 | 13.23 | 0.47 |
| 309 | 12.93 | 0.22 | 12.76 | 0.23 | 12.12 | 1.08 | 12.49 | 0.36 | 12.54 | 0.17 |
| 310 | 12.12 | 0.36 | 12.65 | 0.38 | 10.82 | 1.63 | 10.67 | 0.39 | 12.09 | 0.25 |
| 311 | 16.06 | 0.58 | 15.30 | 1.07 | 14.36 | 2.01 | 13.52 | 0.73 | 14.84 | 1.16 |
| 313 | 12.80 | 0.42 | 11.54 | 0.53 | 11.73 | 1.66 | 11.42 | 0.88 | 12.17 | 0.79 |
| 314 | 13.13 | 0.20 | 12.52 | 0.06 | 12.30 | 0.29 | 12.14 | 0.36 | 12.66 | 0.42 |
| 317 | 12.15 | 0.30 | 11.62 | 0.22 | 11.35 | 0.60 | 11.22 | 0.73 | 11.60 | 0.57 |
| 318 | 12.40 | 0.16 | 11.79 | 0.16 | 11.29 | 0.64 | 11.38 | 0.31 | 11.62 | 0.32 |
| 319 | 11.05 | 0.15 | 10.77 | 0.27 | 10.59 | 1.02 | 10.20 | 0.16 | 10.43 | 0.24 |
| 322 | 12.35 | 0.28 | 11.61 | 0.42 | 11.26 | 0.62 | 10.45 | 0.23 | 11.73 | 0.24 |
| 323 | 9.74 | 0.26 | 9.59 | 0.09 | 8.94 | 1.24 | 9.16 | 0.58 | 9.34 | 0.13 |
| 324 | 12.85 | 0.30 | 11.30 | 0.46 | 12.18 | 0.56 | 11.31 | 0.33 | 11.50 | 0.45 |
| 326 | 12.67 | 0.49 | 12.74 | 0.59 | 12.59 | 0.48 | 11.71 | 0.43 | 12.43 | 0.54 |
| 327 | 14.65 | 0.39 | 14.00 | 0.33 | 13.57 | 0.32 | 13.14 | 0.14 | 13.39 | 0.20 |
| 329 | 12.44 | 0.19 | 12.05 | 0.25 | 11.61 | 0.74 | 11.14 | 0.41 | 11.68 | 0.46 |
| 330 | 19.20 | 0.67 | 17.19 | 0.44 | 18.31 | 1.88 | 16.66 | 1.04 | 18.21 | 1.00 |
| 331 | 11.57 | 0.51 | 11.06 | 0.44 | 10.64 | 0.17 | 10.47 | 0.57 | 11.18 | 0.56 |
| 332 | 5.23 | 0.21 | 5.20 | 0.34 | 4.97 | 0.46 | 4.94 | 0.39 | 5.20 | 0.49 |
| 335 | 16.38 | 0.17 | 14.38 | 0.50 | 11.77 | 2.90 | 13.58 | 1.07 | 15.90 | 0.87 |
| 338 | 13.20 | 0.32 | 13.04 | 0.36 | 11.93 | 1.63 | 12.15 | 0.29 | 12.75 | 0.74 |
| 340 | 13.21 | 0.27 | 13.92 | 0.52 | 11.78 | 0.48 | 12.06 | 1.02 | 13.05 | 0.44 |
| 341 | 12.59 | 0.46 | 12.73 | 0.58 | 11.85 | 0.59 | 12.49 | 0.49 | 12.34 | 0.69 |
| 346 | 13.12 | 0.24 | 12.37 | 0.12 | 12.04 | 0.81 | 11.93 | 0.29 | 12.32 | 0.15 |
| 347 | 14.93 | 1.10 | 14.54 | 1.04 | 14.84 | 2.78 | 13.36 | 1.33 | 14.83 | 0.84 |
| 349 | 12.73 | 0.63 | 11.85 | 0.30 | 12.38 | 0.82 | 12.88 | 1.56 | 11.97 | 0.57 |
| 350 | 13.55 | 0.39 | 12.91 | 0.52 | 12.15 | 0.53 | 11.90 | 0.60 | 12.13 | 0.45 |
| 351 | 15.12 | 1.30 | 14.15 | 0.85 | 14.81 | 1.15 | 13.78 | 0.79 | 13.78 | 0.41 |
| 352 | 14.90 | 0.21 | 14.49 | 0.26 | 14.31 | 0.42 | 13.43 | 0.32 | 13.62 | 0.20 |
| 353 | 13.91 | 0.30 | 9.14 | 1.73 | 12.88 | 1.61 | 11.51 | 2.01 | 12.97 | 0.37 |
| 355 | 14.13 | 0.45 | 13.44 | 0.44 | 12.67 | 1.03 | 12.69 | 0.45 | 12.73 | 0.28 |
| 356 | 15.73 | 0.66 | 14.92 | 0.57 | 15.67 | 1.83 | 15.00 | 1.33 | 14.90 | 0.93 |
| 358 | 8.72 | 0.47 | 8.48 | 0.36 | 8.17 | 1.08 | 8.29 | 0.62 | 8.49 | 0.45 |
| 360 | 15.59 | 0.29 | 14.76 | 0.41 | 14.09 | 1.14 | 13.39 | 0.54 | 13.99 | 0.31 |
| 361 | 14.42 | 0.50 | 13.59 | 0.47 | 13.97 | 2.63 | 13.58 | 1.01 | 14.36 | 0.89 |
| 362 | 10.87 | 0.11 | 10.60 | 0.14 | 10.39 | 0.56 | 10.66 | 0.14 | 10.74 | 0.19 |
| 367 | 13.12 | 0.26 | 12.74 | 0.31 | 12.76 | 0.80 | 12.23 | 0.59 | 12.97 | 0.30 |
| 369 | 12.16 | 0.73 | 11.59 | 0.68 | 10.52 | 0.74 | 11.39 | 1.37 | 10.58 | 0.45 |
| 372 | 7.96 | 0.27 | 8.00 | 0.25 | 7.66 | 0.36 | 7.89 | 0.18 | 7.72 | 0.18 |

**Table S10**: *R_1_* relaxation rates measured from HARD experiment for FUS-RRM (pH 4.6) with ATP at 600 MHz NMR spectrometer.

| **Residue number** | **R_1_ (s^-1^)** | |
| --- | --- | --- |
|  | **Value** | **Error** |
| 283 | 1.53 | 0.02 |
| 285 | 1.43 | 0.05 |
| 286 | 1.64 | 0.09 |
| 287 | 1.87 | 0.10 |
| 288 | 1.62 | 0.13 |
| 290 | 1.45 | 0.08 |
| 291 | 1.55 | 0.10 |
| 293 | 1.51 | 0.07 |
| 294 | 1.32 | 0.04 |
| 295 | 1.42 | 0.03 |
| 296 | 1.25 | 0.10 |
| 297 | 1.52 | 0.07 |
| 298 | 1.52 | 0.07 |
| 299 | 1.40 | 0.05 |
| 300 | 1.54 | 0.04 |
| 302 | 1.44 | 0.04 |
| 303 | 1.54 | 0.08 |
| 304 | 1.67 | 0.06 |
| 305 | 1.73 | 0.06 |
| 306 | 1.51 | 0.04 |
| 307 | 1.53 | 0.03 |
| 308 | 1.72 | 0.09 |
| 309 | 1.53 | 0.05 |
| 310 | 1.21 | 0.12 |
| 311 | 1.65 | 0.18 |
| 313 | 1.41 | 0.09 |
| 317 | 1.42 | 0.02 |
| 318 | 1.39 | 0.02 |
| 319 | 1.59 | 0.04 |
| 322 | 1.59 | 0.11 |
| 323 | 1.53 | 0.04 |
| 324 | 1.32 | 0.07 |
| 326 | 1.42 | 0.03 |
| 327 | 1.46 | 0.02 |
| 329 | 1.37 | 0.02 |
| 331 | 1.45 | 0.03 |
| 332 | 1.55 | 0.02 |
| 333 | 1.47 | 0.05 |
| 338 | 1.60 | 0.10 |
| 339 | 1.49 | 0.20 |
| 340 | 1.55 | 0.04 |
| 341 | 1.45 | 0.11 |
| 343 | 1.58 | 0.03 |
| 346 | 1.49 | 0.07 |
| 350 | 1.53 | 0.11 |
| 351 | 1.63 | 0.08 |
| 352 | 1.49 | 0.04 |
| 353 | 1.42 | 0.10 |
| 354 | 1.58 | 0.08 |
| 355 | 1.54 | 0.02 |
| 356 | 0.99 | 0.06 |
| 358 | 1.55 | 0.03 |
| 359 | 1.50 | 0.06 |
| 360 | 1.50 | 0.04 |
| 361 | 1.42 | 0.06 |
| 362 | 1.48 | 0.02 |
| 367 | 1.56 | 0.02 |
| 369 | 1.45 | 0.05 |
| 372 | 1.67 | 0.09 |
| 374 | 1.60 | 0.01 |

**Table S11**: *R_1ρ_* relaxation rates measured using HSn pulses (n=1,2,4,6,8) from HARD experiment for FUS-RRM (pH 4.6) with ATP at 600 MHz NMR spectrometer.

| **Residue**  **Number** | ***R_1ρ_* (s^-1^)** | | | | | | | | | |
| --- | --- | --- | --- | --- | --- | --- | --- | --- | --- | --- |
|  | **HS1** | | **HS2** | | **HS4** | | **HS6** | | **HS8** | |
|  | **Value** | **Error** | **Value** | **Error** | **Value** | **Error** | **Value** | **Error** | **Value** | **Error** |
| 283 | 3.20 | 0.13 | 3.71 | 0.14 | 4.23 | 0.17 | 3.86 | 0.09 | 4.63 | 0.17 |
| 285 | 3.97 | 0.25 | 5.02 | 0.22 | 5.57 | 0.24 | 5.56 | 0.27 | 5.76 | 0.25 |
| 286 | 4.14 | 0.69 | 4.55 | 0.62 | 5.03 | 0.52 | 5.18 | 0.86 | 8.75 | 2.16 |
| 287 | 4.40 | 0.53 | 5.16 | 0.56 | 6.35 | 1.08 | 5.49 | 0.38 | 6.16 | 0.49 |
| 288 | 4.47 | 0.64 | 4.82 | 0.20 | 5.87 | 0.39 | 5.82 | 0.39 | 6.25 | 0.35 |
| 290 | 4.02 | 0.38 | 3.83 | 0.36 | 3.41 | 0.34 | 4.00 | 0.47 | 5.91 | 0.46 |
| 291 | 3.29 | 0.18 | 4.87 | 0.08 | 4.90 | 0.50 | 5.67 | 0.38 | 5.64 | 0.20 |
| 293 | 4.40 | 0.31 | 5.74 | 0.48 | 6.28 | 0.30 | 6.86 | 0.26 | 6.98 | 0.48 |
| 294 | 5.12 | 0.19 | 6.59 | 0.39 | 8.13 | 0.26 | 8.34 | 0.32 | 9.53 | 0.41 |
| 295 | 3.08 | 0.18 | 5.01 | 0.22 | 5.97 | 0.47 | 5.46 | 0.37 | 4.98 | 0.26 |
| 296 | 2.88 | 0.39 | 5.20 | 0.25 | 6.25 | 0.71 | 4.82 | 0.38 | 5.19 | 0.93 |
| 297 | 4.44 | 0.24 | 5.48 | 0.35 | 6.53 | 0.48 | 5.95 | 0.26 | 6.88 | 0.44 |
| 298 | 4.27 | 0.32 | 4.85 | 0.28 | 5.86 | 0.59 | 5.43 | 0.46 | 6.65 | 0.42 |
| 299 | 2.89 | 0.11 | 3.56 | 0.17 | 3.15 | 0.13 | 2.90 | 0.13 | 3.36 | 0.12 |
| 300 | 4.63 | 0.26 | 5.31 | 0.25 | 6.37 | 0.64 | 5.89 | 0.36 | 6.93 | 0.46 |
| 302 | 4.60 | 0.37 | 5.24 | 0.38 | 6.26 | 0.51 | 5.74 | 0.45 | 7.51 | 0.29 |
| 303 | 4.95 | 0.37 | 5.42 | 0.28 | 6.23 | 0.45 | 6.13 | 0.20 | 7.42 | 0.62 |
| 304 | 4.48 | 0.18 | 4.22 | 0.22 | 4.67 | 0.13 | 4.73 | 0.11 | 7.42 | 0.76 |
| 305 | 4.97 | 0.39 | 5.45 | 0.32 | 6.76 | 0.44 | 5.94 | 0.23 | 7.88 | 0.60 |
| 306 | 3.68 | 0.37 | 4.53 | 0.15 | 5.35 | 0.38 | 5.37 | 0.22 | 5.71 | 0.41 |
| 307 | 4.91 | 0.30 | 5.88 | 0.23 | 6.97 | 0.43 | 6.41 | 0.16 | 7.48 | 0.45 |
| 308 | 4.76 | 0.51 | 5.81 | 0.30 | 7.02 | 0.58 | 6.47 | 0.42 | 7.79 | 0.65 |
| 309 | 4.79 | 0.31 | 5.78 | 0.35 | 6.57 | 0.47 | 6.27 | 0.41 | 7.33 | 0.72 |
| 310 | 2.65 | 0.64 | 5.33 | 0.53 | 5.99 | 0.43 | 5.80 | 0.43 | 4.22 | 0.61 |
| 311 | 3.88 | 0.35 | 4.94 | 0.45 | 6.83 | 0.37 | 6.03 | 0.48 | 6.93 | 0.58 |
| 313 | 3.56 | 0.27 | 4.00 | 0.13 | 4.67 | 0.28 | 4.61 | 0.28 | 4.73 | 1.68 |
| 317 | 4.26 | 0.21 | 5.02 | 0.23 | 5.59 | 0.28 | 5.43 | 0.29 | 6.11 | 0.23 |
| 318 | 4.04 | 0.22 | 4.68 | 0.23 | 5.71 | 0.27 | 5.55 | 0.26 | 5.77 | 0.22 |
| 319 | 3.27 | 0.13 | 3.96 | 0.20 | 4.35 | 0.17 | 4.22 | 0.11 | 5.03 | 0.31 |
| 322 | 3.89 | 0.42 | 4.10 | 0.33 | 5.22 | 0.23 | 4.40 | 0.18 | 6.28 | 0.49 |
| 323 | 3.40 | 0.13 | 5.15 | 0.24 | 5.46 | 0.35 | 5.16 | 0.20 | 4.91 | 0.15 |
| 324 | 4.11 | 0.33 | 4.94 | 0.38 | 6.10 | 0.37 | 5.13 | 0.38 | 6.04 | 0.31 |
| 326 | 4.32 | 0.06 | 5.16 | 0.14 | 6.00 | 0.24 | 6.33 | 0.41 | 7.12 | 0.46 |
| 327 | 4.81 | 0.44 | 5.32 | 0.14 | 6.46 | 0.36 | 6.32 | 0.20 | 7.37 | 0.44 |
| 329 | 4.67 | 0.22 | 5.44 | 0.12 | 6.53 | 0.26 | 6.05 | 0.23 | 7.29 | 0.24 |
| 331 | 3.70 | 0.12 | 4.37 | 0.18 | 5.14 | 0.39 | 4.98 | 0.18 | 5.54 | 0.22 |
| 332 | 2.77 | 0.10 | 3.41 | 0.09 | 3.19 | 0.10 | 3.27 | 0.10 | 3.58 | 0.08 |
| 333 | 4.43 | 0.25 | 5.30 | 0.33 | 6.79 | 0.53 | 6.10 | 0.33 | 6.97 | 0.39 |
| 338 | 3.33 | 0.35 | 4.37 | 0.39 | 5.53 | 0.45 | 5.28 | 0.19 | 6.09 | 0.31 |
| 339 | 4.69 | 0.49 | 4.78 | 0.53 | 6.77 | 0.90 | 4.74 | 0.58 | 6.55 | 0.84 |
| 340 | 4.58 | 0.29 | 5.14 | 0.19 | 5.66 | 0.23 | 5.49 | 0.20 | 6.51 | 0.41 |
| 341 | 3.53 | 0.17 | 4.25 | 0.17 | 5.20 | 0.29 | 4.89 | 0.32 | 6.31 | 0.56 |
| 343 | 3.74 | 0.13 | 4.30 | 0.21 | 4.68 | 0.24 | 4.22 | 0.14 | 4.83 | 0.13 |
| 346 | 3.71 | 0.17 | 5.27 | 0.25 | 6.40 | 0.52 | 6.27 | 0.48 | 6.31 | 0.67 |
| 350 | 5.42 | 0.51 | 5.70 | 0.20 | 6.22 | 0.52 | 6.32 | 0.26 | 7.81 | 0.58 |
| 351 | 3.97 | 0.32 | 4.48 | 0.40 | 5.42 | 0.29 | 5.39 | 0.54 | 6.73 | 0.61 |
| 352 | 4.62 | 0.25 | 5.36 | 0.20 | 6.03 | 0.34 | 5.73 | 0.20 | 7.02 | 0.42 |
| 353 | 4.08 | 0.16 | 5.99 | 0.64 | 7.00 | 0.32 | 6.25 | 0.28 | 6.41 | 0.18 |
| 354 | 4.17 | 0.43 | 5.43 | 0.27 | 6.53 | 0.41 | 5.84 | 0.31 | 6.70 | 0.57 |
| 355 | 4.51 | 0.18 | 5.09 | 0.21 | 5.75 | 0.23 | 5.85 | 0.22 | 6.77 | 0.33 |
| 356 | 2.95 | 0.33 | 3.29 | 0.28 | 3.19 | 0.32 | 3.00 | 0.38 | 3.35 | 0.47 |
| 358 | 2.65 | 0.11 | 2.67 | 0.07 | 2.75 | 0.15 | 2.68 | 0.05 | 3.23 | 0.12 |
| 359 | 3.59 | 0.26 | 4.14 | 0.28 | 5.41 | 0.38 | 5.85 | 0.29 | 5.19 | 0.39 |
| 360 | 4.78 | 0.31 | 5.82 | 0.30 | 5.91 | 0.32 | 5.98 | 0.12 | 7.31 | 0.27 |
| 361 | 3.31 | 0.18 | 4.91 | 0.28 | 5.84 | 0.30 | 5.90 | 0.17 | 6.42 | 0.39 |
| 362 | 3.88 | 0.15 | 4.36 | 0.16 | 4.96 | 0.18 | 5.12 | 0.11 | 5.69 | 0.26 |
| 367 | 3.77 | 0.07 | 5.00 | 0.18 | 5.16 | 0.29 | 5.76 | 0.21 | 6.05 | 0.09 |
| 369 | 4.52 | 0.61 | 5.20 | 0.41 | 5.71 | 0.76 | 5.37 | 0.43 | 6.85 | 0.65 |
| 372 | 3.82 | 0.38 | 3.78 | 0.39 | 4.75 | 0.46 | 4.03 | 0.29 | 4.90 | 0.17 |
| 374 | 3.75 | 0.11 | 4.48 | 0.21 | 4.84 | 0.35 | 4.30 | 0.27 | 4.60 | 0.09 |

**Table S12**: *R_2ρ_* relaxation rates measured using HSn pulses (n=1,2,4,6,8) from HARD experiment for FUS-RRM (pH 4.6) with ATP at 600 MHz NMR spectrometer.

| **Residue**  **Number** | ***R_2ρ_* (s^-1^)** | | | | | | | | | |
| --- | --- | --- | --- | --- | --- | --- | --- | --- | --- | --- |
|  | **HS1** | | **HS2** | | **HS4** | | **HS6** | | **HS8** | |
|  | **Value** | **Error** | **Value** | **Error** | **Value** | **Error** | **Value** | **Error** | **Value** | **Error** |
| 283 | 10.85 | 0.33 | 9.61 | 0.29 | 9.65 | 0.23 | 9.03 | 0.29 | 8.26 | 0.43 |
| 285 | 12.23 | 0.61 | 13.31 | 0.51 | 12.27 | 0.74 | 11.25 | 0.58 | 11.93 | 0.69 |
| 286 | 11.48 | 0.34 | 11.00 | 1.16 | 11.17 | 1.44 | 11.52 | 1.74 | 10.91 | 1.60 |
| 287 | 17.12 | 0.97 | 14.19 | 0.96 | 16.22 | 1.19 | 13.81 | 1.00 | 11.91 | 1.13 |
| 288 | 14.50 | 1.49 | 12.27 | 0.33 | 15.75 | 1.49 | 12.19 | 0.58 | 12.31 | 0.54 |
| 290 | 12.19 | 0.70 | 12.93 | 0.98 | 12.45 | 5.04 | 12.35 | 2.75 | 10.89 | 1.50 |
| 291 | 13.05 | 0.38 | 13.84 | 1.06 | 13.70 | 0.71 | 12.13 | 0.49 | 11.80 | 0.76 |
| 293 | 15.91 | 1.27 | 14.68 | 0.76 | 13.78 | 0.36 | 12.38 | 0.67 | 12.28 | 0.82 |
| 294 | 22.40 | 0.69 | 20.17 | 0.74 | 20.36 | 1.08 | 17.86 | 0.99 | 15.98 | 1.22 |
| 295 | 12.02 | 0.44 | 11.59 | 0.47 | 11.19 | 0.34 | 10.56 | 0.19 | 10.48 | 0.62 |
| 296 | 10.12 | 0.70 | 9.14 | 0.51 | 9.14 | 0.56 | 9.65 | 0.24 | 8.31 | 0.80 |
| 297 | 13.74 | 1.11 | 12.84 | 0.59 | 13.64 | 1.28 | 12.31 | 0.61 | 12.34 | 1.08 |
| 298 | 15.05 | 0.75 | 14.29 | 0.48 | 15.08 | 0.69 | 14.19 | 0.96 | 13.20 | 0.58 |
| 299 | 3.87 | 0.23 | 5.67 | 0.09 | 3.75 | 0.35 | 7.09 | 2.46 | 4.79 | 0.04 |
| 300 | 12.29 | 0.76 | 11.62 | 0.70 | 11.72 | 0.66 | 10.85 | 0.59 | 10.46 | 0.49 |
| 302 | 14.87 | 0.85 | 14.32 | 0.84 | 13.49 | 1.70 | 12.36 | 0.80 | 12.88 | 0.67 |
| 303 | 15.11 | 1.05 | 13.87 | 0.36 | 14.77 | 0.99 | 13.02 | 0.27 | 12.68 | 0.19 |
| 304 | 15.12 | 0.45 | 13.92 | 0.21 | 13.88 | 0.33 | 12.82 | 0.55 | 13.16 | 0.57 |
| 305 | 14.77 | 0.69 | 13.89 | 0.49 | 13.57 | 0.53 | 12.48 | 0.62 | 11.65 | 0.45 |
| 306 | 13.26 | 0.20 | 11.97 | 0.42 | 12.46 | 0.82 | 11.13 | 0.41 | 10.86 | 0.48 |
| 307 | 14.33 | 0.46 | 13.79 | 0.48 | 13.01 | 0.76 | 12.53 | 0.66 | 12.47 | 0.34 |
| 308 | 14.69 | 0.68 | 14.05 | 0.61 | 16.18 | 1.61 | 13.06 | 0.25 | 12.16 | 1.29 |
| 309 | 14.11 | 1.10 | 12.81 | 0.82 | 13.00 | 1.17 | 12.05 | 1.07 | 12.13 | 0.76 |
| 310 | 16.19 | 1.99 | 14.79 | 1.52 | 6.27 | 2.24 | 7.80 | 2.72 | 12.67 | 2.04 |
| 311 | 18.70 | 2.40 | 14.99 | 1.12 | 17.42 | 1.38 | 14.76 | 0.76 | 14.77 | 1.28 |
| 313 | 12.38 | 0.66 | 12.27 | 0.52 | 12.02 | 1.23 | 11.33 | 0.57 | 10.63 | 0.77 |
| 317 | 13.03 | 0.57 | 11.61 | 0.35 | 12.12 | 0.95 | 11.35 | 0.63 | 11.08 | 0.73 |
| 318 | 12.11 | 0.99 | 11.04 | 0.64 | 10.92 | 0.75 | 10.85 | 0.53 | 11.03 | 0.59 |
| 319 | 11.74 | 0.35 | 11.36 | 0.19 | 12.01 | 0.39 | 10.54 | 0.40 | 10.53 | 0.37 |
| 322 | 13.55 | 0.57 | 12.70 | 0.60 | 12.90 | 0.57 | 11.25 | 0.43 | 11.01 | 0.38 |
| 323 | 8.64 | 0.58 | 7.79 | 0.20 | 7.83 | 0.63 | 7.33 | 0.40 | 6.99 | 0.59 |
| 324 | 12.23 | 0.37 | 12.02 | 0.76 | 12.65 | 0.91 | 11.15 | 0.19 | 10.93 | 0.70 |
| 326 | 15.25 | 1.14 | 15.18 | 0.72 | 14.05 | 0.83 | 12.74 | 0.90 | 13.57 | 0.94 |
| 327 | 17.41 | 0.41 | 15.56 | 0.36 | 16.22 | 0.79 | 14.58 | 0.56 | 14.40 | 0.86 |
| 329 | 15.85 | 0.25 | 12.50 | 2.84 | 14.87 | 0.24 | 13.73 | 0.42 | 13.66 | 0.50 |
| 331 | 10.65 | 0.89 | 9.02 | 0.85 | 9.96 | 1.20 | 8.71 | 0.63 | 8.36 | 0.43 |
| 332 | 5.84 | 0.23 | 5.59 | 0.13 | 5.93 | 0.19 | 5.50 | 0.10 | 5.34 | 0.13 |
| 333 | 15.59 | 0.66 | 13.45 | 0.35 | 13.45 | 4.57 | 12.33 | 0.79 | 12.39 | 0.18 |
| 338 | 15.49 | 0.58 | 14.94 | 0.66 | 14.24 | 0.49 | 12.86 | 0.82 | 13.36 | 1.08 |
| 339 | 17.20 | 1.41 | 11.37 | 0.62 | 10.42 | 2.33 | 11.20 | 1.00 | 10.51 | 1.26 |
| 340 | 14.46 | 0.75 | 12.56 | 0.51 | 12.57 | 0.83 | 11.69 | 0.27 | 11.92 | 0.29 |
| 341 | 13.86 | 0.42 | 12.45 | 0.81 | 12.34 | 0.60 | 13.09 | 0.61 | 12.15 | 0.65 |
| 343 | 7.28 | 0.39 | 6.90 | 0.20 | 7.04 | 0.52 | 6.69 | 0.31 | 6.57 | 0.24 |
| 346 | 13.43 | 1.06 | 12.06 | 0.56 | 12.31 | 0.95 | 10.89 | 0.94 | 9.70 | 0.80 |
| 350 | 13.48 | 1.31 | 11.97 | 0.85 | 12.65 | 1.21 | 9.43 | 0.40 | 9.47 | 0.72 |
| 351 | 17.92 | 1.88 | 15.13 | 0.63 | 15.58 | 0.70 | 16.18 | 0.58 | 14.64 | 0.95 |
| 352 | 16.48 | 0.67 | 15.33 | 0.38 | 16.19 | 0.51 | 14.46 | 0.30 | 14.82 | 0.26 |
| 353 | 15.79 | 0.94 | 13.86 | 0.32 | 14.40 | 0.34 | 13.86 | 0.82 | 13.12 | 0.62 |
| 354 | 19.71 | 0.93 | 17.90 | 0.99 | 16.53 | 2.02 | 14.90 | 1.33 | 14.54 | 1.49 |
| 355 | 16.14 | 0.27 | 14.65 | 0.30 | 14.94 | 0.32 | 13.86 | 0.19 | 13.91 | 0.52 |
| 356 | 3.11 | 0.44 | 3.11 | 0.28 | 2.96 | 0.52 | 2.43 | 0.39 | 2.71 | 0.39 |
| 358 | 5.93 | 0.29 | 5.61 | 0.22 | 5.68 | 0.33 | 5.18 | 0.27 | 5.27 | 0.33 |
| 359 | 17.09 | 0.50 | 16.69 | 0.27 | 17.04 | 1.68 | 15.02 | 0.97 | 16.01 | 0.87 |
| 360 | 17.13 | 0.64 | 16.77 | 0.45 | 16.20 | 0.66 | 14.71 | 0.59 | 16.17 | 1.02 |
| 361 | 14.94 | 0.41 | 15.34 | 0.52 | 14.28 | 0.38 | 14.80 | 0.40 | 14.82 | 0.63 |
| 362 | 11.77 | 0.40 | 10.76 | 0.29 | 11.13 | 0.25 | 10.32 | 0.12 | 10.64 | 0.29 |
| 367 | 14.35 | 0.58 | 13.47 | 0.23 | 13.39 | 0.45 | 12.51 | 0.33 | 12.37 | 0.26 |
| 369 | 12.86 | 1.02 | 12.41 | 1.62 | 10.38 | 1.30 | 10.47 | 0.70 | 8.79 | 0.81 |
| 372 | 8.22 | 0.69 | 7.96 | 0.40 | 8.12 | 0.59 | 7.82 | 0.30 | 6.88 | 0.30 |
| 374 | 5.91 | 0.51 | 8.38 | 2.26 | 5.51 | 0.47 | 5.09 | 0.41 | 5.14 | 0.35 |
